# High-quality peptide evidence for annotating non-canonical open reading frames as human proteins

**DOI:** 10.1101/2024.09.09.612016

**Authors:** Eric W Deutsch, Leron W Kok, Jonathan M Mudge, Jorge Ruiz-Orera, Ivo Fierro-Monti, Zhi Sun, Jennifer G Abelin, M Mar Alba, Julie L Aspden, Ariel A Bazzini, Elspeth A Bruford, Marie A Brunet, Lorenzo Calviello, Steven A Carr, Anne-Ruxandra Carvunis, Sonia Chothani, Jim Clauwaert, Kellie Dean, Pouya Faridi, Adam Frankish, Norbert Hubner, Nicholas T Ingolia, Michele Magrane, Maria Jesus Martin, Thomas F Martinez, Gerben Menschaert, Uwe Ohler, Sandra Orchard, Owen Rackham, Xavier Roucou, Sarah A Slavoff, Eivind Valen, Aaron Wacholder, Jonathan S Weissman, Wei Wu, Zhi Xie, Jyoti Choudhary, Michal Bassani-Sternberg, Juan Antonio Vizcaíno, Nicola Ternette, Robert L Moritz, John R Prensner, Sebastiaan van Heesch

**Affiliations:** Institute for Systems Biology (ISB), Seattle, WA, 98109, USA; Princess Máxima Center for Pediatric Oncology, Utrecht, 3584 CS, The Netherlands; Oncode Institute, Utrecht, The Netherlands; European Molecular Biology Laboratory, European Bioinformatics Institute (EMBL-EBI), Wellcome Genome Campus, Hinxton, CB10 1SD, UK; Cardiovascular and Metabolic Sciences, Max Delbrück Center for Molecular Medicine in the Helmholtz Association (MDC), Berlin, 13125, Germany; Broad Institute of MIT and Harvard, Cambridge, MA, 02142, USA; Hospital del Mar Research Institute, Barcelona, Spain; Catalan Institute for Research and Advanced Studies (ICREA), Barcelona, Spain; School of Molecular and Cellular Biology, Faculty of Biological Sciences, University of Leeds, Leeds, LS2 9JT, UK; Stowers Institute for Medical Research, Kansas City, MO, 64110, USA; Department of Molecular and Integrative Physiology, University of Kansas Medical Center, Kansas City, KS, 66160, USA; HUGO Gene Nomenclature Committee (HGNC), Department of Haematology, University of Cambridge School of Clinical Medicine, Cambridge, UK; Pediatrics Department, University of Sherbrooke, Sherbrooke, Québec, Canada; Centre de Recherche du Centre hospitalier universitaire de Sherbrooke (CRCHUS), Sherbrooke, Québec, Canada; Human Technopole, Milan, 20157, Italy; Department of Computational and Systems Biology, School of Medicine, University of Pittsburgh, Pittsburgh, PA, 15213, USA; Pittsburgh Center for Evolutionary Biology and Medicine, School of Medicine, University of Pittsburgh, Pittsburgh, PA, 15213, USA; Centre for Computational Biology and Program in Cardiovascular and Metabolic Disorders, Duke-NUS (National University of Singapore) Medical School, Singapore; Department of Pediatrics, Division of Pediatric Hematology/Oncology, University of Michigan Medical School, Ann Arbor, MI, 48109, USA; Department of Biological Chemistry, University of Michigan Medical School, Ann Arbor, MI, 48109, USA; School of Biochemistry and Cell Biology, University College Cork, Cork, Ireland; Centre for Cancer Research, Hudson Institute of Medical Research, Clayton, VIC, Australia; Monash Proteomics & Metabolomics Platform, Department of Medicine, School of Clinical Sciences, Monash University, Clayton, VIC, Australia; Charité-Universitätsmedizin Berlin, Berlin, 10117, Germany; Helmholtz-Institute for Translational AngioCardioScience (HI-TAC) of the Max Delbrück Center for Molecular Medicine in the Helmholtz Association (MDC) at Heidelberg University, Heidelberg, 69117, Germany; DZHK (German Center for Cardiovascular Research), Partner Site Berlin, Berlin, 13347, Germany; Department of Molecular and Cell Biology, Center for Computational Biology, University of California, Berkeley, Berkeley, CA, 94720-3202, USA; Department of Pharmaceutical Sciences, University of California, Irvine, Irvine, CA, 92617, USA; Department of Biological Chemistry, University of California, Irvine, Irvine, CA, 92617, USA; Chao Family Comprehensive Cancer Center, University of California, Irvine, Irvine, CA, 92617, USA; Biobix, Lab of Bioinformatics and Computational Genomics, Department of Mathematical Modelling, Statistics and Bioinformatics, Ghent University, Ghent, Belgium; Department of Biology, Humboldt University Berlin, Berlin, 10117, Germany; Berlin Institute of Medical Systems Biology (BIMSB), Max Delbrück Center for Molecular Medicine in the Helmholtz Association, Berlin, 10115, Germany; University of Southampton, Southampton, UK; Department of Biochemistry and Functional Genomics, Université de Sherbrooke, Sherbrooke, Québec, Canada; Department of Chemistry, Yale University, New Haven, CT, 06520, USA; Department of Molecular Biophysics and Biochemistry, Yale University, New Haven, CT, 06520, USA; Institute for Biomolecular Design and Discovery, Yale University, West Haven, CT, 06516, USA; Department of Biosciences, University of Oslo, Oslo, Norway; Whitehead Institute for Biomedical Research, Cambridge, MA, 02142, USA; Department of Biology, Massachusetts Institute of Technology, Cambridge, MA, 02142, USA; Howard Hughes Medical Institute, Massachusetts Institute of Technology, Cambridge, MA, 02138, USA; David H. Koch Institute for Integrative Cancer Research, Massachusetts Institute of Technology, Cambridge, MA, 02139, USA; Singapore Immunology Network (SIgN), Agency for Science, Technology and Research (A*STAR), Singapore; Department of Pharmacy & Pharmaceutical sciences, National University of Singapore (NUS), Singapore; State Key Laboratory of Ophthalmology, Zhongshan Ophthalmic Center, Sun Yat-sen University, Guangzhou, China; Functional Proteomics Group, Institute of Cancer Research, Chester Betty Labs, London, SW3 6JB, UK; Ludwig Institute for Cancer Research, University of Lausanne, Lausanne, 1005, Switzerland; Department of Oncology, Centre hospitalier universitaire vaudois (CHUV), Lausanne, 1005, Switzerland; Agora Cancer Research Centre, Lausanne, 1011, Switzerland; School of Life Sciences, Division Cell Signalling and Immunology, University of Dundee, Dundee, DD1 5EH, UK; Centre for Immuno-Oncology, University of Oxford, Oxford, OX37DQ, UK

**Author notes:** **Address correspondence to:** Sebastiaan van Heesch, PhD, Princess Máxima Center for Pediatric Oncology Heidelberglaan 25, 3584 CS Utrecht The Netherlands; John R. Prensner, MD, PhD, Department of Pediatrics and Biological Chemistry, University of Michigan Medical Science Research Building II, Room 2560B, 1150 Medical Center Drive Ann Arbor, MI, 48109; Robert L. Moritz, PhD Institute for Systems Biology 401 Terry Ave N, Seattle, WA, 98109. Co-first authors (Eric W Deutsch, Leron W Kok, Jonathan M Mudge). Co-senior / corresponding authors (Robert L Moritz, John R Prensner, Sebastiaan van Heesch).

**Keywords:** GENCODE, Ribo-seq, Human Proteome Project, mass spectrometry, immunopeptidomics, proteomics, microproteins, non-canonical ORFs, translation

## Abstract

A major scientific drive is to characterize the protein-coding genome as it provides the primary basis for the study of human health. But the fundamental question remains: what has been missed in prior genomic analyses? Over the past decade, the translation of non-canonical open reading frames (ncORFs) has been observed across human cell types and disease states, with major implications for proteomics, genomics, and clinical science. However, the impact of ncORFs has been limited by the absence of a large-scale understanding of their contribution to the human proteome. Here, we report the collaborative efforts of stakeholders in proteomics, immunopeptidomics, Ribo-seq ORF discovery, and gene annotation, to produce a consensus landscape of protein-level evidence for ncORFs. We show that at least 25% of a set of 7,264 ncORFs give rise to translated gene products, yielding over 3,000 peptides in a pan-proteome analysis encompassing 3.8 billion mass spectra from 95,520 experiments. With these data, we developed an annotation framework for ncORFs and created public tools for researchers through GENCODE and PeptideAtlas. This work will provide a platform to advance ncORF-derived proteins in biomedical discovery and, beyond humans, diverse animals and plants where ncORFs are similarly observed.

## Introduction

The consensus set of protein-coding genes is the foundation upon which biomedical science has developed. While annotation of human proteins began in the 1980s, systematic protein-coding gene annotation was only enabled by the Human Genome Project, and this catalog is still being revised by reference annotation projects such as Ensembl-GENCODE (henceforth GENCODE) and UniProtKB/Swiss-Prot (henceforth UniProt)^1–3^. Thus, determining the set of human protein-coding genes is a dynamic process, involving the discovery of new sequences as well as the removal of those that are reappraised as incorrect.

A tantalizing insight into this process has emerged in the form of translated unannotated human open reading frames (ORFs), which have now been reported widely in human physiology and diseases including cancer, Mendelian disorders, and immunology^4–10^. These translation events have been experimentally detected with ribosome profiling (Ribo-seq), which isolates RNA sequences obtained via ribosome footprinting^11^. Collectively, such ‘non-canonical’ ORFs (ncORFs) may be considered a distinctive category of gene translation, as the vast majority are under 100 codons in size and lack deep evolutionary conservation^12,13^. Most ncORFs are located within presumed long noncoding RNAs (lncRNAs) or untranslated regions (UTRs) of mRNAs. While many important questions about their functionality remain, there is particular excitement about their potential to express microproteins. Certainly, the promise of ncORFs and microproteins to advance medical science is becoming increasingly clear: their investigation informs expanded etiologies for the genetic basis of disease^14–16^, mechanisms of cancer biology^17^, and novel targets for immunotherapy^18,19^. Moreover, a variety of studies have proffered large catalogs of prospective microproteins in cancer through immunopeptidomics modalities^4,20–24^. Nonetheless, while individual research groups have searched Ribo-seq and proteomics data for the presence of ncORF and their products^25–28^, reference annotation catalogs such as GENCODE and UniProt have thus far annotated few of these as canonical proteins. These projects are not primarily disease-focused and have conferred this status thus far only onto translated ORFs for which clear evidence of physiological or cellular function is available. Furthermore, immunopeptidomics data are not yet used as a data source in gene annotation.

We formed a global consortium focused on ncORF gene annotations in 2022, with the initial aim to produce a standardized catalog of translated ncORFs discovered via Ribo-seq^29^, but at that time we did not focus on the questions of protein expression or functionality. Here, we now expand our international collaboration to incorporate proteomics experts, working directly with GENCODE^1^, the Ribo-seq ORF Consortium^29^, PeptideAtlas^30,31^, and the Human Proteome Organization-Human Proteome Project (HUPO-HPP)^32–34^, including leadership of the HUPO-Human ImmunoPeptidome Project (HUPO-HIPP)^35^ (**Figure 1a**). Our goal is to further refine this ncORF catalog with reference annotation-quality peptide spectrum matches (PSMs) observed in data-dependent acquisition (DDA) tandem mass spectrometry (MS/MS) data, and to make this information available in the PeptideAtlas resource for proteomics data.

**Figure 1.**
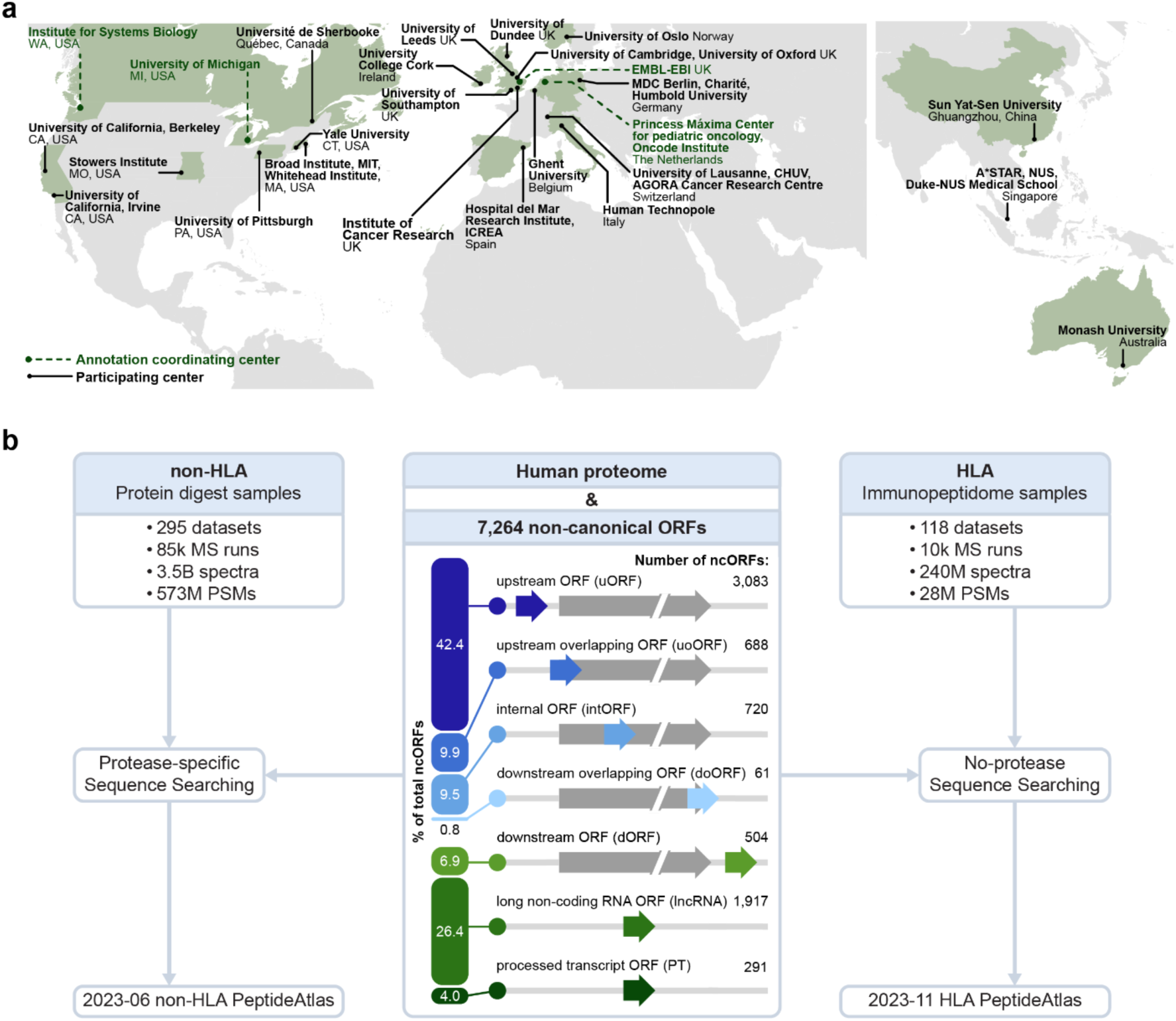
Overviews of the centers participating in the annotation effort and the PeptideAtlas framework for protease-digested (mostly trypsin) sample MS and immunopeptidomics builds. (**a**) Map showing the participating institutions included in the annotation effort. Coordinating centers are highlighted. (**b**) Schematic overview of the datasets included in the non-HLA and HLA builds. The biotypes of the 7,264 ncORFs are shown in the middle.

Using 413 datasets comprising 95,520 MS runs and 3.8 billion tandem mass spectra, we find evidence that at least 25% of 7,264 Ribo-seq ORFs give rise to translation products, primarily detected through HLA immunopeptidomics as opposed to protease-digested samples (usually with trypsin) for traditional MS proteomics. To improve stringency, we manually validate supporting spectral matches, and we use these efforts to systematically classify ncORFs according to a tier system of evidence that integrates manually inspected proteomics and Ribo-seq data. We demonstrate the ability for a multi-Consortia group of experts to prioritize ncORF candidates for potential annotation as protein-coding genes. We additionally describe patterns of ncORF amino acid composition and specific ncORF features that can contribute to increased chances of ncORF immunopeptide presentation, which may inform the targeting of ncORFs in cancer or autoimmune disease through cellular immunotherapies or vaccines. Lastly, we propose a research agenda for the field based on consensus among the multi-consortium group, intended to guide future efforts to bring ncORFs from research discoveries to biological, societal, and biomedical impact via ongoing standardized annotation.

## Results

### Integrating non-canonical ORFs into protein annotation workflows

We sought to provide reference annotation-quality proteomics evidence to identify ncORFs that are translated into human proteins. To do this, we expanded the purview of the PeptideAtlas platform, which is the basis for certification of human protein-coding genes via HUPO and the HPP^32–34^. With 295 protease-digested mass spectrometry (MS) proteomics datasets comprising 3.5 billion MS/MS spectra and 118 HLA immunopeptide-enriched datasets comprising 240 million MS/MS spectra that are publicly available in ProteomeXchange data repositories^36^, we created the Human non-HLA PeptideAtlas 2023-06 and Human HLA PeptideAtlas 2023-11 builds (**Figure 1b**). We built these using a search space that contains the comprehensive THISP (Tiered Human Integrated Search Proteome) level 4 database^37^ plus 7,264 non-canonical ORFs detected by Ribo-seq and supported by GENCODE^29^ (see **Methods**).

We collected and reprocessed these using the Trans-Proteomic Pipeline (TPP) MS data analysis suite^38,39^, wherein raw MS data for each build are annually searched against a comprehensive search database (see **Methods**)^2^. The existence of canonical and non-canonical human proteins is verified with the application of a stringent decoy-estimated false-discovery rate (FDR) of <0.1% at the protein level plus adherence to peptide quality and coverage guidelines set forth in the HUPO-HPP Mass Spectrometry Data Interpretation Guidelines 3.0^40^. This approach led to a peptide-level FDR of 0.0009% for the non-HLA build and 0.0041% for the HLA build (see **Methods**), which is substantially more conservative than many studies due to the requirement that annotation-level proteomics evidence requires higher stringency. Historically, the total number of peptides mapping to canonical proteins has continued to increase steadily as we continued to add datasets to each new PeptideAtlas build. However, progress in the number of validated canonical proteins in the non-HLA build is now very slow, at an average of ∼1 newly verified protein per million PSMs added to the build, computed over the last 100 million PSMs (**Supplementary Figure S1a-d**).

### Bottom-up MS support for non-canonical ORFs: the Human non-HLA build

We first explored proteomics support for 7,264 GENCODE ncORFs using peptide data from conventional enzymatic digests (96.3% of experiments are digested with trypsin, see **Supplementary Table S1**). HUPO-HPP guidelines for human protein verification require two distinct uniquely mapping peptides of length 9 or more residues and minimum protein coverage of 18 residues^40^. We found 484 peptides passing FDR thresholds that map to 183 of the 7,264 ncORFs (∼2.5%) (**Figure 2a-b, Supplementary Table S2**), with 37 ncORFs appearing to have enough evidence to satisfy these requirements (**Supplementary Table S3**). By contrast, 83.0% (16,888/20,359) of the human canonical proteins achieved this level of evidence.

**Figure 2.**
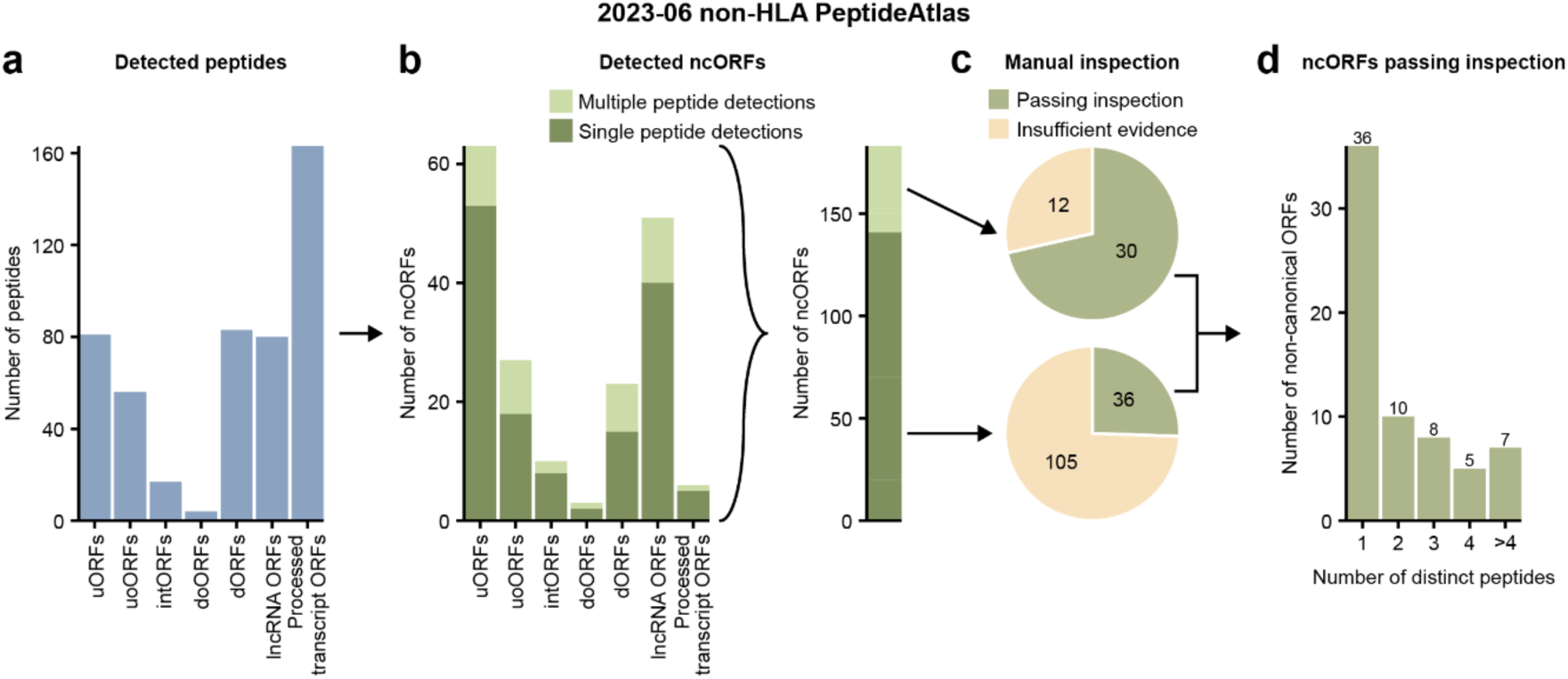
Overview of the 2023-06 non-HLA PeptideAtlas analysis. (**a**) Number of detected peptides in the non-HLA data categorized per ncORF biotype. (**b**) The left graph displays the number of detected ncORFs categorized per ncORF biotype. Bars are shaded by whether an ncORF was detected by a single or multiple peptides. The right bar shows the total number of ncORFs, shaded similar to the bars on the left. (**c**) Pie chart displaying the number of ncORFs that pass after manual inspection of the peptides. The upper pie chart shows the inspection results of the 42 ncORFs detected by multiple peptides. The bottom pie chart shows the inspection results of the 141 ORFs detected by a single peptide. (**d**) Bar plot showing the number of ncORFs passing inspection, categorized by the number of peptides by which they were detected.

Because ncORFs are typically much smaller than annotated coding sequences (CDSs), with a median size of ∼30-40 codons^29^, we next asked whether the size of these prospective proteins was a factor in confident detection by proteomics data. Using a high-confidence manually curated set of small GENCODE proteins^41^, we found that only 2 of 36 known proteins under 50 aa (5.6%) satisfy benchmarks for HUPO-HPP verification. Thus, while small proteins have the potential to be supported by tryptic peptides, the likelihood becomes reduced for very small proteins, potentially indicating a bias in the ability to detect them in tryptic proteomics data, which supports prior evidence^42^.

Further, despite our high-stringency approach, even a small percentage of false positives could yield a substantial number of incorrect identifications when searching 3.5 billion spectra. We therefore manually inspected both the MS spectra, as well as Ribo-seq data, for 42 ncORFs with two unique supporting peptides and 141 ncORFs with one supporting peptide (**Figure 2b, Supplementary Tables S2** and **S3**). In total, 30 ncORFs passed inspection criteria with 2 or more peptides, and an additional 36 with only one good peptide (**Figure 2c-d**). We conclude that high-quality evidence for ncORF translation exists in the human proteome, but that only ∼0.9% of ncORFs yield such results. Implications for gene annotation are discussed in a later section.

### Widespread HLA-peptide support for ncORFs: the Human HLA build

Prior studies have shown that more ncORFs are observed in HLA immunopeptidomics datasets compared to conventional proteomics approaches^4,5,8,20,23,43,44^. Thus, we next sought to identify ncORFs in the ∼240 million MS spectra aggregated within the Human HLA PeptideAtlas 2023-11 build, which we reprocessed according to our established data pipeline (see **Methods**). Overall, we found 3,116 peptides mapping to 1,785 out of 7,264 Ribo-seq ncORFs (24.6%) (**Figure 3a-b, Supplementary Figure S2a, Supplementary Tables S4** and **S5**). While 366 ncORFs had >50% amino acid sequence coverage by HLA peptides, ncORFs exhibiting multiple peptides reflected significantly longer coding sequences compared to ncORFs detected by only one peptide (50.0 aa vs. 44.7 aa respectively; p = 2.3×10^-4^; two-sided Wilcoxon test) (**Figure 3c**).

**Figure 3.**
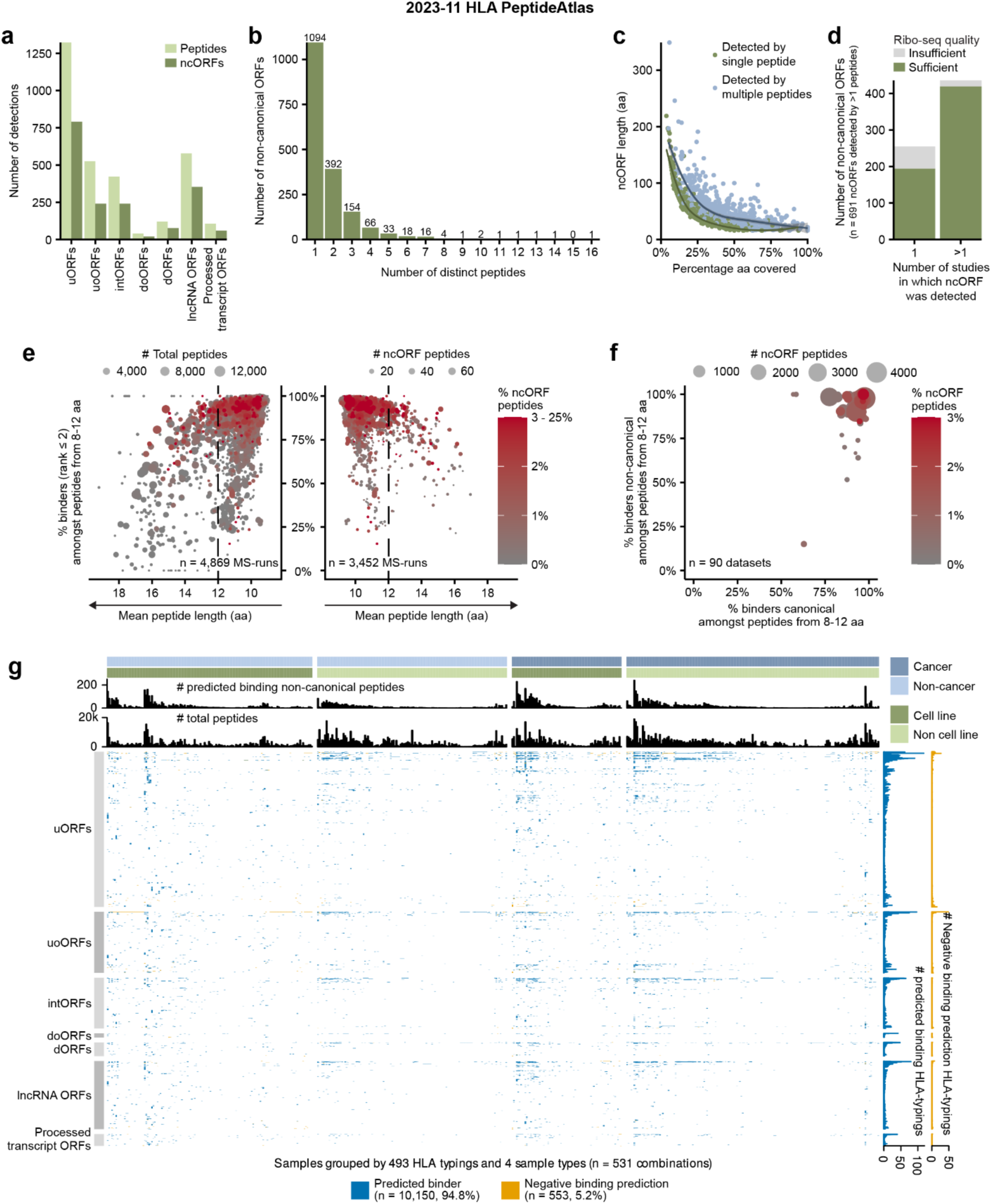
Overview of the 2023-11 HLA PeptideAtlas detected ncORFs. **(a)** The number of distinct peptides and ncORFs detected in the HLA data grouped by ncORF biotype. **(b)** The number of distinct peptides by which an ORF was detected. **(c)** The percentage of the total ncORF sequence covered by HLA peptides plotted against ncORF length. Colors indicate whether a ncORF was detected by one or multiple peptides. Lines were fitted through both groups using Local Polynomial Regression Fitting. Confidence intervals of those lines are shown in gray. **(d)** The number of ncORFs for which the Ribo-seq data quality after manual inspection was judged to be sufficient or insufficient. Only 691 ncORFs detected with two HLA peptides are included. ncORFs are grouped by whether they were detected in a single or multiple studies. **(e)** Dot plots showing the outcomes of the binding affinity predictions. The plots visualize the correlation between mean peptide length and the percentage of predicted binders amongst peptides with a length between 8 and 12 amino acids (NetMHCpan rank ≤ 2) per sample. The left side encompasses all MS-runs, while the right side focuses on samples with at least one ncORF-derived peptide (“ncORF peptide”). Dot size on the left corresponds to the total number of peptides per MS-run, while on the right it corresponds to the count of ncORF-derived peptides. Dot color corresponds with the percentage of ncORF-derived peptides per MS-run. One outlier MS-run (average length 22.75 aa) is not shown. **(f)** Dot plot contrasting the percentage of predicted binders (NetMHCpan rank ≤ 2) per dataset for canonical and ncORF-derived peptides. Dot color corresponds with the percentage of ncORF-derived peptides per dataset. Datasets PXD000171 and PXD022194 are not shown because they have no ncORFs with binding predictions. **(g)** Heatmap indicating whether ncORF peptide detections were verified by NetMHCpan portioned by sample type. HLA typing groups samples based on their associated set of one to six HLA alleles. The upper bar plots display the total number of non-canonical peptides predicted to bind to HLA alleles within a typing and the total distinct peptides associated with it. The right bar plots indicate for each peptide the total count of positive and negative predictions for the HLA typings. Differences in peptide detectability exist across various HLA typings. Overall, peptide detectability concurs with binding predictions.

We observed that virtually all peptides (2,937 out of 3,116; 94.3%) matching ncORFs were found presented by HLA Class-I (HLA-I) alone, with scant evidence for ncORFs in HLA Class-II (HLA-II) datasets (**Figure S2a-f**). This observation contrasts with canonical proteins, presented peptides of which could often also be found in HLA-II data^20,23,44^. This indicates that ncORF peptides are most often sourced from the intracellular pool of protein translation products and less likely from extracellular sources. The lack of observation in HLA-II data could also suggest that ncORF translation products may be unstable and rapidly degraded, making them more likely to be sampled by the HLA-I pathway. The distribution of ncORF-derived peptides across ORF types varied substantially from the baseline prevalence of each type of ORF in the 7,264 Ribo-seq ncORF set (**Supplementary Figure S3a-b**). uoORFs and intORFs showed significant enrichments, and dORFs and lncRNA ORFs showed significant depletions compared to the mean detection rate of 24.6% (p = 7.4×10^-9^; p = 6.3×10^-7^; p = 3.2×10^-6^; p = 1.2×10^-9^ respectively, two-sided Binomial test + Bonferroni correction). Short ncORF length (16-30 aa) led to a 29.3% reduction in the probability of detection as compared to longer ncORFs (>30 aa) (28.1% to 19.9%, **Supplementary Figure S3c-d**). Similarly, we did not observe significant differences in the ability to detect ncORFs between cancer (3,818 MS runs) or non-cancer (5,958 MS runs) sources, and their distribution between cancer or non-cancer samples was not influenced by ncORF peptide mass, hydrophobicity (Kyte-Doolittle), or isoelectric point (**Supplementary Figure S3e-f**).

For these analyses, we estimated an FDR of < 0.1% based on a strategy employing a matched target decoy peptide for each of the 7,264 ncORFs searched (**Supplementary Table S4** and **Methods**). To provide additional rigor, we manually inspected Ribo-seq data for 691 ncORFs with at least two uniquely mapping peptides in the PeptideAtlas HLA build (**Figure 3c**). Overall, 88.7% (613/691) of ncORFs had sufficient Ribo-seq signal to confirm translation at that genomic locus, which could be further parsed into a high-confidence group (96.1% (419/436) verified) and low-confidence group (76.1% (194/255) verified), based on the number of studies in which a ncORF was reported (see **Methods**) (**Figure 3d**). We conclude that the vast majority of peptide identifications are well supported.

### HLA-I binding prediction and allele preferences for ncORF presentation

The PeptideAtlas HLA build provides an opportunity to evaluate the concordance between HLA binding prediction algorithms and actual detections of high-quality PSMs across public HLA immunopeptidomics datasets. Because HLA binding algorithms predict which peptides can bind a specific HLA-I molecule, the concordance between predictions and immunopeptidomics data can be used to further support peptide identification of rare source proteins, such as ncORFs. Therefore, we manually annotated and successfully determined the HLA types of samples used in 4,870 of the 6,479 MS runs (**Supplementary Table S6** and **Methods**).

We next performed *in silico* HLA-I binding predictions with NetMHCpan 4.1^45^ for a subset of 2,711 out of 3,116 ncORF peptides, based on the requirements to have a length of 8-12 amino acids and for being detected within an HLA-I dataset with a known HLA type. For 4,308 out of the 4,870 (88.5%) analyzed HLA-I MS-runs, >70% of detected HLA-I peptides were predicted as binders (percent rank score < 2%) (**Figure 3e**). Higher-quality datasets, as defined by the percentage of strong binders and a mean peptide length smaller than 12 amino acids, yielded proportionally more ncORF peptides than lower-quality datasets (**Figure 3e**). The percentage of ncORF peptides predicted to bind to the annotated HLA-type was high and comparable to the percentage for all canonical peptides in each dataset (**Figure 3f**).

As cells harbor up to six classical HLA-I alleles (two each of HLA-A, HLA-B, and HLA-C), we next used the binding predictions to assign each ncORF peptide to the most likely HLA allele reported or predicted for a dataset. For each detected peptide matching a ncORF, we then checked the individual binding predictions per HLA-typing, ORF biotype, and source material. We observed a strong concordance (94.8%) between predictions and detected peptides across ORF biotypes and independent of the source material (e.g. cancerous or non-malignant cell lines or tissues) (**Figure 3g**). We conclude that the vast majority of ncORF peptides in HLA-I datasets are likely to bind to the HLA molecules expressed by each sample, lending additional confidence to their detection.

### Determinants of ncORF HLA-I peptide presentation and detection

We next asked whether there are key determinants that would make a certain ncORF more likely to be detected in HLA-I immunopeptidomics data. We pursued this question in three ways, through amino acid sequence analyses, DNA sequence analyses, and tissue expression analyses.

First, to evaluate amino acid sequence determinants, we investigated binding predictions, sequence length, mean mass per amino acid, and isoelectric point (IEP) (**Supplementary Figure S4a-b** and **Methods**). We additionally tested statistical learning models to analyze how several amino acid sequence based features influence detectability by MS (**Supplementary Document S2, Supplementary Tables S7** and **S8**), finding that the number of predicted HLA-I binder peptides per unit length, the ORF biotype, and the overall ORF length were the best predictors of whether an ncORF is detected. However, with an area under the curve (AUC) of 0.68, the model is not strongly predictive.

When investigating ncORF characteristics, sequence length and isoelectric point (IEP) were increased in detected ncORFs compared to undetected ncORFs (**Figure 4a**). This appears concordant with recent studies pointing toward sequence determinants impacting the stability of ncORF translation products and/or their detectability using mass spectrometry^46,47^. Interestingly, while IEP was increased in detected ncORFs, detected canonical proteins displayed the opposite pattern (**Figure 4a**), suggesting that the IEP could be linked to HLA presentation selectively for ncORF-derived proteins.

**Figure 4.**
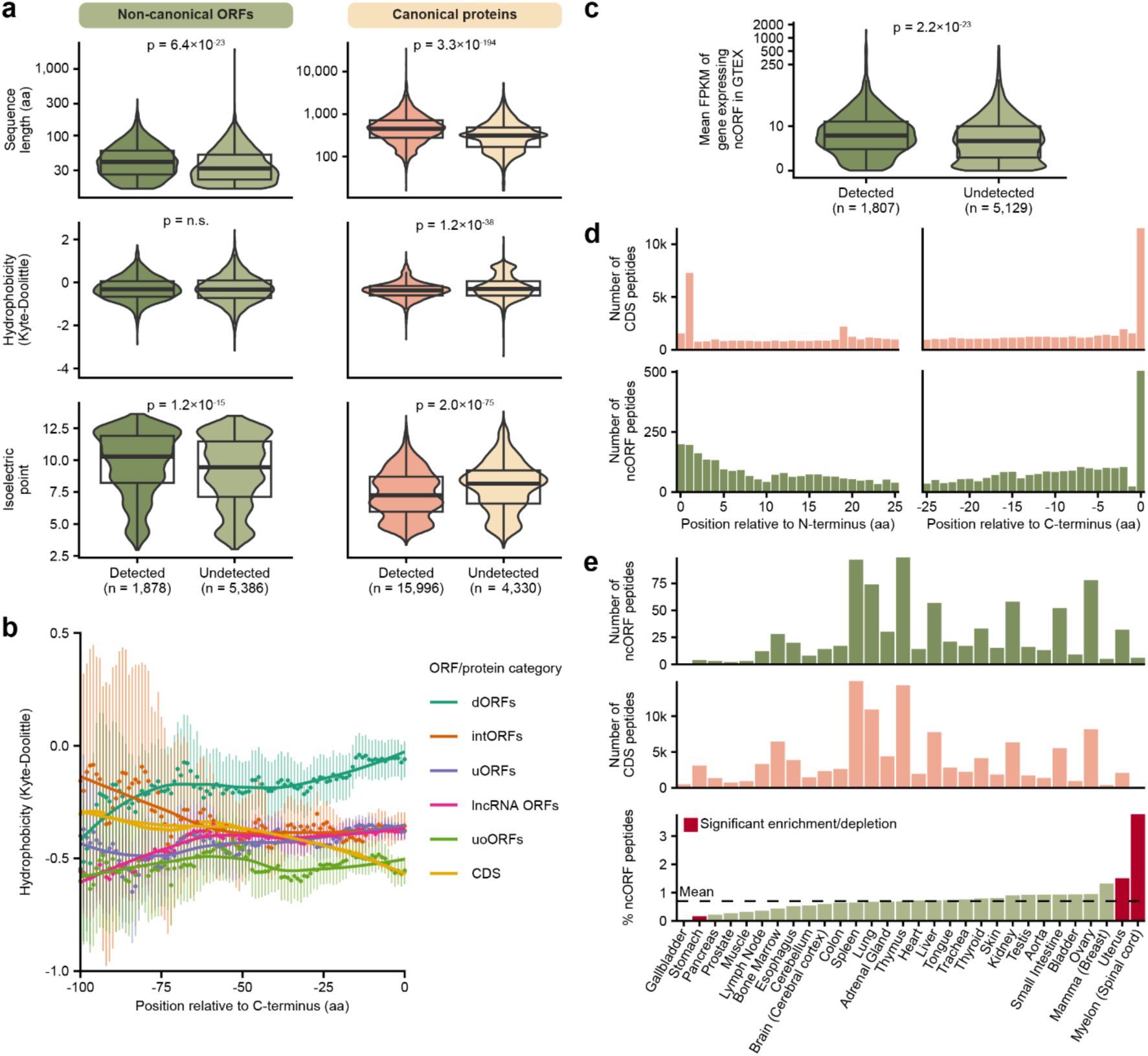
Determinants of ncORF peptide detection. (**a**) Comparison of different sequence properties between detected and undetected ncORFs and canonical proteins (the number of canonical proteins is larger than in (**Supplementary Figure S1d**) because these were selected using less stringent criteria than the PeptideAtlas workflow). The comparisons are based on sequence length, hydrophobicity by the Kyle-Doolittle scale, and the isoelectric point. Statistical tests were performed with the two-sided Wilcoxon test, reported p-values were adjusted for multiple testing with Bonferroni correction. (**b**) Comparison of the hydrophobicity per ncORF biotype. Each dot represents the average hydrophobicity of the amino acids at that position and the 14 amino acids before that position per ncORF biotype or CDS. The lines were fitted using Local Polynomial Regression Fitting. Vertical bars represent 95% confidence intervals. doORFs and processed transcript ORFs are not shown because of their relatively low abundance. Note that because ncORFs are mostly smaller than 100 aa, confidence intervals get larger with increasing C-terminus offset. (**c**) Comparison of the expression levels of detected and undetected ncORFs. On the y-axis, the mean FPKM in GTEX of genes expressing an ncORF is shown on a pseudo-log scale. 326 ncORFs for which the gene id was not present in GTEX are not shown. Significance was determined using the two-sided Wilcoxon test. (**d**) Overview of the location of detected peptides within the full protein (top) and ncORF (bottom) sequence. The left histograms show the distance between the start codon and the start of the detected peptides. The right histograms show the distance between the end of the detected peptides and the last amino acid of the sequence. (**e**) Overview of HLA ligand atlas data grouped by tissue. The top two plots show the number of ncORF peptides and canonical peptides per tissue. The bottom bar graph shows the percentage of ncORF peptides per tissue relative to the total number of ncORF and canonical peptides. Significant differences as determined by Fisher exact tests and Bonferroni correction are colored red. The dashed line shows the mean percentage of ncORFs.

Conversely, we observed no significant difference in sequence hydrophobicity between detected and undetected ncORFs (**Figure 4a**), which contrasts with recent reports^46,47^. When considering C-terminal hydrophobicity, we observed some variability between ncORF biotypes, which may be due to the sequence context for each biotype (**Figure 4b**). However, such ncORF-type specific differences in C-terminal hydrophobicity did not explain the detectability of these ncORFs in the HLA-I data, as detected and undetected ncORFs were equally hydrophobic at their C-terminus (**Supplementary Figure S4c-d**). This suggests that C-terminal hydrophobicity may play a role in how ncORF translation is regulated, but it does not enhance the chances that such products are presented by the HLA-I system.

Next, we investigated whether the detectability of ncORFs correlates with their expression across human tissues^48^. We observed a significant increase in expression for genes encoding detected ncORFs compared to genes encoding undetected ncORFs (14.3 vs. 10.7 respectively; p = 2.2×10^-23^; two-sided Wilcoxon test) (**Figure 4c**). These results are consistent when separated by ORF biotype (**Supplementary Figure S4e**) or by tissue (**Supplementary Figure S4f**), indicating that higher gene expression correlates with the detectability of the ncORF’s peptide product.

To define whether specific parts of ncORFs are preferentially sourced for HLA presentation, we investigated the positional origin of each detected ncORF peptide within the complete ncORF. By determining the N- and C-terminal offset for all detected ncORF and CDS peptides, we observed a preference for peptides originating from the N-terminus or directly at the C-terminus^49^ (**Figure 4d**). N-terminal enrichment was stronger for canonical proteins than for ncORFs, and a strong preference for peptides initiating at the 2nd residue after methionine cleavage was observed (a 4.6-fold vs. 1-fold change between the number of peptides starting at the second and first N-terminal residues, p = 2.4×10^-46^, Fisher’s exact test). C-terminal enrichment was observed for both canonical and ncORFs, but was more significant for ncORFs (20.3-fold vs 7.2-fold, respectively, p = 9.8×10^-9^, Fisher’s exact test). This positional effect could result from a lower number of cleavages required to process peptides from the termini and might be influenced by sequence length and the capacity of the proteasome to digest short sequences.

Second, we assessed whether protein sequence conservation is associated with the detectability of ncORFs in HLA-I data. We used a previously published evolutionary classification of the 7,264 ORFs^12^. For a direct comparison between CDSs and ncORFs, we identified a set of 406 ncORFs and 29 canonical CDSs with sizes below 50 codons classified as being conserved across mammals^12^. Between these two groups, we found a similar percentage were detected in HLA-I data (154/406 (37.9%) ncORFs; 11/29 (37.9%) canonical proteins). By contrast, evolutionarily young ncORFs below 50 codons that emerged during primate evolution showed lower support by HLA-I peptides (1,033/4,798; 21.5%, p = 7.6×10^-13^; Fisher’s exact test for conserved vs. young ncORFs). This observation is not merely a consequence of the shorter length of evolutionarily young ncORFs (mean length of 29 codons) compared to conserved ncORFs (mean length of 37 codons). To control for this variable, we generated a subset of 406 young ncORFs matched in length to conserved ncORFs. The support by HLA-I peptides remained significantly lower in this set of young ncORFs (92/406; 22.6%, p = 3.0×10⁻⁶; Fisher’s exact test, conserved vs. length-corrected young ncORFs). Therefore, evolutionary conservation may correlate with ncORF peptide detectability.

Finally, we looked at the influence of tissue type on ncORF peptide detection. To investigate whether certain tissues display more ncORF peptides than others, we used the HLA-ligand-atlas data^50^, which provides immunopeptidomics data categorized by tissue type (**Supplementary Figure S4g**). Comparing the relative proportions of ncORF peptides and CDS HLA-I peptides per tissue, the proportion of ncORF peptides in stomach tissue showed a subtle decrease compared to the mean percentage of ncORF peptides across all tissues (-0.6%, p = 1.7×10^-4^, Fisher’s exact test) (**Figure 4e**). On the other hand, the spinal cord and uterus showed mild enrichments in ncORF peptides (0.8%, p = 1.9×10^-3^; 3.1%, p = 0.029) (**Figure 4e**). These observations were not explained by differences in RNA transcript expression in these specific tissues (**Supplementary Figure S4h**). Although modest in effect size, these results may point to tissue-specific regulation of ncORF translation and presentation in the immunopeptidome.

### An annotation framework for protein-coding ncORFs

A primary goal of this work is to develop a standardized analytical framework and nomenclature system for assigning peptide evidence to ncORFs^43^. In this system, we utilize proteomics, immunopeptidomics, and Ribo-seq as complementary techniques that guide a tier-based classification of ncORFs (**Figure 5a** and **Methods**). The level of evidence supporting whether a given ncORF is (i) translated and (ii) expressed as a protein, is conveyed to users via an intuitive framework. Furthermore, this resource will now play a core role in the next phase of our project that focuses on functional characterization of ncORFs in reference gene annotation.

**Figure 5.**
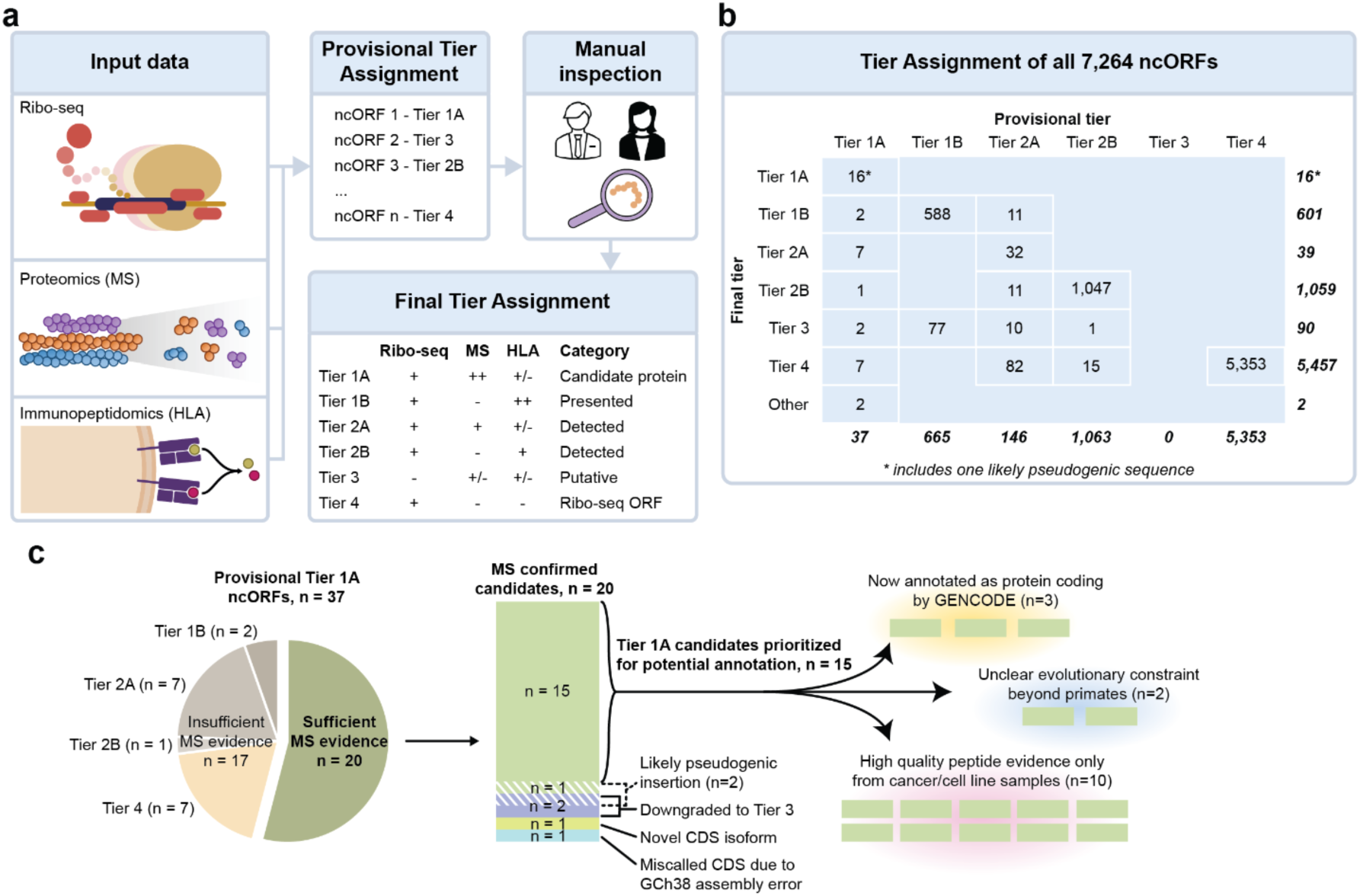
Overview of the Tier system. (**a**) Schematic showing how provisional and final tiers can be assigned to ncORFs. First Ribo-seq, proteomics and immunopeptidomics data can be (computationally) integrated to assign provisional tiers based on the quality of each data entity. Manual inspection of each data entity is then necessary to assign a final tier to each ncORF. In this figure, ‘+’ denotes detection, ‘++’ denotes abundant detection, ‘+/-’ denotes either presence or absence of detection, and ‘-’ denotes absence of detection. (**b**) Results of the provisional and final tier assignment for the 7,264 ncORFs analyzed for this study. (**c**) Overview of the curation process for the provisional Tier 1A ncORFs.

Here, we highlight several key observations in the deployment of our tier system, and preliminary gene annotation changes. First, 37 of 7,264 (0.5%) ncORFs with mass-spectrometry support would provisionally be classified as Tier 1A status, indicating the highest level of experiment support in conventional proteomics and Ribo-seq data. These ncORFs have multiple tryptic proteomic peptides that satisfy HUPO-HPP guidelines for protein verification^40^. However, as listed in **Supplementary Table S3** and described in further detail in **Supplementary Document S1**, inspection of spectra quality reduced this number to 20 candidates for Tier 1A status (**Figure 5b**). Yet, among these, manual inspection of gene models and Ribo-seq signal determined that 2 candidates were most likely pseudogenic sequences, 1 candidate reflected a problem with the GRCh38 reference genome that falsely bisected a CDS into a ‘ncORF’, 1 candidate is better explained as a novel protein isoform of an existing CDS, and 2 candidates had insufficient Ribo-seq evidence for confident assessment. Thus, we conclude that 15 ncORFs represent high-quality Tier 1A candidates for which annotation as a protein-coding gene could be considered (**Figure 5c**). This process illustrates the value of our fully integrated approach in producing reference-quality manual annotation, a workflow that will be maintained as this project progresses. We also note that the reappraisal of ncORFs as previously unannotated alternative proteins isoforms or pseudogenes also leads to productive GENCODE annotation, and that novel isoforms especially may turn out to be functionally important.

GENCODE have so far annotated three of the 15 ncORFs with Tier 1 support as protein-coding genes. One of these is c11riboseqorf4, a 171 aa upstream overlapping ORF (uoORF) of *PIDD1*^51^ (**Figure 6a**). While c11riboseqorf4 does not show protein-level evolutionary constraint^52^, GENCODE have now annotated this uoORF as a new protein-coding gene, ENSG00000293685, based on the strength of the experimental evidence. The ncORF exhibits eight MS peptides that we deem as ‘excellent’ (see **Methods** for evaluation details) and do not map elsewhere in the human proteome, either directly or as allelic variants. Furthermore, these peptides are found in non-malignant tissue samples in addition to cancer samples and cell lines, suggesting a physiological role for the protein (see **Box 1**). Manual inspection of Ribo-seq data for this ORF also shows a clear site of translational initiation supported by translation inhibitors enriching ribosomes at translation initiation sites (**Figure 6a**).

**Figure 6.**
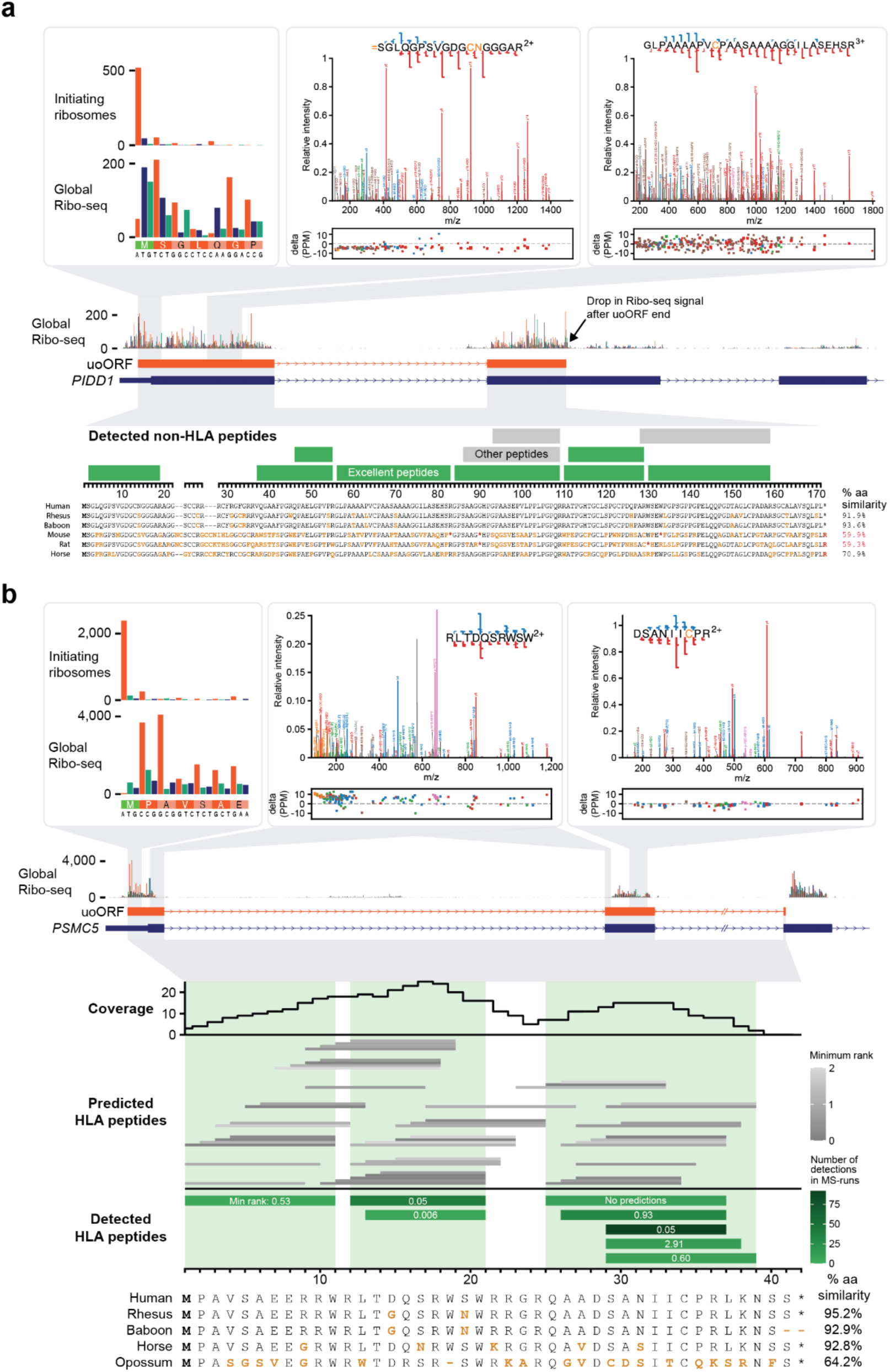
Examples of two ncORFs detected by either non-HLA or HLA data. (**a**) Ribo-seq, mass spectrometry, and evolutionary information for c11riboseqorf4, one of the best detected ncORFs in tryptic digests. This ncORF has 11 distinct peptides across 94 different experiments, 8 of which we classified as excellent evidence (green). The spectra for peptides SGLQGPSVGDGCNGGGAR and GLPAAAAPVCPAASAAAAGGILASEHSR are depicted with nearly complete y ion coverage and substantial b ion coverage, providing highly compelling evidence. We also note that SGLQGPSVGDGCNGGGAR begins as position 2 of the ORF and has peptide N-terminal acetylation, indicating ORF N-terminal acetylation after removal of the initiator methionine. (**b**) Overview of data available for c17norep146, an uoORF in the *PSMC5* gene. Ribo-seq data shows the initiation of translation at the methionine translation initiation codon (green). A-sites are colored by the reading frame (orange for the uoORF, blue for *PSMC5*. Two peptide spectral matches for HLA-I peptides RLTDQSRWSW and DSANIICPR are shown (USIs are mzspec:PXD004894:20141214_QEp7_MiBa_SA_HLA-I-p_MMf_4_2:scan:31976:RLTDQSRWSW/2, mzspec:PXD029567:UPN20_class_I_Rep3:scan:6685:DSANIIC[Cysteinyl]PR/2, respectively). The lowest panel shows the position of all 8 peptides that were observed in the immunopeptidomics data. The color shading indicates the number of MS runs in which each peptide was observed. The middle panel shows all peptides that are predicted with NetMHCpan to be observable in the MS runs (i.e. they are predicted to bind with NetMHCpan score <2 to at least one allele in one of the samples in which peptides were observed). The top part shows the number of predicted binding peptides in which each amino acid was located. Green shadings indicate which part of the ORF sequence was observed. Detected peptides occurred in the regions with the highest numbers of predicted binders.

While Tier 1A candidates are the obvious first candidates for potential new protein-coding genes, our Tier system further prioritizes other ncORFs. Tier 2A candidates harbor one peptide, which may therefore capture possible microproteins that are too short to produce multiple peptides. Close inspection of these candidates was particularly important: of 146 candidates with a nominated peptide, only 32 (26.7%) harbored high-confidence peptide spectra and Ribo-seq signals that passed our manual evaluation criteria. Yet, we identify 21 Tier 2A candidates with a high-confidence tryptic peptide, Ribo-seq data, and additional high-confidence HLA peptides that would previously not be considered as supporting evidence (**Supplementary Table S3** and **Figure 5**). Although these 21 candidates exhibit only 1 peptide in tryptic MS data suitable for potential annotation, they have up to 14 peptides in HLA data (**Supplementary Table S3)**. We have elevated this set of ncORFs for critical examination as potential protein-coding genes.

Elsewhere, this Tier system assists in the stratification of HLA-based evidence. Tiers 1B and 2B reflect high-confidence evidence for certain ncORFs as *presented* HLA ligands. The biological interpretation of abundant HLA peptidomics support for a given ncORF remains a point of active discussion, especially in the absence of conventional proteomics evidence or an evolutionary argument^46,53^. Nonetheless, the confident detection of HLA peptides remains immensely important for researchers, as this method can help identify potential candidates for therapeutic targets, such as autoimmune disease, cancer vaccines or other immuno-oncology approaches^4,19,20,22,24^. Intriguingly, our preliminary gene annotation work highlights a subset of upstream overlapping and internal ORFs (uoORFs and intORF) with exceptional evidence for dual-frame translation of two overlapping coding sequences in HLA data. For example, c17norep146 is a uoORF that overlaps the annotated CDS of *PSMC5* (PSMC5, UniProtKB accession P62195). It displays clear Ribo-seq translation in the correct frame, far exceeding the translation rate of the annotated CDS, and is supported by 8 distinct HLA-I peptides with excellent spectra across 22 HLA-I datasets, covering 85% of this uoORF’s reading frame (**Figure 6b**).

Similarly, c5norep142 - an internal ORF (intORF) within the canonical CDS of *MATR3* (MATR3, UniProtKB accession P43243) - is supported by eight distinct HLA peptides found across 27 HLA-I datasets comprising 270 unique MS runs (**Supplementary Figure S5**). This entire intORF is found in almost all of the 241 evaluated mammalian genomes^54^. However, we do not observe support for protein-level constraint, which makes the nature of its implied function hard to predict. A deeper analysis of the *MATR3* locus implies that c5norep142 translation is mediated by alternative splicing: the exon containing the canonical start of the *MATR3* CDS is commonly spliced out, and the alternative frame intORF is exposed as the first plausible translation in transcripts where this happens.

At this point, while we are confident that both the c17norep146 and c5norep142 ncORFs generate protein products, uncertainties about their physiological nature have led to both ORFs being held back from protein-coding gene annotation.

Finally, to improve the accessibility of these data and visibility of ncORF evidence, we have integrated all ncORFs, peptides, and spectra evaluated as part of this project into PeptideAtlas (https://peptideatlas.org/builds/human/#ncORFs). Users can search for an individual ncORF by name or sequence and retrieve relevant peptide data.

#### Box 1

**setting an agenda to advance the ncORF field**

A unique capacity of our multi-consortium collaboration is the ability to develop consensus on the key challenges that we feel the ncORF field needs to address. Moving forward, we see seven areas where the research community should be engaged:

1. *Are HUPO-HPP guidelines for protein verification suitable for ncORFs?* These require two peptides of length 9 aa or more, and spanning at least 18 aa of the ORF^40^. Yet, many ncORFs are smaller than 18 aa^10,12,29^, and 28.3% (2,059/7,264) of ncORFs in this study are <25 amino acids, making it inherently difficult to meet these guidelines.
2. *Should HLA immunopeptidomics be used as evidence that a ncORF encodes a protein-coding gene?* 1,785 out of 7,264 ncORFs are observed with high-quality HLA data, including 24 ncORFs with only 1 peptide in tryptic MS data suitable for potential annotation but up to 14 peptides in HLA data.
3. *Should peptides detected in cancer samples or immortalised cell lines support protein-coding gene annotation?* 2.36 billion out of 3.53 (66.9%) billion MS2 spectra searched in the non-HLA PeptideAtlas are from cancer tissue or cancer cell line samples. Proteins supported by such data are potentially cancer-specific products, which has implications for gene annotation.
4. *What is the role of evolutionary inference in annotation for ncORFs?* Most of the 7,264 Ribo-seq ncORFs analyzed here are evolutionarily young and lack measurable sequence constraint^12,13^, and so cannot be annotated as protein-coding solely on this basis. What is less clear is the extent to which the lack of observable constraint argues *against* function^55,56^.
5. *Which alternative forms of experimental analysis could be used to support protein-coding gene annotation?* Clearly, it would make sense for any ncORF to be annotated as protein-coding if evidence is provided not only for the existence of the protein, but also the nature of at least one biological function (e.g.,^57,58^). Of note, immune recognition of a peptide is not currently considered a biological function.
6. *How should we annotate cellular proteins for which function can be neither demonstrated nor inferred?* Ideally, a strategy would have support across reference annotation projects as well as community buy-in. It would also be adaptable and able to accommodate rapidly emerging scientific insights.
7. *Should deep learning approaches inform gene or protein annotation?* While annotation is historically rooted in manual inspection, advancements in deep learning may offer an opportunity to classify high-quality mass spectrometry spectra and Ribo-seq data for future annotation efforts^59–62^.

## Discussion

Here, we address the central question of how to systematize efforts to detect ncORFs in proteomics data in a manner that can be aligned with gene and protein annotation projects, pairing experts within the HUPO-HPP/PeptideAtlas project with members of the HIPP immunopeptidomics project, the Ribo-seq ORF Consortium and the GENCODE reference gene annotation group. We describe 1,715 individually-inspected ncORFs as having peptide evidence in addition to Ribo-seq support, which we thus consider as potential proteins. These ncORFs have at least one manually validated peptide in either tryptic or HLA datasets, totaling 3,035 peptides. We present these data at the evidence level in a format that can be used by both researchers and annotation projects, formalized within our Tier classification system. Of note, our efforts emphasize large-scale manual inspection of both peptide data and ribosome profiling data because of the central role manual inspection plays in reference gene annotation efforts. However, we appreciate that manual inspection of thousands of candidates is not feasible for most researchers outside of the gene annotation ecosystem, who may employ tools assessing peptide retention time prediction and ion mobility prediction^59,60,63–65^ to achieve rigorous data evaluations in a scalable way for high-throughput research studies.

As previously observed^4,5,20,22,66,67^, we show that ncORFs are highly abundant in HLA-I immunopeptidomics datasets – wherein upwards of 24.6% of ncORFs may be observed – but rarely detected in conventional proteomics datasets, where ∼0.9% are nominated. There has recently been significant speculation on this point, with growing consensus pointing towards lower stability for many ncORF-derived proteins, perhaps due to BAG6-mediated degradation in the proteasome^23,46^. The proteasome also processes peptides generated through the ribosome quality control machinery, which could further explain the observation of ncORF-derived peptides in HLA-I but not HLA-II data^68^, the latter of which instead samples proteome-derived peptides through an endosomal processing pathway^69^. Regardless of the mechanism of degradation, ncORF-derived protein products, if short-lived, may have greater technical challenges in being identified in standard tryptic mass spectrometry approaches, which has been discussed for annotated proteins with short half-lives^70,71^.

As a community focused on annotation, we advise against over-interpretation of the absence of ncORF-encoded proteins in tryptic mass spectrometry data: fewer than 5.6% (2/36) of previously annotated protein-coding genes < 50 aa meet criteria to verify protein existence in the current PeptideAtlas dataset. Tryptic MS also conventionally biases against proteins with a transmembrane domain^72^, which may be present in some microproteins^73,74^. Thus, the lack of MS peptide data for a given short Ribo-seq ORF does not indicate that it is *not* protein-coding. Instead, these observations support our view that, while proteomics is demonstrably a useful tool for the validation of microproteins, it cannot be the sole adjudicator, perhaps especially for *very* small proteins. We additionally point out that our work has focused on data-dependent acquisition (DDA) for mass spectrometry; data-independent acquisition (DIA) may also be informative for ncORF-derived proteins in specific contexts^75^, and future work will need to establish best-practices guidelines for DIA approaches.

More broadly, this work introduces a new challenge as the concepts of *protein identification* and *protein-coding gene annotation* are distinct. While protein identification refers to the experimental detection of an actual molecule, protein-coding gene annotation is historically rooted in the idea that the translated protein imparts a biological function. Therefore, GENCODE is proceeding with open-minded caution towards the annotation of ncORFs as *protein-coding genes*: ncORFS typically lack the central notion of inferred biological function (e.g. through evolutionary constraint), but are nevertheless robustly detected in many cases. As an important consideration, we note that GENCODE annotates alternative proteins that are found within the same protein-coding locus but do not share any overlapping CDS in the same frame as separate genes. The major reason for this pragmatic: users of annotation commonly wish to work with a single model per protein-coding gene, e.g. via the MANE Select set, and if multiple independent proteins from the same locus were grouped together as a single gene then only one would make it into such analyses.

*How then might the research community reconcile the pragmatic aspects of protein-coding gene annotation with the large-scale detection of ncORF peptides in proteomics data?* This question is particularly acute for immunopeptidomics data, where already thousands of ncORFs are detected, and we expect many more to be found as more comprehensive RiboSeq ORF catalogs with HLA support emerge.

Ultimately, we view proteomics data as a tool to spotlight ncORFs of high potential to be annotated as protein-coding loci. This potential is most tangible for tryptic MS data, which implies intracellular stability of a ncORF-derived protein. Yet, we propose that HLA peptide evidence may also illuminate ncORFs that are *potentially* translated as veritable proteins, noting that it remains to be determined what fraction of HLA-detected ncORFs lack evidence in tryptic MS proteomics for either technological or biological reasons. In annotation terms, we consider such ncORFs to be “on deck” for further evaluation as this work moves forward. Because protein-coding gene annotation remains an essentially manual endeavor, such observations can be valuable for triaging top candidates.

However, our current efforts have fallen short of resolving several important questions (**Box 1**). One central theme that emerges is the ability for detected peptides to inform biological insights about the nature of the ncORF, which is central to the idea that a *protein-coding gene* should be conceived as the producer of a biological actor in a cell. This is a particularly vexing question given that small ORFs typically lack evolutionary signatures that suggest a conventional protein-coding gene, either because small ORFs are truly less conserved or, alternatively, less well captured by conventional tools used for measuring protein sequence constraint^76,77^. Thus, it remains possible that certain ncORF peptides reflect aberrant proteins whose existence is deemed out of context with the canonical proteome. Such ‘aberrancies’ could manifest as translations specific to cancer or autoimmune disease, where diminished ribosome fidelity perhaps produces peptides with no physiological basis in normal biology^78^, ribosome scanning byproducts that are presented by the HLA system^46^, or reactions to cell stress such as amino acid deprivation^79–81^.

For example, we observe that ncORFs in *STK11*, *ZNF219* and *CIRBP* are well supported by tryptic mass spectrometry peptides, and yet these ORFs are not found beyond the primate order. In the case of these three ncORFs, each of the supporting peptides is derived either from cancer samples or immortalized cell lines. Thus, while we are confident that these proteins exist in these specific contexts, we do not yet have evidence for their expression under normal cellular conditions. Given these considerations, GENCODE has not annotated these examples as *protein-coding genes* at this time, although they are clearly encoding proteins.

*What, then, is the importance of detected ncORF proteins that are not (yet) annotated as protein-coding genes?* We believe the identification of these newly-confirmed ncORF proteins is immensely important. Indeed, there have always been numerous experimentally detected proteins that are not annotated as protein-coding genes, such as cellular proteins encoded by transposons and retroviruses. For ncORFs, their proteins and HLA-I presented peptides may have direct biomedical relevance, which is manifested in the growing interest in targeting such cryptic peptides with cancer immunotherapy, including cellular therapies and therapeutic vaccines^19,82,83^. Additionally, the human genetics community has intensified scrutiny on how variants impacting ncORFs contribute to human genetic disease, which may also implicate these peptides^15,84^. Therefore, we deem inclusion of HLA presented ncORF peptides into the reference annotation ‘ecosystem’ to be important, and these data have now been integrated into our Ribo-seq ORF annotations.

The wider question then is how reference gene annotation should accommodate ncORFs that may not be *protein-coding genes* according to traditional paradigms, but are nonetheless protein-producing parts of the human genome. Such conversations are already occurring, and we hope that this work will help the scientific community engage in this discourse as well. Ultimately, as additional evidence becomes available – not only in the form of Ribo-seq, HLA peptidomic, proteomic, and transcriptomics datasets but also potentially via the advent of new technologies – both more comprehensive analyses and reappraisals of previous cases will be critical to continue to chart new paths for interpreting the human genome in physiology and disease.

## Conclusion

In summary, we report the international collaborative efforts of multiple global stakeholders in proteomics (PeptideAtlas/HUPO-HPP), immunopeptidomics (HIPP), Ribo-seq ORF discovery, and gene annotation (GENCODE) to initiate a continuing effort to develop a consensus understanding of protein-level evidence for 7,264 ncORFs. After searching 3.8 billion MS/MS spectra derived from ordinary protease digests from human cell lines, tissues, and fluids, as well as immunopeptidomics datasets, we manually validated evidence for 1,715 ncORFs as having compelling evidence of translation by proteomics, warranting further exploration. This evidence is now being used to advance the status of annotation efforts for ncORFs, and we hope that our results will serve as an initial reference catalog of known peptides that support ncORF translation.

## Acknowledgements

We are grateful to Prof. Pavel Baranov of the University College Cork for suggestions and critical comments during the preparation of this manuscript. This work was funded in part by the National Institutes of Health grants R01 GM087221 (EWD, RLM), U19 AG023122 (RLM), S10 OD026936 (RLM) and by the National Science Foundation grants DBI-1933311 (EWD) and MRI-1920268 (RLM). European Union’s Horizon 2020 research and innovation programme under the Marie Sklodowska-Curie grant agreement No. 945405 (IFM). J.R.P. acknowledges funding from the National Institutes of Health / National Cancer Institute [K08-CA263552-01A1]; the V Foundation for Cancer Research [V2024-013]; Alex’s Lemonade Stand Foundation Young Investigator Award [21-23983]; Hyundai Hope on Wheels Foundation; the Yuvaan Tiwari Foundation; DIPG/DMG Research Funding Alliance; Book for Hope Foundation; Curing Kids Cancer Foundation [20-3388093], and the Andrew McDonough B+ Foundation [1185689]. J.R.P. is the Ben and Catherine Ivy Foundation Clinical Investigator of the Damon Runyon Cancer Research Foundation [CI-127-24]. J.A.V. acknowledges funding from the Wellcome Trust [223745/Z/21/Z] and from the European Molecular Biology Laboratory (EMBL) core funding. I.F.M acknowledges funding from the EU Horizon 2020 programme (Marie Sklodowska-Curie grant agreement No. 945405 - ARISE programme). S.v.H. acknowledges funding from Fonds Cancers (FOCA, Belgium), Stichting Reggeborgh (the Netherlands), and Villa Joep. This publication is part of the project “Evolutionarily young microproteins in childhood brain cancer” (with project number VI.Vidi.223.022 of the research programme NWO talent programme Vidi, which is (partly) financed by the Dutch Research Council (NWO), awarded to S.v.H. Research reported in this publication was supported by Oncode Accelerator, a Dutch National Growth Fund project under grant number NGFOP2201, awarded to S.v.H. T.F.M. acknowledges financial support from NIH grant K01CA249038 J.G.A and S.C. are supported in part by grants P01CA206978 from the NIH, and grants U24CA270823 and U01CA271402 from National Cancer Institute (NCI) Clinical Proteomic Tumor Analysis Consortium program, as well as a grant from the Dr. Miriam and Sheldon G. Adelson Medical Research Foundation to S.A.C. NH was supported by ERC Advanced Grant (EU Horizon 2020, AdG788970), Deutsche Forschungsgemeinschaft (SFB 1470, B03), and EU Horizon 2020 Pathfinder Program.

P.F. was supported by a Victorian Cancer Agency Mid-Career Fellowship and the National Health and Medical Research Council of Australia (NHMRC). J.M.M. is supported by the Wellcome Trust (grant number 108749/Z/15/Z), the National Human Genome Research Institute (NHGRI) of the US National Institutes of Health (NIH) under award number 2U41HG007234, and EMBL core funding. E.B. is funded by the National Human Genome Research Institute (NHGRI) grant U24HG00334. The content is solely the responsibility of the authors and does not necessarily represent the official views of the National Institutes of Health. Ensembl is a registered trademark of EMBL.

## Author Contributions

Conceptualization: E.W.D., L.W.K., J.M.M., R.L.M., J.R.P., S.v.H.; methodology, E.W.D., L.W.K., J.M.M., R.L.M., J.R.P., S.v.H., I.F-M., J.R-O., N.T., M.B-S., J.C., J.A.V.; formal analysis, E.W.D., L.W.K., J.M.M., J.R-O., I.F-M., Z.S., S.C.; investigation, E.W.D., L.W.K., J.M.M., J.R-O., I.F-M., Z.S.; resources, R.L.M., J.R.P., S.v.H.; data curation, E.W.D., L.W.K., J.M.M., R.L.M., J.R.P., S.v.H.; writing - original draft, E.W.D., L.W.K., J.M.M., R.L.M., J.R.P., S.v.H., J.R-O., I.F-M., J.C., M.B-S., J.A.V., N.T.; writing - review & editing, E.W.D., L.W.K., J.M.M., R.L.M., J.R.P., S.v.H., J.G.A., M.M.A., J.L.A., M.A.B., S.C., A.A.B., E.A.B., L.C., S.A.C., J.C., A-R.C., K.D., P.F., N.H., N.T.I., M.M., M.J.M., T.F.M., G.M., U.O., S.O., O.R., X.R., S.A.S., E.V., A.W., J.S.W., W.W., Z.X., J.R-O., I.F-M., Z.S., J.C., M.B-S., J.A.V., N.T.; visualization, E.W.D., L.W.K.; supervision, R.L.M., J.R.P., S.v.H.; project administration: R.L.M., J.R.P., S.v.H.; funding acquisition, R.L.M., J.R.P., S.v.H.

## Methods

### Data availability and code availability statement

All mass spectrometry data in this manuscript is publicly available through the PeptideAtlas database at https://peptideatlas.org/ and ProteomeXchange (https://proteomecentral.proteomexchange.org/. Specific dataset identifiers are listed in **Supplementary Table S1**. All ribosome profiling data inspected in this manuscript are publicly viewable at GWIPS-viz, as detailed in the methods. Code generated for this manuscript is posted on https://github.com/VanHeeschLab/deutsch_kok_et_al_2024. The code for the Multi-Layer Perceptron Classifier Model can be accessed via https://git.embl.de/ivfimo/machine_learning_scripts.

### PeptideAtlas database construction and searching

The Human non-HLA PeptideAtlas 2023-06 build contains 295 ProteomeXchange datasets (PXDs)^36^ split into 1,172 different experiments and that comprise a total of 3.5 billion MS/MS spectra. Sequence database searching was performed with MSFragger^85^

3.7 using search parameters appropriate for each dataset, depending on alkylation, labeling, fragmentation type, instrument, enrichment strategy, and more. All datasets were searched with semi-enzymatic settings (typically semi-tryptic). The search database was 2023-02 THISP level 4 database^37^ (https://peptideatlas.org/thisp/), which included the 7,264 Ribo-seq ORFs from Mudge et al.^29^ as well as other contributed sequences that might be translated. All datasets were searched with generic artifactual variable modifications methionine oxidation, protein N-terminal acetylation, peptide N-terminal pyro-glutamic acid from glutamic acid or glutamine, and asparagine and glutamine deamidation. The alkylation modification was set as a fixed modification (typically carbamidomethylated cysteine).

The statistical validation of the results for each experiment was performed with the TPP^38,39^ 7.0 tools PeptideProphet^86^, iProphet^87^, and PTMProphet^88^, the results mapped to the human proteome with ProteoMapper^89^ taking known variants into account and to the genome with the ENSEMBL^90^ toolkit as previously described^91^. A complete list of datasets used and a summary of the search results in each build are available via hyperlinks found at https://peptideatlas.org/builds/human/non-hla/. **Supplementary Table S9** provides FDR metrics at the PSM-, peptide-, and protein-levels, as well as for certain subsets of proteins, including the neXtProt core proteome, the 7264 Ribo-seq ncORFs, as well as all CONTRIB sequences, many of which are putative ncORFs.

The Human HLA PeptideAtlas 2023-11 build comprises a set of 118 HLA immunopeptide-enriched publicly available PXDs, which we split into 592 separate experiments, containing 240 million MS/MS spectra from 9776 MS runs (**Supplementary Table S6**). Sequence database searching was performed with MSFragger v3.7 using search parameters appropriate for each dataset, depending on sample handling. All datasets were searched in no-enzyme mode. While some HLA peptides have a lysine or arginine on the C terminus, and thus exhibit fragmentation patterns typical of tryptic peptides, many HLA peptides do not have such characteristics and thus their spectra may have strong b ions and internal fragmentation ions, rather than strong y ions, which are customary in tryptic peptide spectra. **Supplementary Table S10** provides FDR metrics at the PSM-, peptide-, and protein-levels, as well as for certain subsets of proteins, including the neXtProt core proteome, the 7264 Ribo-seq ncORFs, as well as all CONTRIB sequences, many of which are putative ncORFs.

The search database was 2023-07 THISP level 4 database^37^ (https://peptideatlas.org/thisp/), which included the 7,264 Ribo-seq ORFs from Mudge et al.^29^ as well as other contributed sequences that might be translated. 299 common contaminants based on the list from Frankenfield et al.^92^ minus the human proteins are included in the search database (available at https://peptideatlas.org/thisp/). All datasets were searched with generic artifactual variable modifications methionine oxidation, cysteine cysteinylation, protein n-terminal acetylation, peptide n-terminal pyro-glutamic acid from glutamic acid or glutamine, and asparagine and glutamine deamidation. Static carbamidomethylation of cysteine was set for experiments that used Iodoacetamide. For samples that were treated with tandem mass tag (TMT) or SILAC or enriched for phosphorylated peptides, appropriate mass modifications were applied. Statistical validation was performed as described above by the TPP. A complete list of datasets used and a summary of the search results in each build are available via hyperlinks found at https://peptideatlas.org/builds/human/hla/.

### Protein identifications and categories

Peptides are preferentially mapped using ProteomeMapper^89^ (in TPP 7.0) to the 20,389 entries (“core proteome”) and their isoforms of the 2023 version of neXtProt^93^ taking into account all single amino acid variants encoded in neXtProt. Proteins that have 2 or more uniquely mapping non-nested (contained completely within the other) peptides of length 9 or more amino acids, together covering at least 18 amino acids are categorized as “canonical” by PeptideAtlas. If a protein entry meets the above 2-peptide criteria with peptides that cannot be mapped to the core proteome, they are termed “non-core canonical”. There are 9 additional categories, including “indistinguishable representative”, “indistinguishable”, “representative”, “marginally distinguished”, “subsumed”, “weak”, “insufficient evidence” for various scenarios of ambiguous and redundant evidence. Finally, the categories “identical” are assigned to entries that are sequence-identical to another entry, and proteins that have no peptide evidence whatsoever are categorized as “not detected”. See van Wijk et al.^94^ for an extensive description of the PeptideAtlas protein categories. For reasons of integration with the HPP annual metrics^32–34^, only sequence entries that belong to the core set of ∼20,389 neXtProt^93^ and UniProtKB/Swiss-Prot^2^ protein coding genes can achieve canonical status.

### Manual inspection of ORF MS spectra

Despite extraordinary efforts to minimize false positives, both builds do contain some false positives, and they are most easily found mapping to proteins that are unlikely to be detected. For gene annotation purposes, manual inspection is therefore crucial to ensure that few false positives are reported for extraordinary detections, as described extensively by Deutsch et al. (2019)^40^. We manually inspected each of the peptides corresponding to ncORFs and provided a manual categorization as well as a commentary. The manual categories are as follows: “excellent” (highly compelling evidence that the peptide identification is completely correct); “good” (the PSM is likely correct but lacks sufficient quality and coverage of the residues to provide highly compelling evidence); “false positive”; “close but false positive” (the PSM has many matching ions and is likely to be almost the correct peptidoform, but slight discrepancies indicate that the true identification is very close but not quite the listed sequence); “low information” (the ions that are detected are compatible with the identification, but coverage is too low to be compelling). The best peptide-spectrum match is also listed in the Supplementary Tables as a USI that can be resolved and viewed at https://proteomecentral.proteomexchange.org/usi/, or in cases where a USI cannot be achieved, a direct URL for the spectrum in the PeptideAtlas web interface. In any case, all protein entries, peptides, and spectra may be browsed via the PeptideAtlas web interface starting at the URLs provided above.

### Procedure for manually validating peptide spectrum matches

1. Obtain a listing of PSMs for a given peptide in PeptideAtlas
2. Examine PSMs until at least one PSM provides excellent evidence, and record its USI (Universal Spectrum Identifier) if available, or PeptideAtlas spectrum viewer URL if a USI is not available. For spectra without a PXD number associated with the dataset, a USI is usually not available. This is most common in Clinical Proteomic Tumor Analysis Consortium (CPTAC) datasets, for which a PXD has not been assigned. PSMs with USIs should be checked at https://proteomecentral.proteomexchange.org/usi/.
3. Evaluate the PSM as follows. To obtain the “excellent” rating:

a. The combination of b-ion and y-ion series must yield nearly complete coverage of the proposed peptidoform explanation. For tryptic or tryptic-like peptides (a basic residue on the C terminus), this will typically mean a nearly complete y-ion series and a b ion series that begins at b2 and at least meets the y ion series. For the rules above and below for tryptic-like ions, swap y-ion for b-ion when there is a basic residue instead on the N terminus.
b. If there are any prominent peaks beyond the last matching ion peak, suggesting that the sequence should extend with different residues, the PSM is not “excellent”.
c. Any gaps in the y or b ion series must not have a plausible unannotated candidate in the gap, implying that the true identification is slightly different than the proposed identification. Such a plausible unannotated candidate must have a mass defect between the ions before and after the gap.
d. Gaps should have a plausible explanation for low intensity, such as a y ion C terminal to a proline.
e. For tryptic-like peptides, the y ions N terminal to a proline should be more intense than surrounding ions, although confounding factors such loss of sensitivity at the high m/z end or other nearby prolines should also be considered.
f. Strong b2 and corresponding a2 diketopiperazine ions are preferred in HCD spectra. There may be a gap at b1 ions as these are usually not visible unless there is an N terminal mass modification.
g. Internal fragmentation ions should be considered when annotating peaks, especially for peptides without basic residues at either terminus.
h. Mass modifications should be kept to a minimum.
i. For long peptides especially, there must not be a substantial region with no ions.
j. There should be no prominent unannotated peaks that suggest contamination or misassignment. Internal fragmentation ions and neutral losses should be considered for peaks that are not attributable to ordinary b and y ions.

### Gene annotation

The gene annotation work in this study has been carried out as part of the ongoing GENCODE project using existing workflows^1^.

### Annotating immunopeptidomics MS-runs

All HLA-I MS-runs were annotated for the source material (cancer vs. non-cancer and cell line vs. non-cell line) (**Supplementary Table S6**). These annotations were largely based on what was documented by PeptideAtlas, but for several instances the category was changed based on data in the publication corresponding to the MS-run. HLA typings of MS-runs were determined by manually searching the publications corresponding to each MS-run. For 4,879 MS-runs the full four-digit HLA typing could be retrieved.

### Categorizing HLA peptides

Starting with the 865,922 peptides from the Human HLA PeptideAtlas 2023-11 build, 99 peptides starting with “LLLLLLL”, “PPPPPPP” or “QQQQQQQ” were filtered out. Mappings to entries starting with DECOY, CONTRIB_smORFs_Cui, CONTRIB_sORFs, CONTRIB_Fedor, CONTRIB_Bazz, CONTRIB_HLA, CONTRIB_GENCODE_nearcognate were ignored. All peptides with a length of at least 8 amino acids, mapping to UniProtKB/Swiss-Prot entries with at most 30 distinct mappings were considered to be derived from canonical proteins. Peptides with a length of at least 8 amino acids, mapping to ncORFs and not canonical proteins, with at most 10 distinct mappings were considered to be derived from ncORFs. For peptides with mappings against multiple ncORFs, one ncORF was selected based on the first one alphanumerically. All remaining ncORFs were put in the ‘Other mappings’ category.

### ncORF expression in cancer tissues

To determine whether ncORFs were preferentially expressed in cancer or non-cancer tissues, each ncORF peptide was categorized to originate exclusively from MS-runs from cancer samples, exclusively from MS-runs from non-cancer samples or from both. Additionally, each ncORF (and corresponding peptides) were classified to originate from a cancer gene based on the Cancer Gene Census genes (accessed January 4th 2024)^95^.

### HLA binding predictions

Binding predictions were performed with NetMHCpan 4.1^4596^. Predictions were done for MS-runs with a known four-digit HLA-typing. For nine MS-runs with A24:01, B43:01, or C12:01 as one of the alleles, no predictions could be made because these alleles were not known to NetMHCpan. **Supplementary Table S6** shows an overview of MS-runs for which binding predictions were made. Peptides were predicted to bind to an allele if the rank score was smaller than or equal to 2. If the HLA-typing of an MS-run consisted of multiple alleles, the peptide was assigned to the allele with the lowest predicted rank score, irrespective of whether this rank score was smaller than 2 or not.

### Detectability determinants

Canonical proteins were categorized as detected and undetected based on whether they were detected by a single peptide. Canonical proteins shorter than 16 aa and proteins with amino acid symbol “U” in their sequence were filtered out. ncORFs sequences were categorized similarly to the canonical proteins. Contrary to most other analyses, peptides were not exclusively assigned to a single ncORF, due to which the number of detected ncORFs was larger than in **Figure S2a**. For the ncORF analysis taking into account only the first or last 30% of the sequence, the requirement was that this 30% was again 16 aa long. Significance was determined by the two-sided Wilcoxon test. Values were adjusted using Bonferroni multiple testing correction for the eight comparisons.

### Hydrophobicity analysis

For the hydrophobicity analysis, all sequences were aligned by the C-terminus. Starting at that position and moving towards the N-terminus, the average hydrophobicity of the 15 previous amino acids across the sequences was determined. For every position, only sequences long enough to still contain 15 amino acids before the position were taken into account. A line was fit through measurements using Local Polynomial Regression Fitting. 95% confidence intervals were determined using a two-sided T-test.

### Expression analysis

To compare the expression of detected and undetected ncORFs, we used data from GTEx^48^. The mean FPKM of all genes per tissue (excluding testis) was used. Tissues from the same organ (e.g. all brain derived tissues) were grouped together. For each ncORF, the expression was determined using the gene ids. Contrary to most other analyses, peptides were not exclusively assigned to a single ncORF, due to which the number of detected ncORFs was larger than in **Supplementary Figure S2a**. However, for 326 ncORFs the associated gene id was not present in GTEx, so these were excluded.

### Tissue comparison

For comparing the expression of ncORFs in tissues, the data from the HLA Ligand Atlas (PXD019643) was used^50^. Tissue names were extracted from the MS-run file names. For each tissue, the number of distinct ncORF and CDS peptides was determined, as well as the percentage of ncORF peptides. Statistical significance was determined using multiple Fisher’s exact tests and Bonferroni multiple testing correction for the 30 tissues. Gene expression levels were determined using mean FPKM values per gene across tissues from GTEx^48^. Only genes which expressed ncORFs in the HLA Ligand Atlas that were present in GTEx were considered. A selection of GTEx tissues that showed resemblance to the HLA Ligand Atlas tissues was used. Resemblance was based on the similarity of the HLA Ligand Atlas and GTEx tissue names.

### ncORF visualization

For the visualization of the ribosome profiling data of ncORFs, GWIPS-viz data was used (accessed July 5th 2024)^97^. For initiation p-sites, for the group ‘Initiating Ribosomes (P-site)’ and track ‘Global Aggregate’, all corresponding tables were downloaded and the p-sites were merged. For the global a-sites, for the group ‘Elongating Ribosomes (A-site)’ and track ‘Global Aggregate’, all corresponding tables that were not used for the initiation a-sites were downloaded and the a-sites were again merged.

### Analysis of Ribo-Seq data

We manually inspected Ribo-seq data for 183 ncORFs with at least one peptide nominated in the nonHLA build and 699 ncORFs with at least one peptide nominated in the HLA build. We used the GWIPS-viz browser^97^ to assess evidence of ncORF translation with a publicly-accessible web portal that enables the research public to examine our assessment of these ncORFs independently. The GWIPS-viz parameters were: the Elongating Ribosomes (A-site) with the Global Aggregate track on “Full”, which reflects native Ribo-seq; and the Initiating Ribosomes (P-site) with the Global Aggregate track on “Full”, which represents Ribo-seq signal from ribosomes enriched at initiation sites. We independently evaluated the native Ribo-seq and “initiation” Ribo-seq data. For each of these data types, we classified the data as “Insufficient”, “Sufficient” or “Excellent” for supporting the translation of a given ncORF. A given ncORF was considered to be verified at the level of Ribo-seq data if either the Elongating Ribosomes or Initiating Ribosomes track data was “Sufficient” or “Excellent”. We defined “Excellent” if there were four sequential clearly identified Ribo-seq peaks in-frame within the first 100 nucleotides of the ncORF. We defined “Sufficient” if there were three sequential clearly identified Ribo-seq peaks in-frame within the first 100 nucleotides of the ncORF. We defined “Insufficient” if there were not clearly sequential in-frame reads. We additionally collated selected ncORFs in the GENCODE set that were first identified in Gaertner et al.^98^ and van Heesch et al.^7^ but considered “Insufficient” in the GWIPS-viz database. Because GWIPS-viz does not include the data for these two studies, we evaluated the raw data for these selected ncORFs in the primary datasets and categorized their support according to these data. We additionally calculated the percentage of in frame ribosome footprints (PIF) and uniformity of ribosome coverage for each of the ncORFs supported by one or more peptides in each PeptideAtlas build, as observed in the human body map.

### Employment of the Tier classification system

We applied the Tier classification system for ncORFs initially proposed in Prensner et al.^43^. Specifically, ncORFs were given an Initial or Provisional Tier based on the information available from the large-scale mass spectrometry search. Following manual review of the nomination data, ncORFs were then assigned a Final Tier. Tiers were defined as follows:

- Tier 1A: Two non-nested peptides in MS proteome data, with or without HLA immunopeptidomics data, with Ribo-seq data
- Tier 1B: Two non-nested peptides in HLA immunopeptidomics data with Ribo-seq data
- Tier 2A: One peptide in MS proteome data, with or without HLA immunopeptidomics data, with Ribo-seq data
- Tier 2B: One peptide in HLA immunopeptidomics data with Ribo-seq data
- Tier 3: Any HLA immunopeptidomics and/or tryptic proteome LC-MS/MS evidence without Ribo-seq evidence
- Tier 4: Ribo-seq evidence without proteomic evidence
- Tier 5: *In silico* prediction of an ORF on an expressed transcript without any Ribo-seq or proteomic evidence

### Multi-Layer Perceptron (MLP) Classifier Model

A dataset comprising 677 ncORF peptide sequences of 9 amino acids, each annotated with 22 attributes, was utilized to develop a Multi-Layer Perceptron Classifier model. The implementation was carried out using Python 3 and the following software libraries: pandas, numpy, matplotlib, and scikit-learn. The dataset was processed by separating the features from the target variable. The data was then split into training and testing sets, with 80% allocated for training and 20% for testing. To ensure reproducibility, the random state was set to 42 during the split. Prior to model fitting, the features were standardized using StandardScaler. This preprocessing step involved removing the mean and scaling the features to have unit variance, thereby normalizing the data. The MLP Classifier model was initialized with a maximum of 8000 iterations and a random state of 42 to ensure reproducibility. The model was then subjected to hyperparameter tuning using grid search with cross-validation. The hyperparameters explored included: hidden layer sizes: (280,), activation function: ‘tanh’, and regularization parameter (alpha): 0.01. Grid search with cross-validation was employed to systematically evaluate the performance of various hyperparameter combinations and identify the optimal configuration. The best-performing model, as determined by the grid search results, was selected and fitted to the training data. Subsequently, this model was used to make predictions on the test set, which had not been seen by the model during training. The performance of the model was assessed using standard evaluation metrics to determine its predictive capabilities.

### TensorFlow-Keras Model

Due to 1785 ncORFs being detected while 5479 remain undetected, presenting an approximate ratio of 1:3, a balanced weight for imbalanced dataset was used to address the imbalance and a neural network analysis to build, train, and evaluate a TensorFlow-Keras model. The dataset included 7264 ncORF amino acid sequences with a selection of 43 attributes. Using Python 3 we imported the necessary libraries including TensorFlow, Keras, and various components of TensorFlow and Keras for building and evaluating the model. Using the train_test_split function from scikit-learn, we allocated 80% for training and 20% for testing after separating the training and testing sets from the target variable. The features were standardized using the StandardScaler. A sequential model consisting of multiple layers was built with an input of 16 neurons, ReLU activation, and L2 regularization. To prevent overfitting we added batch normalization and dropout layers after each hidden layer. The output layer consisted of a single neuron with sigmoid activation for binary classification. We compiled the model using the Adam optimiser with a learning rate of 0.001, binary cross-entropy loss function, accuracy as the metric, and it was trained on the training data. During training it was run for a total of 60 epochs with a batch size of 12. The target variable for the test set was predicted by the model.

## Supporting Information

### Supplementary Tables

Supplementary Table S1:List of experiments in the non-HLA build and the protease used.

Supplementary Table S2: Listing of peptides that are mapped to Ribo-seq ncORFs from the Human non-HLA PeptideAtlas 2023-06 build

Supplementary Table S3: Listing of Ribo-seq ncORFs annotated in the Human non-HLA PeptideAtlas 2023-06 build

Supplementary Table S4: Listing of peptides that are mapped to Ribo-seq ncORFs from the Human HLA PeptideAtlas 2023-11 build

Supplementary Table S5: Listing of detected Ribo-seq ncORFs from the Human HLA PeptideAtlas 2023-11 build

Supplementary Table S6: List of HLA build MS runs, including the HLA type of each MS run.

Supplementary Table S7: List of 677 HLA-I peptides, including their sequence, best allele, the 22 features that the model used for training, and the output probabilities from the model.

Supplementary Table S8: List of 7,264 ncORFs along with the features that were used to train machine learning models and output probabilities of the model.

Supplementary Table S9: FDR metrics for the non-HLA build analysis Supplementary Table S10: FDR metrics for the HLA build analysis

### Supplementary Documents

Supplementary Document S1 - Discussion of ncORF detections in non-HLA data

Supplementary Document S2 - Discussion of machine learning results predicting detectability of ncORFs.

### Supplementary Figures

**Supplementary Figure S1.**
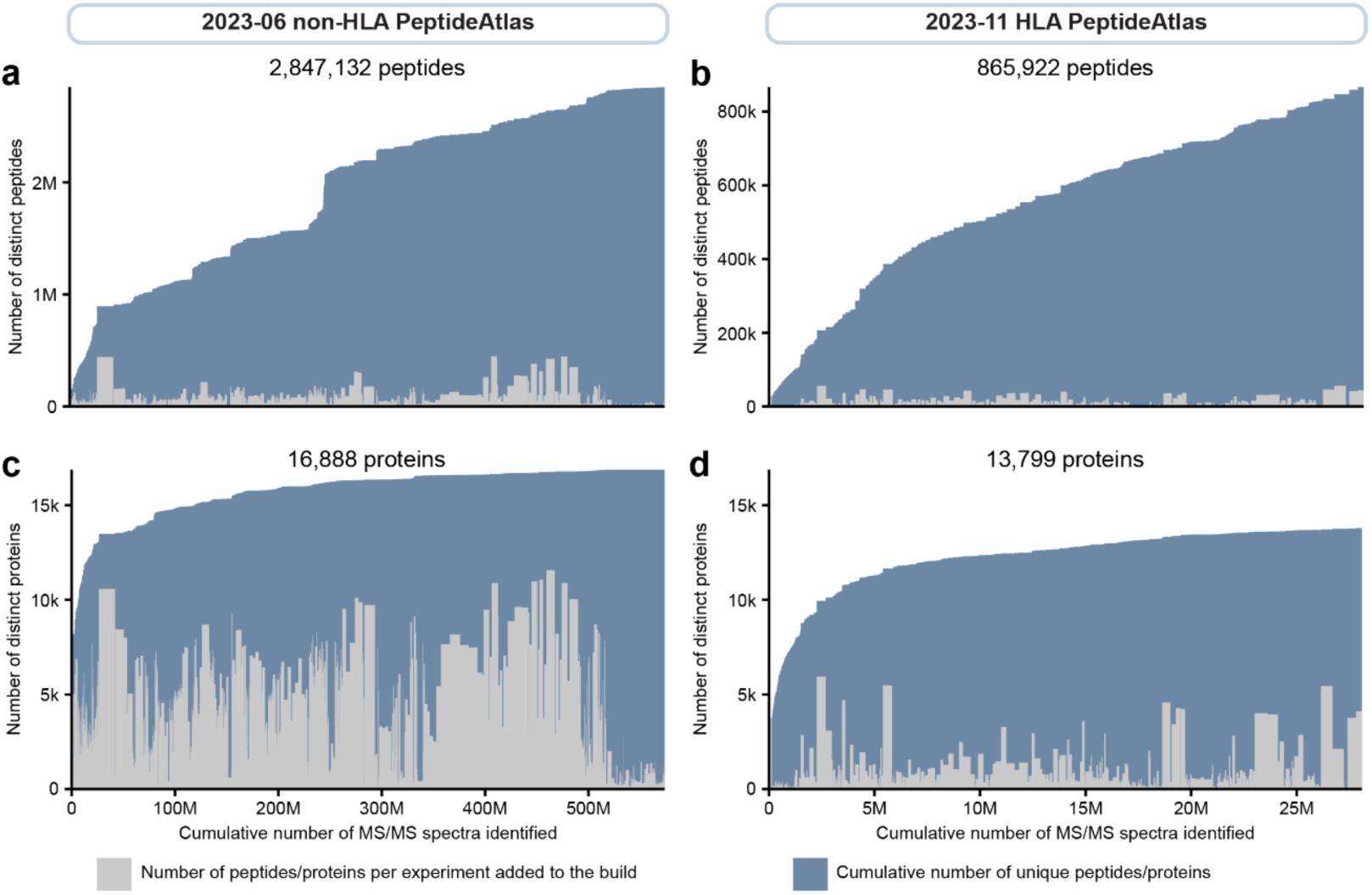
The number of distinct peptides and proteins as datasets were added to the Human non-HLA (left) and HLA (right) PeptideAtlas. (**a**) Over 2.8 million distinct peptides have been observed in the 573 million PSMs in the non-HLA build. Each rectangle is one of the 1,172 experiments. Blue rectangles represent the cumulative number of distinct peptides in the build, while the gray rectangles depict the total number of distinct peptides within each experiment. **(b**) Over 0.86 million distinct peptides have been observed in the 28 million PSMs in the HLA build. Each rectangle is one of the 592 experiments. Blue rectangles represent the cumulative number of distinct peptides in the build, while the gray rectangles depict the total number of distinct peptides within each experiment. (**c**) The blue rectangles depict the cumulative 16,888 canonical proteins that have been cataloged in the 2023-06 Human non-HLA PeptideAtlas, whereas the gray rectangles show the total number of proteins present in each of the 1,172 experiments. (**d**) The blue rectangles depict the cumulative 13,799 canonical proteins that have been cataloged in the 2023-11 Human HLA PeptideAtlas, whereas the gray rectangles show the total number of proteins present in each of the 592 experiments. Although the total number of peptides continues to increase steadily, progress in the number of proteins is now very slow. Over the last 100 million PSMs, the cumulative counts are increasing by ∼2,000 peptides per million PSMs and ∼1 newly identified protein per million PSMs.

**Supplementary Figure S2.**
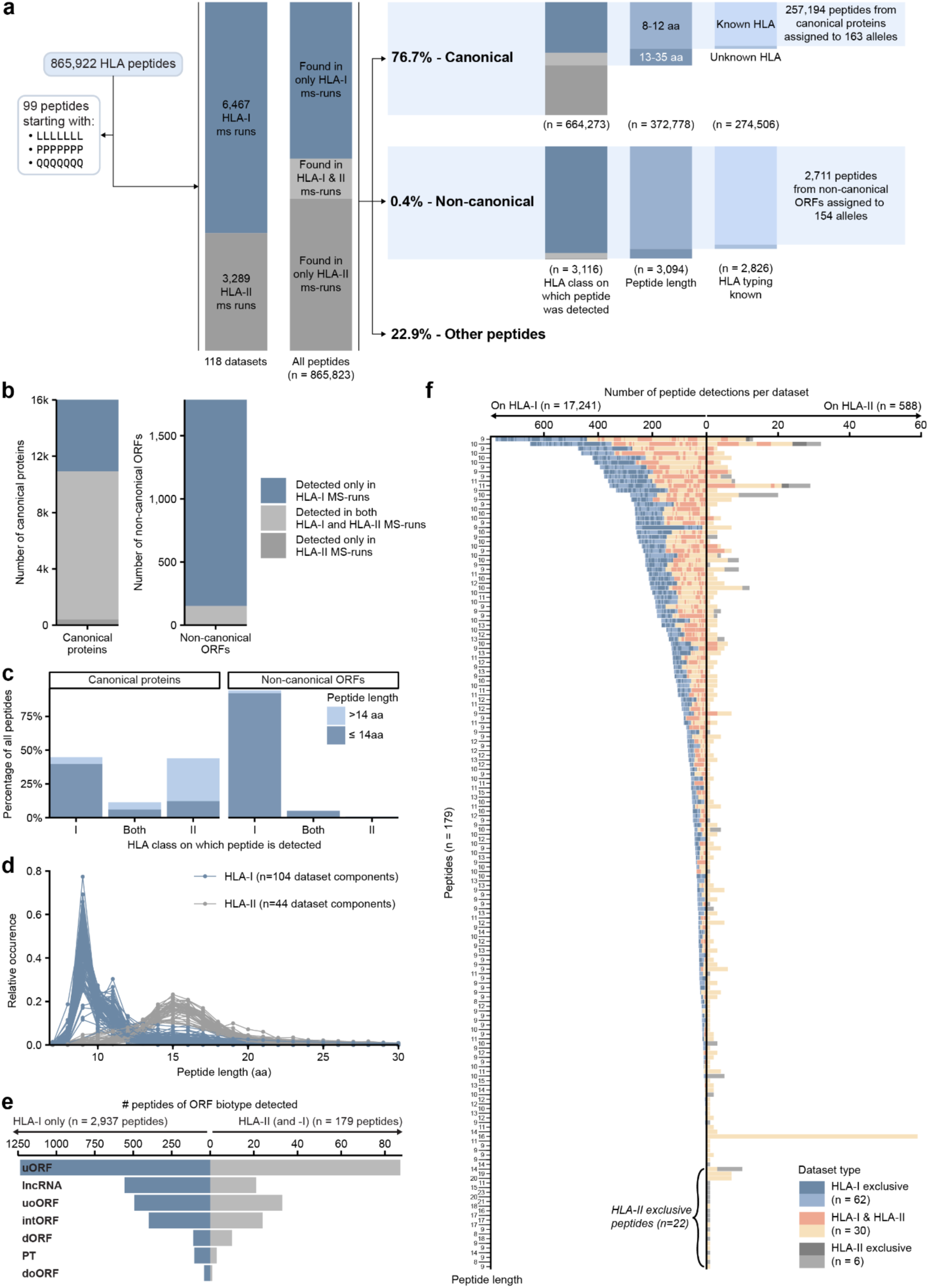
Detection of ncORF peptides in HLA-I and HLA-II, and in cancer and non-cancer samples. (**a**) Schematic illustrating the total numbers of peptides (from both normal proteins and ncORFs) extracted from the total set of peptides. Depending on the analysis, peptides were further selected to those detected on HLA-I, had a length from 8-12 amino acids, and originated from an MS run with a known HLA-typing. The counts below each bar denote the number of distinct peptides. The distinction between “canonical”, “non-canonical”, and “other peptides” is defined in the methods. (**b**) Barplots showing for the detected canonical proteins (left) and detected ncORFs (right) whether their detected peptides were exclusively detected in HLA-I or HLA-II MS-runs, or in both. **(c)** Barplots showing for canonical proteins and ncORFs the percentage of peptides found exclusively in HLA-I and HLA-II MS-runs, or in both. Bars are colored by the peptide length being ≤14aa, or >14aa in length. **(d)** Line graph showing the peptide length distribution per dataset component split by HLA-class. **(e)** Bar plot showing the number of peptides detected per ORF exclusive to HLA-I samples (left) and those present in HLA-II samples, possibly in addition to HLA-I samples (right). Please note the x-axis scales differ by an order of magnitude between the left and right part of this panel. HLA-I and HLA-II peptide detection is not mutually exclusive as HLA-I peptides might be accidentally recovered from HLA-II pulldown experiments. **(f)** The frequency of peptide detection in HLA-I MS-runs (left) and HLA-II MS-runs (right) per peptide. Each alternating shade corresponds to a different dataset, with shades grouped by dataset type. The left axis denotes peptide lengths. Please note x-axis scales differ by an order of magnitude between the left and right part of this panel. Only peptides detected in at least one HLA-II sample are included. 22 of the 179 distinct peptides were exclusively detected in HLA-II samples. Fourteen of these peptides have a length 14 amino acids or greater, suggesting a potential presentation by HLA-II. This is still a minority in contrast to the total amount of 3,116 non-canonical ORF derived HLA peptides.

**Supplementary Figure S3.**
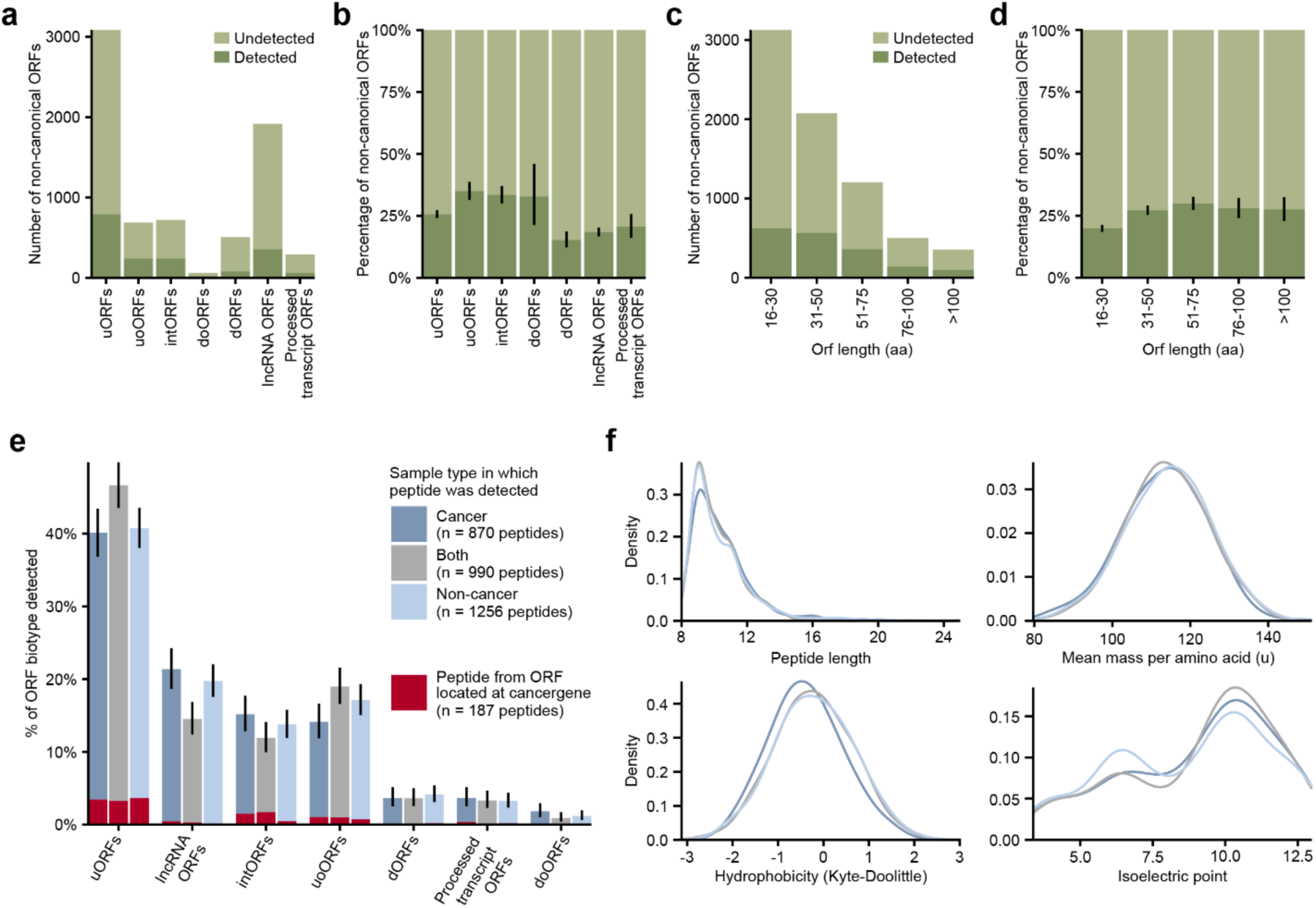
(**a-d**) Comparisons of the detected and undetected non-canonical ORFs. (**a**) The total number of non-canonical ORF per ORF type and the number of ORFs for which a peptide was observed. (**b**) As in (**a**), but now shown in percentages. (**c**) The total number of non-canonical ORF grouped by length and the number of ORFs for which a peptide was observed. (**d**) As in (**c**), but now shown in percentages. Black lines on bar graphs indicate 95% confidence intervals. (**e**) Bar plot showing the proportion of Ribo-seq ORF-derived HLA peptides detected per biotype, categorized by whether the peptide was exclusively identified in immunopeptidomics analyses of cancer tissues or cell lines, non-cancer samples, or both. No significant changes in ORF biotype recovery are observed between these sample types. Peptides originating from ncORFs located on a known cancer gene are colored red. Black lines on bar graphs indicate 95% confidence intervals. (**f**) Density plots comparing the Ribo-seq ORF-derived HLA peptides differentiated on sample type (as depicted in (**e**)): cancer, non-cancer, or both. The plots compare peptides by their length, mass, hydrophobicity (Kyte-Doolittle scale), and isoelectric point. No significant changes between the distributions of these density plots can be observed.

**Supplementary Figure S4.**
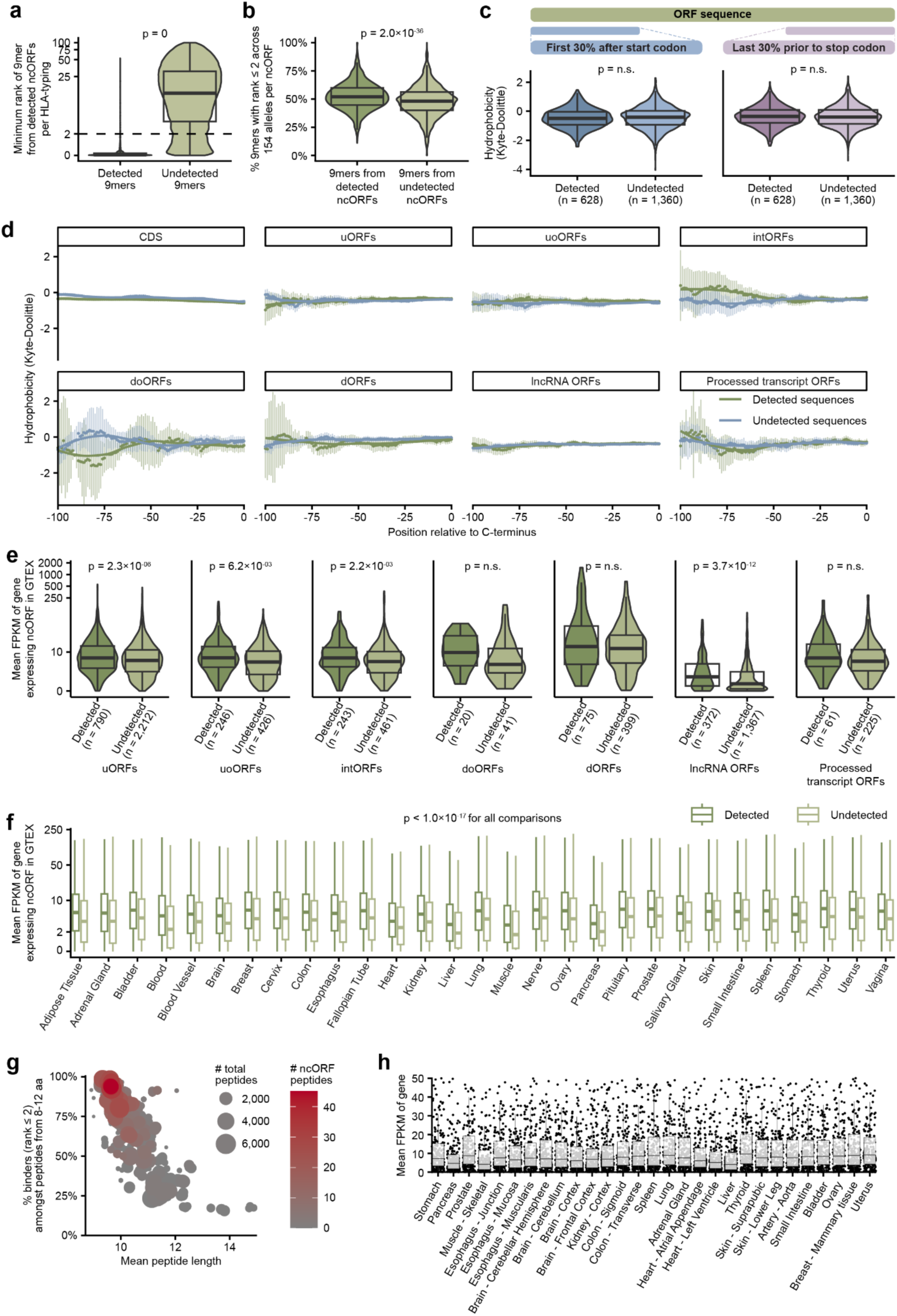
Potential determinants of ncORF detection. **(a)** Violin plot comparing for all MS-runs grouped by HLA-typing the minimum binding prediction rank for detected and undetected 9mers. **(b)** Violin plot comparing all 9mers from detected and undetected non-canonical ORFs. The y-axis shows per ORF the percentage of 9mers with a NetMHCpan rank ≤ 2 across all 154 alleles associated with ncORF peptides. **(c)** Violin plots similar to **(4a)** comparing the hydrophobicity by the Kyte-Doolittle scale between detected and undetected ncORFs for the first 30% of the ncORF sequence after the start codon, or the last 30% of the ncORF sequence. Statistical tests were performed with the two-sided Wilcoxon test, reported p-values were adjusted for multiple testing with Bonferroni correction. **(d)** Comparison of the hydrophobicity similar to **(4b)** between detected and undetected ncORFs/CDS per ncORF biotype. Each dot represents the average hydrophobicity of the amino acids at that position and the 14 amino acids before that position per ncORF biotype or CDS grouped by whether these were detected or not in the immunopeptidomics data. The lines were fitted using Local Polynomial Regression Fitting. Vertical bars represent 95% confidence intervals. Note that because ncORFs are mostly smaller than 100 aa, confidence intervals get larger with increasing C-terminus offset. **(e)** Comparison of the expression levels of detected and undetected ncORFs similar to **(4c)**, but split per biotype. On the y-axis, the mean FPKM in GTEX of genes expressing an ncORF is shown on a pseudo-log scale. 326 ncORFs for which the gene id was not present in GTEX are not shown. Significance was determined using two-sided Wilcoxon tests, reported p-values were adjusted for multiple testing with Bonferroni correction. **(f)** Comparison of the expression levels of detected and undetected ncORFs similar to **(e)**, but split per tissue. Outliers are not shown in the graph. Significance was determined using two-sided Wilcoxon tests, and p-values were adjusted for multiple testing with Bonferroni correction. All comparisons were found to be significant. **(g)** Dot plot similar to **(3e)**, for MS-runs originating from the HLA-ligand-atlas. The plot visualizes the correlation between mean peptide length and the percentage of predicted binders amongst peptides with a length between 8 and 12 amino acids (NetMHCpan rank ≤ 2) per MS run. Dot size corresponds to the total number of peptides per MS-run. Dot color corresponds with the percentage of non-canonical ORF-derived peptides per MS-run. Statistical tests were performed with the two-sided Wilcoxon test, reported p-values were adjusted for multiple testing with Bonferroni correction. **(h)** Comparison of the GTEx expression of 224/277 genes from which ncORFs in the HLA ligand atlas originate (53 genes with ncORFs in the HLA ligand atlas were not present in GTEx). GTEx tissues comparable to those from the HLA ligand atlas were selected, and sorted in the same way as in **(4e)**. Each represents the mean FPKM of a gene across these tissue samples in GTEx. Only genes with a mean FPKM lower than 50 are plotted for clarity, but all 224 genes were included for the boxplots.

**Supplementary Figure S5.**
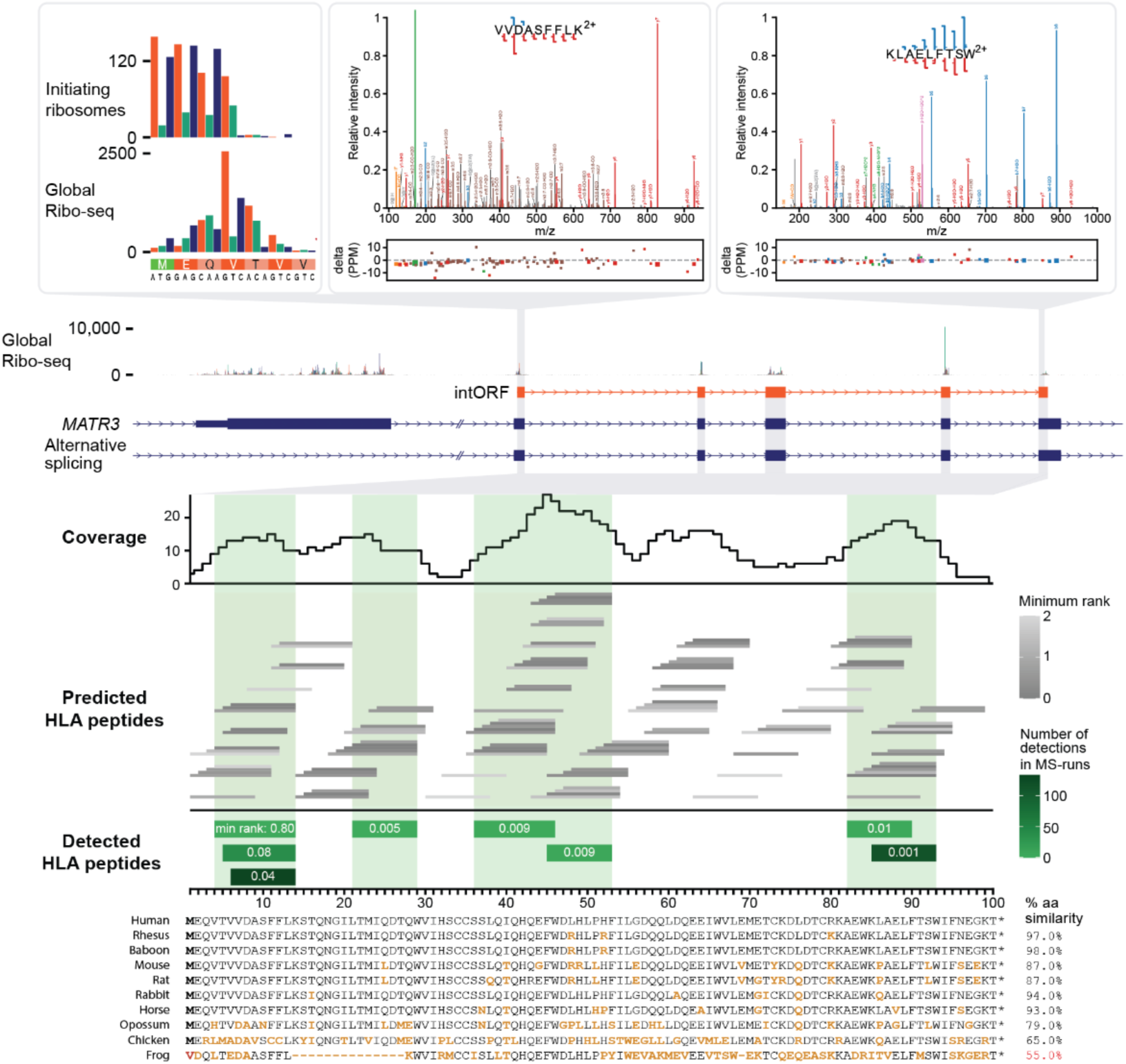
Overview of data available for c5norep142, an intORF in the *MATR3* gene. Ribo-seq data shows the initiation of translation at the methionine translation initiation codon (green), as determined by enrichment of ribosomes at the TIS. Two peptide spectral matches for HLA-I peptides VVDASFFLK and KLAELFTSW are shown having nearly complete sequence coverage (USIs are mzspec:PXD037270:Liv32_1176935F:scan:33690:VVDASFFLK/2 and mzspec:PXD011628:PBMC009_msms37:scan:16281:KLAELFTSW/2, respectively). The lowest panel shows the position of all 8 peptides that were observed in the immunopeptidomics data. The color shading indicates the number of MS runs in which each peptide was observed. The middle panel shows all peptides that are predicted with NetMHCpan to be observable in the MS runs (i.e. they are predicted to bind with NetMHCpan score <2 to at least one allele in one of the samples in which peptides were observed). The top part shows the number of predicted binding peptides in which each amino acid was located. Green shadings indicate which part of the ORF sequence was observed. Except for the region near the offset of 62, detected peptides occurred in the regions with the highest numbers of predicted binders.

