## Supplemental Document S1 for "High-quality peptide evidence for annotating non-canonical open reading frames as human proteins"

### Supplementary Document S1 - Discussion of ncORF detections in non-HLA data

This document provides extended discussion on the detections of ncORFs initially discovered by Ribo-seq (henceforth ncORFs) in non-HLA datasets, accompanying Supplementary Tables S2 and S3. The former table provides the complete list of non-HLA peptides matched to ncORFs. Each row represents an individual peptide, some of which have multiple peptide-spectrum matches (PSMs). For each peptide we list the peptide identifier, sequence, number of PSMs, and the ncORF entry to which it maps as taken from Mudge *et al*, as well as the ORF's PeptideAtlas category. We also provide the Universal Spectrum Identifier (USI)[1] of the best PSM for the peptide. All spectra are available in the PeptideAtlas interface for additional public scrutiny. Furthermore, USIs provide a mechanism with which the original spectrum can be viewed or downloaded for any participating ProteomeXchange resource that holds the spectrum, and annotated with the requested sequence. The easiest location to resolve USIs is at the ProteomeXchange ProteomeCentral webpage https://proteomecentral.proteomexchange.org/usi/.

Table S3 reconfigures this information based on the ncORF, including each ncORF with matched peptides as a separate row. Certain information fields are duplicated from Table S2. In addition, Table S3 includes the official symbol of the gene in which the ncORF is mapped and the GRCh38 coordinates of the translated sequence. We also include information on our manual reappraisal of the Ribo-seq evidence, which we consider to be an essential phase of the annotation process when it comes to the identification of prospective novel proteins.

As discussed in the main text, ncORFs appraised as Tier 1A are considered the strongest candidates for protein-coding annotation. Table S3 includes brief commentary on the annotation of such ncORFs as considered by GENCODE. In each case these comments represent a brief summary of more detailed information that is subsequently listed in this document. Moving forward, as part of our ongoing attempts to understand the broader annotation landscape, GENCODE have now manually examined all loci where the evidence is graded at Tier 2A or higher. By definition, none of these cases have enough peptide evidence to be validated as proteins as per HUPO-HPP guidelines. However, our gene annotation process found that 6 out of 46 Tier 2A ncORFs can be potentially reappraised as alternative protein isoforms. In fact, in certain cases, the existence of the Ribo-seq ORF as originally called and the existence of an overlapping alternative protein isoform may not be mutually exclusive. Table S3 includes commentary on all Tier 2A cases, which is expanded upon below.

Certain peptides map to additional sequences in the proteomics search space, which is substantially broader than our collection of 7,264 Ribo-seq ORFs. In particular, we find mappings to annotations provided by UniProtKB or RefSeq. Having examined these in detail, we find that in almost all cases the alternative annotations are derived from the same locus as our ncORF, i.e. the peptide is (in effect) mapped to the same genomic sequence. Thus, these are not multimappings, i.e. cases where a peptide can genuinely be mapped to more than one genomic location, as would raise doubts into its provenance. Those few exceptions are the putative pseudogene cases, although we also note that several ncORFs incorporate transposon sequence, and it should be considered that additional sequences similar to these may exist outside our search space. Commentary on all alternative mappings is provided in Table S3 and also detailed below.

Following the arguments put forward in the main text, we emphasize that the decision of GENCODE *not* to annotate a given ncORF as protein-coding represents our position at the present time; we do not make the claim that such ORFs are definitively not protein-coding. Indeed, many of these cases are not straightforward from an evidence perspective, and an ORF could be reclassified in the future if more evidence becomes available. Furthermore, we reiterate that protein identification and protein-coding gene annotation are distinct processes, as considered in the Discussion.

Historically, GENCODE will annotate an ORF as protein-coding if it presents strong evidence that the translation is ancient in evolutionary terms and, more importantly, has clearly evolved under constraint at the protein-level as measured by PhyloCSF. This is because we believe that a confident observation along these lines provides evidence of protein function. Here, we anticipate that those ncORFs presenting obvious evolutionary signatures of this type have already been identified by our previous work, especially Mudge *et al*. Thus, for our present purposes, the main driver in annotation decision making is the quality of the peptide evidence.

As noted in the commentary in Table S3 and below, certain peptides are observed solely in cancer samples or cancer-derived cell lines, with this information being obtained via PeptideAtlas. We do not consider data of the latter kind to be ‘bad’ as such; indeed, such ‘non-normal’ peptide data may well validate the existence of the protein in those specific conditions. However, as discussed in the main text, GENCODE are at present holding back on annotating ncORFs that are supported only by cancer-derived peptides as protein-coding genes, where the translation also lacks evolutionary support for protein-level function.

The ncORF commentaries often make reference to 'ORF conservation'. To be clear, this concept considers the existence of the human ncORF in other species, with respect to the structure of the mammalian phylogenetic tree. We manually examine the presence of the initiation codon and termination codon in other genomes, as well as the maintenance of the reading frame, i.e. looking for indels leading to frame shifts or premature termination codons. We emphasise that this is at present a process of observation as opposed to measurement. Furthermore, it is important to note that the analysis of ncORF conservation is an entirely distinct process to the scoring of evolutionary constraint at the protein level as performed by PhyloCSF. Crucially, while deep conservation can be used to infer that a given ORF has a function, it does not - in isolation - directly inform as to the *mode* of function. It is now well established that translation does not simply act to generate proteins. In particular, there is now a well-defined paradigm for the mediation of protein expression via the translation of uORFs, and such 'regulatory ORFs' can be ancient. However, although regulatory ORFs may not be generally translated into bona fide proteins, it is plausible that they do generate unstable, peptide chains that are degraded by cellular proteases, with the potential to be detected by immunopeptidomics assays.

Certain ncORFs are described as '*de novo* emergences'. This is an established term, although one that can carry different shades of meaning. Here, *de novo* is used to reflect our view that the ncORF has evolved from genomic sequence that is ancestrally non-coding, and thus presumably non-translated at that point in evolutionary history. Thus, a ncORF that is described as a *de novo* emergence in human is one that does not have a conserved ORF counterpart in other species, although we do see that there is overall multigenome *sequence* alignment within a clade defined with respect to an ancient common ancestor.

In each case we provide a link for a multispecies alignment of the ncORF sequence produced with the CodAlignView tool, developed by I. Jungreis, M. Lin and M. Kellis at the Broad Institute (manuscript in preparation). CodAlignView uses existing multispecies genome alignments, and can be set to view Multiz alignments generated at UCSC, or 241-or 470-genome mammalian alignments generated using Cactus[2] as part of the Zoonomia project[3]. The link for each gene is set to the view we find to be most informative in terms of species incorporated, although this can be adjusted to any alignment made available for GRCh38.

### ORF cases

Here we provide commentary on each of the Tier 1A and Tier 2A ncORFs as appraised by PeptideAtlas and GENCODE, expanding upon information included in Table S3. The first commentary on c12nore105 is expanded to illustrate aspects of our manual annotation workflow that are also applicable to other ncORFs.

#### c12norep105

This ncORF is very well detected in the PeptideAtlas non-HLA build, with 7 distinct peptides. Of these, 6 have a PSM that was deemed “excellent” upon manual inspection. Only 1 peptide (AVDHGDAPLAAPPCAWALGPPLPR) was only deemed “good”. It is quite likely correct, but missing some coverage at the N terminus. This ncORF is highlighted in the main text and fully discussed there.

A key part of our efforts to elucidate ncORFs is to interpret their translation in the context of gene annotation, noting that the provenance of any translation event is substantially governed by the nature of the initial transcription event that creates the relevant RNA. This process is especially important in cases such as c12norep105, where gene annotation requires a mechanistic explanation as to how translation manages to occur in an alternative reading frame to the canonical CDS. Initially, we saw two main routes for coupling the transcription and translation of this ORF. First, converging transcriptomics data support the presence of an alternative transcript start site (TSS) in intron 2 of *CYP27B1*, which would produce a transcript within which the ATG used in c12norep105 is a plausible cognate initiation codon (existing transcript model ENST00000546567 would include the ORF in this context). In fact, there is an inframe upstream initiation codon on this model, found at chr12:57,765,524-57,765,526, that could be used to extend the ORF in the 5’ direction. Second, usage of an alternative splice acceptor site in exon 3 (as seen on model ENST00000713545, and well supported by RNA-seq data, not shown) introduces a frameshift in the *CYP27B1* CDS, moving the reading frame into that which is supported by the peptides.

However, when manually examining the source Ribo-seq data in more detail - looking at read coverage - we see evidence that c12norep105, as called in our original study, is 5’ truncated. Based on the MANE Select model ENST00000228606, the ORF remains open in the 5’ direction, extending to an inframe ATG found upstream of the canonical *CYP27B1* initiation codon, and we see good Ribo-seq support for this extension. If 5’ extended in this way, the c12norep105 intORF would be recontextualised as a ouORF, with a substantially larger translation of 347aa. It could in fact be that these three possibilities are not mutually exclusive, and that c12norep105 exists as a distinct ORF alongside the 5’ extended form, linked to alternative transcription events. However, in contrast to the initiation codon originally called for c12norep105, this upstream ATG has excellent support in Ribo-seq experiments treated with homoharringtonine (HHT), i.e it is experimentally supported as a true initiation codon.

Regardless of its transcriptional pathway, the functionality of this translation event — i.e. its contribution to normal cellular physiology, if any — remains hard to interpret. The conservation of the ORF - whether c12norep105 or the 5’ extended form - is limited to apes. As such, PhyloCSF inevitably produces a negative score, which means that this metric does not support a protein-level model of sequence evolution. Crucially, however, we observe that peptide PAp11480608 is derived from normal skin samples in a published study[4], indicating that this translation product is not aberrant or cancer specific. Meanwhile, peptide PAp11480608 was obtained from Ket-CT, which is an immortalised cell line derived from keratinocytes (the primary cell type in the outer skin). *CYP27B1* is known to be expressed in skin; it is the enzyme responsible for synthesizing the biologically active form of vitamin D. It is therefore interesting that the TSS found in intron two, with the potential to support the translation called as c12norep105, has its highest expression in keratinocytes; it must be noted though that there is also evidence of keratinocyte expression from the MANE Select TSS of *CYP27B1*.

In our view, annotation of additional coding sequence with the *CYP27B1* locus is justified by the peptide data. However, this is complicated by the outstanding questions as to the mechanism by which alternative translation occurs. GENCODE have annotated c12norep105 as protein-coding on model ENST00000546567, while the 347aa long form of the ORF is annotated as protein-coding on model ENST00000718428 (the latter is not yet present in the GENCODE public release at the time of publication). We have grouped these ORFs as part of *CYP27B1*, i.e. not as an independent gene, due to the possibility that the peptides could be validating an alternative protein isoform rather than a non-overlapping product. This is suggested by model ENST00000713545, which is also annotated as protein-coding.

The 347aa long form of the ncORF, using primate-only genomes in the Cactus alignment:

https://data.broadinstitute.org/compbio1/cav.php?controlsState=show&intervals=chr12%3A57%2C764%2C519-57%2C764%2C550%2Bchr12%3A57%2C764%2C754-57%2C764%2C926%2Bchr12%3A57%2C765%2C011-

[57%2C765%2C211%2Bchr12%3A57%2C765%2C297-57%2C765%2C499%2Bchr12%3A57%2C766%2C007-57%2C766%2C197%2Bchr12%3A57%2C766%2C847-57%2C767%2C090&strand=-&prologue=6&epilogue=6&alnset=hg38_241mammals_primate&wrap=50&spliceSites=human&hideInserts=on&hideJumps=on]

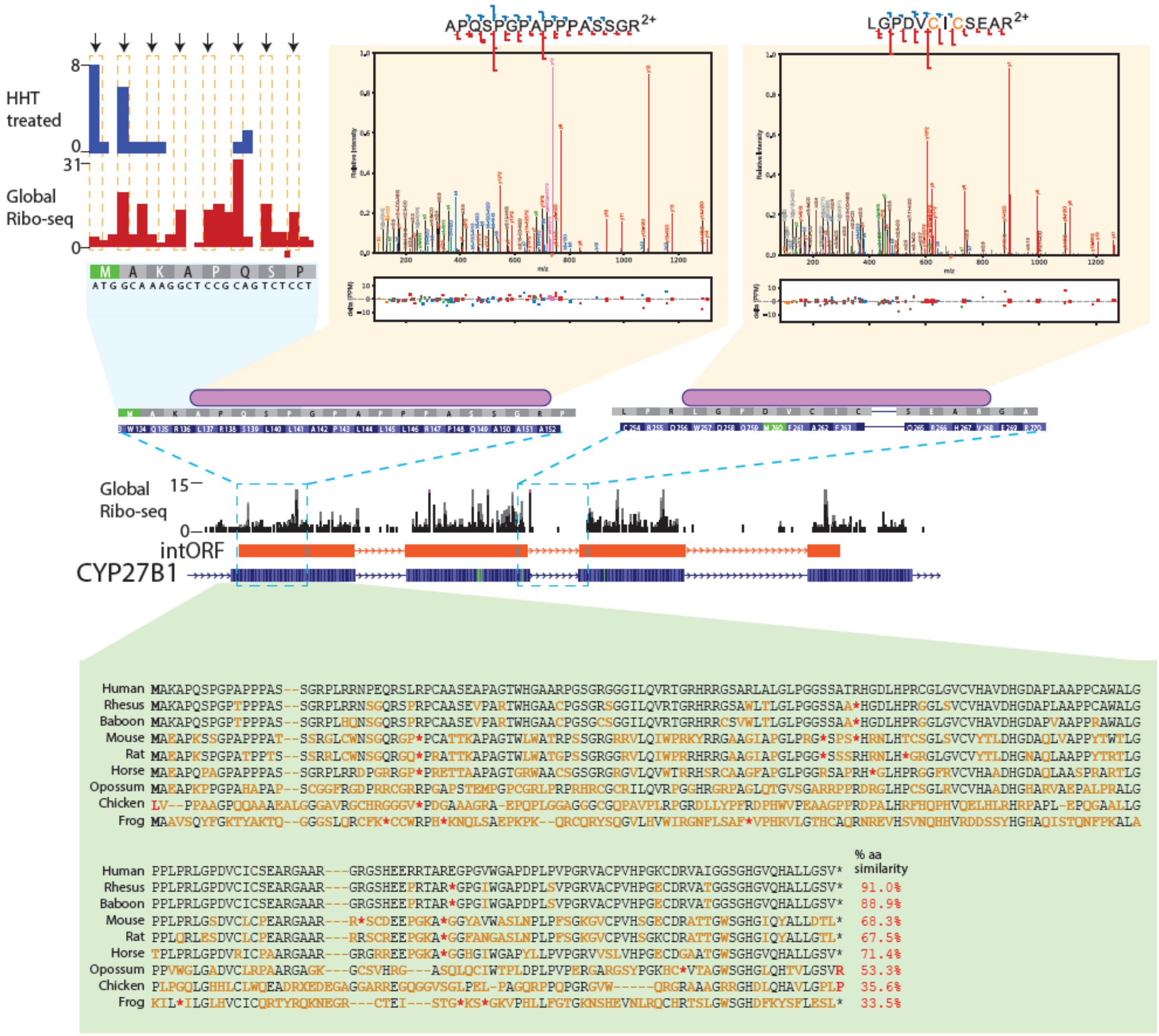


This figure includes the short form of the ORF, i.e. c12norep105, and indicates the lack of ORF conservation beyond apes. The top left panel indicates low-scoring though potentially discernable support for the initiation codon of this ORF in Ribo-seq experiments on HHT-treated cells. On the top right, spectra are shown for two high quality peptides.

#### c19riboseqorf4

c19riboseqorf4 is a uORF of the gene *STK11* found in the MANE Select transcript model ENST00000326873, and relatively large at 209aa. The 5' UTR region is ancestrally non-coding, making the ORF a *de novo* emergence. Most placental mammals have the ATG (mouse does not), but ORF conservation outside of higher primates is not consistent while PhyloCSF does not support a protein-level mode of evolution. GENCODE would not therefore annotate this ORF as protein-coding according to an evolutionary argument alone. The discovery of high quality peptide evidence is very interesting. This ncORF has 4 supporting peptides, all of which have best-PSMs deemed "excellent". For example: https://proteomecentral.proteomexchange.org/usi/?usi=mzspec:PXD020389:20171031_QE5_nLC3_AKP_UBIsite_SY5Y_CBLsKD_L-H_E1_17F_08:scan:11025:GGAFAPGTGGGTGASPETPPAK/2. It is detected in three separate datasets, PXD20389, PXD028647, and MSV000078509, all of which are derived from cancer cell line samples. In fact, each of the non-HLA peptides is derived from similar experiments, thus raising the possibility that it is an aberrant product as opposed to a functional protein molecule.

The ncORF in the 120-mammal Multiz alignment:

https://data.broadinstitute.org/compbio1/cav.php?controlsState=show&intervals=chr19%3A1%2C205%2C901-1%2C206%2C530&prologue=6&epilogue=6&alnset=hg38_120mammals&wrap=50&spliceSites=human&hideInserts=on&hideJumps=on

#### c14norep77

c14norep77 is an upstream overlapping ORF (uoORF) of *MAP3K9*, splicing across the first two exons of MANE Select model ENST00000554752. The ATG is found in all placental mammals, and the ORF is largely conserved as well; certain lineages have earlier STOP codons. As such, it looks promising as a functional element, i.e. according to the evolutionary argument. The 67bp portion of the ORF prior to the dual frame region is slightly PhyloCSF positive, after which the signal becomes harder to interpret. It has 5 supporting peptides, but 3 of the 5 are judged as not very good quality, primarily in the low-complexity poly-glycine region of the protein. For example, while the peptide GGGGGGGGGGGGGPR has some spectra that have multiple matching ions, it must be assumed that the true identification is elusive and not this sequence. These peptides are discounted in this analysis, but remain visible in the PeptideAtlas interface for inspection by the reader. This leaves 2 peptides with excellent spectra such as https://proteomecentral.proteomexchange.org/usi/?usi=mzspec:PXD020389:20171031_QE5_nLC3_AKP_UBIsite_SY5Y_CBLsKD_L-H_E1_17F_09:scan:9958:AAEPAGGHLPQQLR/2 that provide the required HPP-level evidence. Thus, GENCODE view c14norep77 as a potential protein, but it has not yet been annotated as such because the two informative peptides are only seen in cancer cell line experiments.

The ncORF in the 120-mammal Multiz alignment:

https://data.broadinstitute.org/compbio1/cav.php?controlsState=show&intervals=chr14%3A70%2C801%2C068-70%2C801%2C080%2Bchr14%3A70%2C808%2C766-70%2C809%2C238&strand=-&prologue=6&epilogue=6&alnset=hg38_120mammals&wrap=50&spliceSites=human&hideInserts=on&hideJumps=on

#### c14norep4

c14norep4 is a uORF of *ZNF219*. It maps to the MANE Select model; there are alternative first exons in the locus to which the ORF does not map. The ATG is found in all mammal genomes, although the length of the ORF varies substantially between species. There are 4 mapping peptides in PeptideAtlas, but the 7 amino-acid long peptide AAAAAAR is immediately discounted since it maps to many genes and so is not useful. This leaves 3 distinct peptides spread out over most of the length. Peptide AGPPPAAHNAGQGR has many matching ions but was deemed a false positive upon manual inspection. This leaves 2 peptides with excellent spectra that provide the HPP-level evidence. However, nearly all evidence comes from CPTAC cancer samples.

The ncORF in the 120-mammal Multiz alignment:

https://data.broadinstitute.org/compbio1/cav.php?controlsState=show&intervals=chr14%3A21%2C093%2C648-21%2C093%2C674%2Bchr14%3A21%2C098%2C312-21%2C098%2C527&strand=-

&prologue=6&epilogue=6&alnset=hg38_120mammals&wrap=50&spliceSites=human&hideInserts=on&hideJumps=on

#### c19norep7

c19norep7 is a dORF in the 3' UTR of *CIRBP*. The ATG is human specific, making the ORF a lineage specific *de novo* emergence. There are two uniquely-mapping tryptic peptides in PeptideAtlas, one with 7 PSMs and the other with only a single PSM. The PSMs mostly come from cancer cell lines, where only a single peptide is derived from a sample categorized as a non-cancer cell line. The best spectra for both peptides are quite convincing.

The ncORF using primate-only genomes in the Cactus alignment:

https://data.broadinstitute.org/compbio1/cav.php?controlsState=show&intervals=chr19%3A1%2C274%2C539-1%2C274%2C787&prologue=6&epilogue=6&alnset=hg38_241mammals_primate&wrap=50&spliceSites=human&hideInserts=on&hideJumps=on

#### c5norep4

c5norep4 is a dORF in the 3' UTR of *SLC9A3*. The ATG is conserved in apes and gibbon, but the ORF is human specific due to 1bp indel. The 3' UTR region is present in other mammals but not as an ORF, making this a de novo emergence. There are 4 uniquely-mapping tryptic peptides in PeptideAtlas, two of which have a high number of PSMs (43 and 17), although the other two are only single PSM peptides. The peptide with 43 PSMs has highly-convincing complete-coverage spectra associated with it. The other peptides are longer and do not have quite complete coverage but are quite likely correct. All PSMs come from 8 different cancer datasets from CPTAC enriched for phosphopeptides. It may be that this protein is normally of very low abundance, but brought into the detectable range via phosphopeptide enrichment.

The ncORF using primate-only genomes in the Cactus alignment:

https://data.broadinstitute.org/compbio1/cav.php?controlsState=show&intervals=chr5%3A472%2C072-472%2C413&strand=-&prologue=6&epilogue=6&alnset=hg38_241mammals_primate&wrap=50&spliceSites=human&hideInserts=on&hideJumps=on

#### c2riboseqorf90

c2riboseqorf90 is a dORF in the 3' UTR of *PTPN18*. The ORF is conserved in apes and gibbon, and the genomic alignment of the region in other mammals indicates that it is a de novo emergence. There are 3 uniquely-mapping tryptic peptides in PeptideAtlas. However, one is nested within another and thus does not count for HPP guidelines, so only 2 HPP-compliant peptides remain. One has 5 PSMs, some of which are extremely compelling with complete coverage, but the other only has a single PSM. This PSM has good coverage but suffers from contamination from another precursor in the selection window, and is thus not ideal. Most PSMs come from cancer samples. Only a single PSM comes from a deep pituitary gland sample.

The nccORF using primate-only genomes in the Cactus alignment:

https://data.broadinstitute.org/compbio1/cav.php?controlsState=show&intervals=chr2%3A130%2C373%2C716-130%2C373%2C880&prologue=6&epilogue=6&alnset=hg38_241mammals_primate&wrap=50&spliceSites=human&hideInserts=on&hideJumps=on

#### c1riboseqorf94

c1riboseqorf94 is a uORF of *CDKN2C*. The gene has an unusually large 5' UTR exon containing four translated ORFs in the Ribo-seq catalog, of which this is the longest at 180aa. Note that the ORF is not found on the MANE Select transcript ENST00000371761, which has a shortened 5' UTR. The ATG is conserved in mammals, though beyond apes there is inconsistency in the termination codon used. The ORF has its own uORF - c1norep114 - which overlaps on the STOP / ATG. There are 4 uniquely-mapping tryptic peptides in PeptideAtlas. Three have multiple PSMs, some of which are excellent. One peptide has only a single PSM that is severely contaminated by other precursors in the selection window, but may well be correct. All 61 PSMs come from CPTAC cancer samples or cancer cell lines.

The ncORF in the 120-mammal Multiz alignment:

https://data.broadinstitute.org/compbio1/cav.php?controlsState=show&intervals=chr1%3A50%2C968%2C788-50%2C969%2C330&prologue=6&epilogue=6&alnset=hg38_120mammals&wrap=50&spliceSites=human&hideInserts=on&hideJumps=on

#### c11riboseqorf4

c11riboseqorf4 is an overlapping uORF in *PIDD1*, which shares substantial sequence with a second ncORF in the catalog, c11norep6. The two ORFs use distinct splice acceptor sites with respect to the exon 2-3 splice junction of the *PIDD1* MANE Select model (ENST00000347755). This does not result in a frameshift, and so c11norep6 has an additional portion of interior sequence compared to the former. Ultimately, each peptide maps to both ORFs and so cannot distinguish between them in terms of evidence for translation. The evolutionary argument is not straightforward. The ATG is perfectly conserved in mammals, and almost entirely in reptile / avian genomes as well. The exon 2 portion of the ORF codes through in most mammals, with the exception of all rodent genomes, which have a premature STOP. The exon 3 portion is strikingly less well conserved as an ORF due to early STOPs in various lineages; the ORF is conserved as per its human length only in higher primates.

The alternative ATG of c11riboseqorf4 / c11norep6 is 31bp upstream of the canonical CDS. Essentially all transcripts produced by the gene contain both ATGs, and so it is not obvious how to distinguish between the two translational events on a transcriptional basis. The gene itself has a general expression profile.

There are 11 uniquely-mapping tryptic peptides in PeptideAtlas, with hundreds of observations. It is not listed as non-core canonical due to its similarity to c11norep6 and another predicted ncORF. But none of the 11 peptides map to UniProtKB or similar databases. Most of the PSMs are excellent and there is little doubt in this detection. Although there are substantial detections in cancer samples, there is also ample evidence from non-cancer samples. For the peptide SGLQGPSVGDGCNGGGAR at position 2 in the protein (methione at position 1 is never seen), all 15 PSMs are acetylated on the n terminus, as seen for example in this excellent PSM mzspec:PXD006633:HEK293T_N_term_F03:scan:7800:[Acetyl]-SGLQGPSVGDGC[Carbamidomethyl]N[Deamidated]GGGAR/2.

The ncORF in the 120-mammal Multiz alignment:

https://data.broadinstitute.org/compbio1/cav.php?controlsState=show&intervals=chr11%3A803%2C398-803%2C587%2Bchr11%3A804%2C094-804%2C419&strand=-&prologue=6&epilogue=6&alnset=hg38_120mammals&wrap=50&spliceSites=human&hideInserts=on&hideJumps=on

#### c16riboseqorf104

Although c16riboseqord104 has 3 peptides that appear to map uniquely to it, they have been deemed false positives and detection of this ncORF is not likely. All of the PSMs include the extended poly-alanine region, which is found in many other proteins, and therefore these PSMs should likely be mapped to one of those proteins, but either an amino-acid variation or mass modification prevents them from being identified since the correct answer is presumed not to be in our search space.

#### c6norep158

This case is not biologically informative. The peptides simply validate the canonical protein of *CASP8AP2*, which could not be properly annotated by GENCODE due to the presence of a substantial error on the GRCh38 reference genome (an entire canonical coding exon is missing). The original Ribo-seq call onto a processed transcript model was not incorrect per se, rather the pipeline did not recognise that the GENCODE annotation was compromised.

#### c19norep157

On detailed analysis, it was observed that the ncORF, found in the 3’ UTR of *ZNF607*, actually corresponds to a previously unrecognised retroinsertion of the prohibitin (*PHB1*)

mRNA. This process typically results in the formation of a processed pseudogene, but it can also create a functional ‘retrocopy’ of the parent locus. The observation of peptide evidence in this case is potentially interesting. However, the ORF is very similar to P35232, the UniProt entry for the *PHB1* parent gene, and we see only one unique mapping peptide with low observations. There are also at least a dozen other *PHB1* pseudogenes in the genome which are not in the PeptideAtlas search space, while pseudogenes were filtered out during the creation of the ncORF catalog. This specific retroinsertion happened in the primate order, at the base of the monkey radiation. While the ATG is intact in these species, the location of the termination codon is not consistent.

Given also that we see no transcriptional route for the expression of this ORF - i.e. other than being within the 3' UTR of *ZNF607* transcripts, which is a potentially poor translational context - GENCODE do not believe there is enough data to support protein-coding annotation at this time. Instead, the ORF has now been annotated as processed pseudogene ENSG00000289748.

The ncORF using primate-only genomes in the Cactus alignment:

https://data.broadinstitute.org/compbio1/cav.php?controlsState=show&intervals=chr19%3A37%2C696%2C932-37%2C697%2C402&strand=-&prologue=6&epilogue=6&alnset=hg38_241mammals_primate&wrap=50&spliceSites=human&hideInserts=on&hideJumps=on

#### c6norep53

c6norep53 is a dORF in the 3’ UTR of *BTN3A2*. Upon deeper analysis, it became apparent that the ORF is 'pseudoexon' sequence. In other words the C-t of the standard butyrophilin family protein is lost in this primate-specific copy, due to a premature termination codon in exon 7 compared with other full-length paralogs. So the dORF is a remnant of sequence that is coding in other copies, and the ncORF was able to be called during the creation of the catalog because this sequence has not been annotated by GENCODE (ncORFs overlapping known pseudogenes were filtered out). With this knowledge, it becomes clear that the peptides do not distinguish this ORF from O00478, the UniProt entry for *BTN3A2*.

The ncORF using primate-only genomes in the Cactus alignment:

https://data.broadinstitute.org/compbio1/cav.php?controlsState=show&intervals=chr6%3A26%2C376%2C596-26%2C377%2C276&prologue=6&epilogue=6&alnset=hg38_241mammals_primate&wrap=50&spliceSites=human&hideInserts=on&hideJumps=on

#### c21norep46

This former lncRNA ORF is now annotated as protein-coding gene *ERVH48-1* by GENCODE. This annotation decision was made in parallel to the work presented here, during a survey of UniProt / TrEMBL entries lacking corresponding GENCODE translations, and as part of a wider survey into retroelement sequences. The protein is understood to have placental function (there is transcript expression data supporting this profile), and has been considered a syncytin family member by Roberts et al.[5]. The peptide evidence therefore provides good additional support for the protein-coding annotation decision, especially as six of the peptides are derived from placenta experiments. The insertion happened at the base of the old world monkey / ape clade, with the ORF being intact in its human form in apes and gibbon. Interestingly, it is observed that the 160aa protein is not obviously an *env* ORF like other syncytins, and does not in fact appear to be *env* derived. The two exon gene is indeed embedded in a HERV, but the ORF looks more likely to have evolved *de novo* from sequence that was not contributed by the transposon insertion, while the splice junction of the 2-exon model (which has excellent transcriptomics support) also looks like an evolutionary innovation.

The ncORF using primate-only genomes in the 120-mammal Multiz alignment:

https://data.broadinstitute.org/compbio1/cav.php?controlsState=show&intervals=chr21%3A42%2C918%2C524-42%2C919%2C006&strand=-&prologue=6&epilogue=6&alnset=hg38_120mammals_primate&wrap=50&spliceSites=human&hideInserts=on&hideJumps=on

#### c11riboseqorf108

c11riboseqorf108 was called as a lncRNA ORF within *LIPT2-AS1.* On further analysis, it corresponds to an ORF contributed by a Tigger-family transposon. The insertion occurred at the base of the monkey / ape clade, and the transposase-like ORF is intact in these genomes. This could be evidence for a ‘domestication’ event, whereby a functional transposon ORF gains a new role in the host genome. However, all of the peptides found in support are from cancer samples or cell lines. For this reason, GENCODE have not yet decided to annotate this ORF as protein-coding. The CHESS gene annotation project already considers this locus a protein-coding gene with the same ORF, CHS.9489.1.

The ncORF using primate-only genomes in the Cactus alignment:

https://data.broadinstitute.org/compbio1/cav.php?controlsState=show&intervals=chr11%3A74%2C497%2C035-74%2C498%2C084&prologue=6&epilogue=6&alnset=hg38_241mammals_primate&wrap=50&spliceSites=human&hideInserts=on&hideJumps=on

#### c8riboseqorf15

c8riboseqorf15 is a uORF of *CNOT7*, found on the MANE Select transcript ENST00000361272.9. The ORF is perfectly conserved in all mammal genomes, with the exception of platypus. The peptides found in non-cancer samples or cell lines largely consist of the proline repeat sequence, and are not trustworthy to support reference annotation. Therefore GENCODE have not annotated this ORF as protein-coding.

The ncORF in the 120-mammal Multiz alignment:

https://data.broadinstitute.org/compbio1/cav.php?controlsState=show&intervals=chr8%3A17%2C245%2C236-17%2C245%2C247%2Bchr8%3A17%2C246%2C675-17%2C246%2C782&strand=-&prologue=6&epilogue=6&alnset=hg38_120mammals&wrap=50&spliceSites=human&hideInserts=on&hideJumps=on

#### c11norep11 / c11norep12

c11norep11 is a lncRNA ORF on *IGF2-AS,* sharing substantial sequence overlap with c11norep12. The two ORFs different in the usage of alternative splice donor sites leading into the second exon, which does not induce a frameshift; c11norep12 thus has a 37aa insertion. Both versions of the ORF are only fully conserved in ape and gibbon genomes, and look like a *de novo* emergence. UniProtKB already recognises c11norep12 as Q6U949. This protein was curated from Vu et al.[6], which predicts the ORF but does not provide experimental evidence for protein existence.

The peptides which support both translations - or rather cannot distinguish between them - are observed only in cancer samples or cell lines. Interestingly, the overexpression of the *IGF2-AS* transcript has been observed in certain tumours[6,7]. Peptide PAp00139202 spans the unique splice junction of c11norep12, and is observed in blood plasma.

The ncORF using primate-only genomes in the Cactus alignment:

https://data.broadinstitute.org/compbio1/cav.php?controlsState=show&intervals=chr11%3A2%2C140%2C644-2%2C140%2C947%2Bchr11%3A2%2C146%2C356-2%2C146%2C447&prologue=6&epilogue=6&alnset=hg38_241mammals_primate&wrap=50&spliceSites=human&hideInserts=on&hideJumps=on

#### c16riboseqorf59

c16riboseqorf59 is a dORF in the 3' UTR of *RNF40*, found in MANE Select model ENST00000324685.11. The ORF is a *de novo* emergence, found intact in chimp and gorilla only. There is an inframe upstream ATG at chr16:30,775,406-30,775,408 that lacks clear Ribo-seq support, while an inframe downstream ATG at chr16:30,775,583-30,775,585 appears to have substantially higher HHT support than for the ATG originally called. Usage of either of these alternative codons for initiation does not change the evolutionary picture.

PAp14808288 was observed in normal digestive tissue, specifically as part of an N-t enrichment assay designed to obtain initiation sites. However, this is linked to a third downstream ATG at chr16:30,775,781-30,775,783, which does not present with clear HHT data.

The ncORF using primate-only genomes in the Cactus alignment:

https://data.broadinstitute.org/compbio1/cav.php?controlsState=show&intervals=chr16%3A30%2C775%2C469-30%2C776%2C053&prologue=6&epilogue=6&alnset=hg38_241mammals_primate&wrap=50&spliceSites=human&hideInserts=on&hideJumps=on

#### c17riboseqorf149

C17riboseqorf149 is an uoORF in the 5' UTR of *TMEM94*, found on MANE Select transcript ENST00000314256. The ORF is intact in the vast majority of mammal genomes, with some localised movement of the termination codon in certain lineages. The excellent spectrum is from peptide PAp11219913, which is a cancer detection. Peptide PAp11219913 is observed in multiple normal tissues. However, as this peptide is not classed as excellent GENCODE have not annotated this ORF as protein-coding.

The ncORF using the Cactus alignment:

https://data.broadinstitute.org/compbio1/cav.php?controlsState=show&intervals=chr17%3A75%2C471%2C836-75%2C471%2C929%2Bchr17%3A75%2C485%2C428-75%2C485%2C547%2Bchr17%3A75%2C485%2C871-75%2C485%2C905&prologue=6&epilogue=6&alnset=hg38_241mammals&wrap=50&spliceSites=human&hideInserts=on&hideJumps=on

#### c19norep29

c19norep29 is a uoORF in the 5' UTR of *LONP1*, found in MANE Select ENST00000360614. The ATG is found in all therian mammals, and the ORF is largely conserved with the exception of certain lineages which use alternative termination codons. All of the peptide evidence is derived from cancer samples.

The ncORF in the Cactus alignment:

https://data.broadinstitute.org/compbio1/cav.php?controlsState=show&intervals=chr19%3A5%2C719%2C805-5%2C720%2C143&strand=-&prologue=6&epilogue=6&alnset=hg38_241mammals&wrap=50&spliceSites=human&hideInserts=on&hideJumps=on

#### c4riboseqorf52

c4riboseqorf52 was called as a uoORF in the 5’ UTR of *CCNI*, found within MANE Select model ENST00000237654. However, gene annotation suggests an alternative provenance for the experimental evidence. The ORF has two exons. The first is clearly an ancestral coding exon, and indeed GENCODE now annotate it as such (ENST00000507788) but only as a model that then skips MANE coding exon 1 and splices in downstream and continues in the CDS frame. In other words, the first exon of the ORF can be used to make an alternative isoform of *CCNI*. The second exon of the ORF is therefore what happens when the first coding exon of *CCNI* is not skipped, and it translates dual frame with respect to coding exon 1 before reaching a termination codon. Importantly, the peptides all map to the first exon of the ORF, which means that the simpler explanation is that they support the existence of the alternative *CCNI* isoform rather than the ORF as a protein product. Thus, c4riboseqorf52 has not been annotated as a protein-coding gene.

The ncORF in the Cactus alignment:

https://data.broadinstitute.org/compbio1/cav.php?controlsState=show&intervals=chr4%3A77%2C066%2C298-77%2C066%2C405%2Bchr4%3A77%2C075%2C472-77%2C075%2C543&strand=-&prologue=6&epilogue=6&alnset=hg38_241mammals&wrap=50&spliceSites=human&hideInserts=on&hideJumps=on

#### c9norep168

c9norep168 is called as a lncRNA ORF within *ARRDC1-AS*, and it has a substantial same frame overlap with c9riboseqorf120. The two ORFs have the same 3’ end, although c9riboseqorf120 is a single exon sequence whereas c9norep168 contains a distinct first exon as part of a two exon model. Q9H2J1 is a UniProt entry corresponding to c9riboseqorf120. This entry, apparently based on Zou et al.[8], does not provide any functional annotation for the protein. The lncRNA is limited to primates, and the ORF is intact only in apes and gibbon. The peptide evidence, which does not distinguish between the two ncORFs, is derived from cancer samples, hence GENCODE has not annotated either as protein-coding. Interestingly, Zou et al.[8] provide evidence for the expression of the lncRNA in cancer samples.

The ncORF in the primate-only Cactus alignment:

https://data.broadinstitute.org/compbio1/cav.php?controlsState=show&intervals=chr9%3A137%2C615%2C669-137%2C616%2C132%2Bchr9%3A137%2C617%2C388-137%2C617%2C463&strand=-&prologue=6&epilogue=6&alnset=hg38_241mammals_primate&wrap=50&spliceSites=human&hideInserts=on&hideJumps=on

#### c10norep59

c10norep59 is a lncRNA ORF in *LINC00839*. It overlaps in the same frame with c10norep60, which uses a different initiation codon and an intronic termination codon. Q8NAU0 is a TrEMBL entry matching c10norep59, while B4DDC5 is an alternative TrEMBL isoform from the same locus, distinct from both c10norep59 and c10norep60. The single candidate peptide thus matches to all four translations in the same location. This peptide is observed in a cancer sample.

The ncORF using primate-only genomes in the Cactus alignment:

https://data.broadinstitute.org/compbio1/cav.php?controlsState=show&intervals=chr10%3A42%2C475%2C573-42%2C475%2C880%2Bchr10%3A42%2C477%2C261-42%2C477%2C442%2Bchr10%3A42%2C486%2C979-42%2C487%2C055&prologue=6&epilogue=6&alnset=hg38_241mammals_primate&wrap=50&spliceSites=human&hideInserts=on&hideJumps=on

#### c10riboseqorf22

c10riboseqorf22 is a lncRNA ORF on *WAC-AS1*. This ORF used to have a corresponding Trembl entry D3DRW4, which was recently removed from the database. There are three other ncORFs in the gene, none of which overlap. The lncRNA itself is only present in higher primates, with the ORF intact only in ape and gibbon. The single excellent peptide is derived from a cancer sample.

The ncORF using primate-only genomes in the Cactus alignment:

https://data.broadinstitute.org/compbio1/cav.php?controlsState=show&intervals=chr10%3A28%2C523%2C873-28%2C524%2C190&strand=-&prologue=6&epilogue=6&alnset=hg38_241mammals_primate&wrap=50&spliceSites=human&hideInserts=on&hideJumps=on

#### c11norep16

c11norep16 is an doORF in *PHLDA2*, i.e. it initiates in an alternative frame to the CDS then translates downstream of the canonical termination codon and into the 3' UTR. The initiation codon of the ncORF overlaps with the *PHLDA2* termination codon as [ATGA], although - despite strong read coverage for the gene as a whole - no support for this initiation codon in HHT-treated samples is observed. In fact, following manual gene annotation an alternative, simpler explanation for the Ribo-seq signal and peptide support is apparent: usage of a cryptic splice donor site in exon 1 at chr11:2929166-2929168 causes a frameshift leading into exon 2, and switches the *PHLDA2* reading frame into that which is supported by the peptide evidence. Thus, it is likely that the Ribo-seq signal was originally miscalled as a doORF, when in reality it is derived from an alternative isoform of *PHLDA2*. This new coding sequence has been annotated by GENCODE as ENST00000718435.

#### c12norep197

c12norep197 is a uoORF in *MLXIP*, found in MANE Select model ENST00000319080. The ATG is found in all mammals (noting that some genome alignments are missing in Multiz / Cactus), and the ORF is almost entirely intact throughout the order with the exception of a few lineages. The supporting peptide is derived from a cancer sample.

The ncORF in the Cactus alignment:

https://data.broadinstitute.org/compbio1/cav.php?controlsState=show&intervals=chr12%3A122%2C078%2C831-122%2C079%2C253&prologue=6&epilogue=6&alnset=hg38_241mammals&wrap=50&spliceSites=human&hideInserts=on&hideJumps=on

#### c19norep18

c19norep18 is an intORF within the CDS of *NFIC*, found in MANE Select model ENST00000443272. The ATG of the ncORF was called immediately downstream of a splice acceptor site, and without HHT support for initiation, which suggests it may not have an accurately interpreted 5’ end. However, we do not observe a 5’ extension to the ORF that is convincingly supported by Ribo-seq or evolutionary analysis. The ATG of the ncORF is found in all mammals, although the termination codon shows substantial variability and the ORF is fully conserved only in higher primates. The supporting peptide is derived from a cancer sample.

The ncORF in the Cactus alignment:

https://data.broadinstitute.org/compbio1/cav.php?controlsState=show&intervals=chr19%3A3%2C381%2C713-3%2C381%2C964&prologue=6&epilogue=6&alnset=hg38_241mammals&wrap=50&spliceSites=human&hideInserts=on&hideJumps=on

#### c19riboseqorf160

c19riboseqorf160 is a uoORF in the 5’ UTR of *ZNF524*, found in MANE Select model ENST00000301073.4. The ATG is found in all mammals, although the termination codon shows positional variation and the ORF is only fully conserved in higher primates. The supporting peptide is derived from a cancer sample.

The ncORF in the Cactus alignment:

https://data.broadinstitute.org/compbio1/cav.php?controlsState=show&intervals=chr19%3A55%2C600%2C398-55%2C600%2C408%2Bchr19%3A55%2C602%2C075-55%2C602%2C378&prologue=6&epilogue=6&alnset=hg38_241mammals&wrap=50&spliceSites=human&hideInserts=on&hideJumps=on

#### c21norep26

c21norep26 was called as a dORF in the 3' UTR of *SON*. However, manual gene annotation makes it clear that the reading frame should instead be understood as the C-t of an alternative protein isoform of *SON*, which results from an alternative splicing reaction. This alternative isoform was recognised and annotated by GENCODE as ENST00000704334. The supporting peptide is observed in numerous normal tissues.

#### c2riboseqorf167

c2riboseqorf167 is an uoORF in the 5' UTR of *MARCHF4*, found in the MANE Select model ENST00000273067. The ATG initiation codon of the ORF overlaps with the STOP codon of c2norep257 found immediately upstream. The c2riboseqorf167 ORF is fully intact in almost all therian mammal genomes. The supporting peptide is derived from a cancer sample.

The ncORF in the Cactus alignment:

https://data.broadinstitute.org/compbio1/cav.php?controlsState=show&intervals=chr2%3A216%2C371%2C336-216%2C371%2C599&strand=-&prologue=6&epilogue=6&alnset=hg38_241mammals&wrap=50&spliceSites=human&hideInserts=on&hideJumps=on

#### c9riboseqorf46

c9riboseqorf46 is a lncRNA ORF in *UBQLN1-AS1*. The entire ORF is contributed by a TcMar-Mariner transposon, giving a transposase-like putative protein. The insertion happened prior to the radiation of higher primates, although the ORF is only intact in apes and gibbon. As for c11riboseqorf8, this could be a ‘domestication’ event, whereby a functional transposon ORF gains a new role in the host genome. This ORF corresponds to B7Z6N6, an unreviewed TrEMBL entry. However, on further inspection, the peptides also map to Q53H47, the UniProt entry for known protein-coding gene *SETMAR*. Thus, GENCODE will not annotate this ORF as protein-coding until locus-specifc peptide evidence becomes available.

The ncORF using primate-only genomes in the Cactus alignment:

https://data.broadinstitute.org/compbio1/cav.php?controlsState=show&intervals=chr9%3A83%2C712%2C501-83%2C712%2C995&prologue=6&epilogue=6&alnset=hg38_241mammals&wrap=50&spliceSites=human&hideInserts=on&hideJumps=on

#### c11norep103

C11norep103 is a uORF in *MAP3K11*; it was previously called as a ncORF on a processed transcript, as the model used for initial interpretation was annotated as a retained intron model (ENST00000524856.1). The ORF is conserved in higher primates, and represents a *de novo* emergence. The supporting peptide is observed only in cancer cell lines.

The ncORF using primate-only genomes in the Cactus alignment:

https://data.broadinstitute.org/compbio1/cav.php?controlsState=show&intervals=chr11%3A65%2C614%2C947-65%2C615%2C156&strand=-&prologue=6&epilogue=6&alnset=hg38_470mammals_primate&wrap=50&spliceSites=human&hideInserts=on&hideJumps=on

#### c11norep190

C11norep190 is a dORF in *H2AX*. The ATG is present in higher primate genomes, with the ORF conserved only in apes and gibbon. The supporting peptide has been identified in frontal cortex; the gene itself is expressed in all tissues, based on transcriptomics data.

The ncORF using primate-only genomes in the Cactus alignment:

https://data.broadinstitute.org/compbio1/cav.php?controlsState=show&intervals=chr11%3A119%2C094%2C706-119%2C094%2C885&strand=-&prologue=6&epilogue=6&alnset=hg38_241mammals_primate&wrap=50&spliceSites=human&hideInserts=on&hideJumps=on

#### c11norep22

c11norep22 is an ouORF in *TUB*. The ATG and ORF is conserved in placental mammal genomes. The supporting peptide includes an observation in non-cancer cell lines.

The ncORF in the Cactus alignment:

https://data.broadinstitute.org/compbio1/cav.php?controlsState=show&intervals=chr11%3A8%2C081%2C411-8%2C081%2C548%2Bchr11%3A8%2C089%2C610-8%2C089%2C621&prologue=6&epilogue=6&alnset=hg38_470mammals&wrap=50&spliceSites=human&hideInserts=on&hideJumps=on

#### c11norep89

c11norep89 is a lncRNA ORF in PPP1R14B-AS1. The ATG and ORF are conserved in the clade defined by apes, old world monkeys and new world monkeys. The supporting peptide is observed only in cancer samples.

The ncORF using primate-only genomes in the Cactus alignment:

https://data.broadinstitute.org/compbio1/cav.php?controlsState=show&intervals=chr11%3A64%2C247%2C221-64%2C247%2C259%2Bchr11%3A64%2C247%2C953-64%2C248%2C063&prologue=6&epilogue=6&alnset=hg38_241mammals_primate&wrap=50&spliceSites=human&hideInserts=on&hideJumps=on

#### c12norep33

c12norep33 is a uORF in *PRH1*, although these 5' UTR exons are shared with *PRR4*. The ATG and ORF are conserved in higher primates. The peptide has been observed in normal tissues. F1T0A8 is an unreviewed TrEMBL entry corresponding to the same ORF.

The ncORF using primate-only genomes in the Cactus alignment:

https://data.broadinstitute.org/compbio1/cav.php?controlsState=show&intervals=chr12%3A11%2C121%2C140-11%2C121%2C180%2Bchr12%3A11%2C171%2C422-11%2C171%2C599&strand=-&prologue=6&epilogue=6&alnset=hg38_241mammals_primate&wrap=50&spliceSites=human&hideInserts=on&hideJumps=on

#### c12norep97

c12norep97 is a dORF in *RBMS2*. The ATG and ORF are conserved only in apes and gibbon. The peptide has an observation in normal tissues. The dORF is potentially reinterpreted an alternative final coding exon of *RBMS2*, as skipping of the penultimate exon will bring the annotated CDS into the correct frame. There is transcriptomics support for this, so a new GENCODE model can be annotated (RefSeq already have a corresponding model for this, prediction XM_005269060.6). Ribo-seq data does not show HHT support for the dORF ATG, but it does not provide unequivocal read coverage for the novel isform either. The dORF may yet be correct, or potentially both the dORF and novel isoform are translated.

The ncORF using primate-only genomes in the Cactus alignment:

https://data.broadinstitute.org/compbio1/cav.php?controlsState=show&intervals=chr12%3A56%2C589%2C190-56%2C589%2C261&prologue=6&epilogue=6&alnset=hg38_241mammals_primate&wrap=50&spliceSites=human&hideInserts=on&hideJumps=on

#### c13riboseqorf32

c13riboseqorf32 is a uORF in *CAB39L*. The ATG and ORF are almost entirely conserved in primates genomes; conservation in other mammal genomes is harder to interpret, with the ORF disrupted in several lineages. The single supporting peptide is non-tryptic, and only observed in cancer cell lines.

The ncORF using primate-only genomes in the Cactus alignment:

https://data.broadinstitute.org/compbio1/cav.php?controlsState=show&intervals=chr13%3A49%2C433%2C382-49%2C433%2C393%2Bchr13%3A49%2C434%2C086-49%2C434%2C151&strand=-&prologue=6&epilogue=6&alnset=hg38_470mammals&wrap=50&spliceSites=human&hideInserts=on&hideJumps=on

#### c14riboseqorf98

c14riboseqorf98 is an ouORF in *FOXN3*. The ORF is not resolved in Multiz alignments due to repetitive sequence, but Cactus shows that the ATG and ORF are well conserved in mammals, with some localised movement of the termination codon. The single peptide is only observed in cancer samples.

The ncORF in the Cactus alignment:

https://data.broadinstitute.org/compbio1/cav.php?controlsState=show&intervals=chr14%3A89%2C412%2C451-89%2C412%2C490%2Bchr14%3A89%2C416%2C871-89%2C417%2C070&strand=-&prologue=6&epilogue=6&alnset=hg38_470mammals&wrap=50&spliceSites=human&hideInserts=on&hideJumps=on

#### c15riboseqorf103

c15riboseqorf103 is a uORF in *CHSY1*. The ATG and ORF are conserved in mammalian genomes, with few exceptions. The single peptide has an observation in placenta; the gene is very highly expressed in this tissue based on RNAseq and CAGE data. The ORF is 124aa, so if protein-coding it would not be classed as a microprotein.

The ncORF in the Cactus alignment:

https://data.broadinstitute.org/compbio1/cav.php?controlsState=show&intervals=chr15%3A101%2C251%2C543-101%2C251%2C914&strand=-&prologue=6&epilogue=6&alnset=hg38_470mammals&wrap=50&spliceSites=human&hideInserts=on&hideJumps=on

#### c17norep1

c17norep1 is a lncRNA ORF in RPH3AL-AS1, conserved only in the human genome. The single peptide is observed only in cancer samples.

The ncORF using primate-only genomes in the Cactus alignment:

https://data.broadinstitute.org/compbio1/cav.php?controlsState=show&intervals=chr17%3A331%2C275-331%2C598&prologue=6&epilogue=6&alnset=hg38_470mammals_primate&wrap=50&spliceSites=human&hideInserts=on&hideJumps=on

#### c18riboseqorf10

c18riboseqorf10 is a lncRNA ORF in *LINC00526*, conserved only in apes. The single peptide is observed only in cancer samples. UniProt annotate this precise ORF as Q96FQ7, although it is classified by them as PE5 (‘product of a dubious gene prediction’) so is not taken forward by HUPO-HPP.

The ncORF using primate-only genomes in the Cactus alignment:

https://data.broadinstitute.org/compbio1/cav.php?controlsState=show&intervals=chr18%3A5%2C237%2C364-5%2C237%2C651&strand=-&prologue=6&epilogue=6&alnset=hg38_470mammals_primate&wrap=50&spliceSites=human&hideInserts=on&hideJumps=on

#### c19norep108

c19norep108 is a lncRNA ORF in ENSG00000266976. The gene is entirely contained within a HERV retrotransposon element, an insertion that seems to have occurred at the base of the clade defined by apes and old world monkeys. The ATG and ORF are conserved only in apes. The single supporting peptide is observed only in cancer cell lines. Q6ZNE2 is an unreviewed TrEMBL entry corresponding to the same ORF.

The ncORF using primate-only genomes in the Cactus alignment:

https://data.broadinstitute.org/compbio1/cav.php?controlsState=show&intervals=chr19%3A28%2C606%2C956-28%2C607%2C104%2Bchr19%3A28%2C610%2C260-28%2C610%2C331%2Bchr19%3A28%2C610%2C982-28%2C611%2C309&prologue=6&epilogue=6&alnset=hg38_470mammals_primate&wrap=50&spliceSites=human&hideInserts=on&hideJumps=on

#### c19norep8

c19norep8 is an overlapping dORF in *C19orf24* / *FAM174C*. Conservation of the ATG and ORF is limited to higher primates. The single peptide is observed in a non-cancerous cell line.

The ncORF using primate-only genomes in the Cactus alignment:

https://data.broadinstitute.org/compbio1/cav.php?controlsState=show&intervals=chr19%3A1%2C277%2C297-1%2C277%2C299%2Bchr19%3A1%2C278%2C777-1%2C279%2C073&prologue=6&epilogue=6&alnset=hg38_470mammals_primate&wrap=50&spliceSites=human&hideInserts=on&hideJumps=on

#### c19riboseqorf180

c19riboseqorf180 is a dORF in *A1BG*. The ATG and ORF are conserved in higher primates. The single peptide is observed in blood plasma, which is interesting because A1BG is a glycoprotein secreted into plasma. The ORF as called by Ribo-seq is potentially translated from a two exon model based on a strong TSS found at approximately chr19:58347673, i.e. as seen on current GENCODE model ENST00000598345. Alternatively, the usage of an alternative splice donor site in the penultimate exon of the *A1BG* mRNA at chr19:58347427 would bring the *A1BG* CDS into the same frame as the dORF, and in this case the peptide could be explained as supporting a new protein isoform. This alternative splice junction has substantial transcriptomics support, but is nonetheless a minor transcript in comparison to MANE Select model ENST00000263100. These two explanations may not be mutually exclusive. Ultimately, the locus is intriguing, though hard to comprehensively appraise without further data.

The ncORF using primate-only genomes in the Cactus alignment:

https://data.broadinstitute.org/compbio1/cav.php?controlsState=show&intervals=chr19%3A58%2C346%2C882-58%2C347%2C022&strand=-&prologue=6&epilogue=6&alnset=hg38_470mammals_primate&wrap=50&spliceSites=human&hideInserts=on&hideJumps=on

#### c19riboseqorf183

c19riboseqorf183 is a lncRNA ORF in ENSG00000232098. The ATG and ORF are conserved in apes. The single peptide is observed in non-cancer cell lines.

The ncORF using primate-only genomes in the Cactus alignment:

https://data.broadinstitute.org/compbio1/cav.php?controlsState=show&intervals=chr19%3A58%2C406%2C715-58%2C406%2C770%2Bchr19%3A58%2C408%2C127-58%2C408%2C343&strand=-&prologue=6&epilogue=6&alnset=hg38_470mammals_primate&wrap=50&spliceSites=human&hideInserts=on&hideJumps=on

#### c1riboseqorf33

c1riboseqorf33 is lncRNA ORF in *EMC1-AS1*, found directly antisense to CDS of *EMC1*. The ATG and ORF are conserved in higher primates, with termination codon movement in some lineages. The single supporting peptide is observed in cancer samples.

The ncORF using primate-only genomes in the Cactus alignment:

https://data.broadinstitute.org/compbio1/cav.php?controlsState=show&intervals=chr1%3A19%2C240%2C242-19%2C240%2C453&prologue=6&epilogue=6&alnset=hg38_470mammals_primate&wrap=50&spliceSites=human&hideInserts=on&hideJumps=on

#### c20riboseqorf22

c20riboseqorf22 ouORF in *KIF16B*. The ATG and ORF are conserved in mammals with some localised movement of the termination codon. The single peptide is observed in cancer cell lines.

The ncORF in the Cactus alignment:

https://data.broadinstitute.org/compbio1/cav.php?controlsState=show&intervals=chr20%3A16%2C526%2C147-16%2C526%2C205%2Bchr20%3A16%2C528%2C371-16%2C528%2C440%2Bchr20%3A16%2C573%2C229-16%2C573%2C282&strand=-&prologue=6&epilogue=6&alnset=hg38_470mammals&wrap=50&spliceSites=human&hideInserts=on&hideJumps=on

#### c20riboseqorf61

c20riboseqorf61 is a lncRNA ORF in *MHENCR*. The ATG and ORF are human specific. The single peptide is observed only in cancer samples and cell lines.

The ncORF using primate-only genomes in the Cactus alignment:

https://data.broadinstitute.org/compbio1/cav.php?controlsState=show&intervals=chr20%3A63%2C627%2C323-63%2C627%2C392%2Bchr20%3A63%2C628%2C190-63%2C628%2C362&prologue=6&epilogue=6&alnset=hg38_470mammals&wrap=50&spliceSites=human&hideInserts=on&hideJumps=on

#### c2norep161

c2norep161 is a lncRNA ORF in *LCT-AS1*. The ATG and ORF are specific to apes. The single peptide is observed only in cancer cell lines.

The ncORF using primate-only genomes in the Cactus alignment:

https://data.broadinstitute.org/compbio1/cav.php?controlsState=show&intervals=chr2%3A135%2C820%2C546-135%2C820%2C639%2Bchr2%3A135%2C821%2C675-135%2C821%2C871&prologue=6&epilogue=6&alnset=hg38_470mammals&wrap=50&spliceSites=human&hideInserts=on&hideJumps=on

#### c2norep170

c2norep170 is an internal ORF in *FMNL2*. The peptide also maps to RefSeq model XP_005246322.1, an alternative isoform of *FMNL2* not yet represented in GENCODE. This model includes an additional cassette exon in the final intron, with coordinates chr2:152645434-152645480; when spliced in it introduces a frameshift into the *FMNL2* translation that terminates in an alternative reading frame. The peptide supports this new C-terminus. The cassette exon is conserved in vertebrates, and highly transcribed. Thus, it is considered that this a superior explanation for the peptide evidence. The peptide itself is found in multiple normal tissues.

#### c2riboseqorf145

c2riboseqorf145 is a uORF in *PLCL1*. The ATG and ORF are conserved in mammal genomes, with very few exceptions. The ORF is highly alanine-rich, in a manner that is linked to length variation in different species. The single peptide is observed in cancer cell lines only.

The ncORF in the Cactus alignment:

https://data.broadinstitute.org/compbio1/cav.php?controlsState=show&intervals=chr2%3A197%2C804%2C922-197%2C804%2C987&prologue=6&epilogue=6&alnset=hg38_470mammals&wrap=50&spliceSites=human&hideInserts=on&hideJumps=on

#### c3riboseqorf132

c3riboseqorf132 is a lncRNA ORF in *TMCC1-DT*, located entirely within a HERV retrotransposon which inserted in higher primates. Non-ape primates have premature termination codons. The single unique-mapping peptide is observed in blood cells.

The ncORF using primate-only genomes in the Cactus alignment:

https://data.broadinstitute.org/compbio1/cav.php?controlsState=show&intervals=chr3%3A129%2C905%2C678-129%2C906%2C013&prologue=6&epilogue=6&alnset=hg38_470mammals&wrap=50&spliceSites=human&hideInserts=on&hideJumps=on

#### c3riboseqorf16

c3riboseqorf16 is a lncRNA ORF in *FGD5-AS1*, located entirely with a TcMar-Mariner family DNA transposon; see also comments for c9riboseqorf46. It overlaps with c3norep21 and c3norep22 in alternative reading frames. The transposon inserted in higher primates, and the ORF is only conserved in apes. The single unique-mapping peptide is observed in non-cancer cell lines. Other peptides cannot distinguish this ORF from e.g. *SETMAR* protein.

The ncORF using primate-only genomes in the Cactus alignment:

https://data.broadinstitute.org/compbio1/cav.php?controlsState=show&intervals=chr3%3A14%2C945%2C419-14%2C945%2C598&strand=-&prologue=6&epilogue=6&alnset=hg38_470mammals&wrap=50&spliceSites=human&hideInserts=on&hideJumps=on

#### c3riboseqorf166

c3riboseqorf166 is a uoORF in *FNDC3B*, with the start codon immediately upstream of the CDS. The ATG is conserved in vertebrate genomes, with the position of the termination codon showing variability. The peptide is noted as multimapping by PeptideAtlas because it also maps to smORFs within the locus published by Cui *et al*. It is observed in cancer samples only.

The ncORF in the 100 tetrapod alignment:

https://data.broadinstitute.org/compbio1/cav.php?controlsState=show&intervals=chr3%3A172%2C112%2C476-172%2C112%2C590%2Bchr3%3A172%2C133%2C471-172%2C133%2C490&prologue=6&epilogue=6&alnset=hg38_100_tetrapod&wrap=50&spliceSites=human&hideInserts=on&hideJumps=on

#### c4riboseqorf34

c4riboseqorf34 is a uORF in *OCIAD1*. The ATG and ORF are conserved only in chimp and gorilla genomes. The supporting peptide includes observations in normal tissues. The peptide maps also to RefSeq models XP_024309873.1,XP_024309877.1 within the locus, both of which contain an alternative N-t that represents the first portion of the Ribo-seq ORF before a splice junction leads the translation into the same frame as the canonical CDS. The peptide maps to this shared sequence and so it is plausible that it validates this putative isoform as opposed to the ORF. These possibilities are not mutually exclusive however, and the Ribo-seq data does provide support for the uORF as an intact translated sequence, i.e. as originally called.

The ncORF using primate-only genomes in the Cactus alignment:

https://data.broadinstitute.org/compbio1/cav.php?controlsState=show&intervals=chr4%3A48%2C831%2C102-48%2C831%2C206&prologue=6&epilogue=6&alnset=hg38_470mammals_primate&wrap=50&spliceSites=human&hideInserts=on&hideJumps=on

#### c5norep142

c5norep142 is an intORF in *MATR3*, discussed in the main text on the basis of its immunopeptidomics support. The single non-HLA peptide is observed in cancer cell lines. The peptide also maps to smORFs called within the locus by Cui *et al*.

#### c5riboseqorf2

c5riboseqorf2 was called as a lncRNA ORF in *SLC9A3-OT1*, but it is also a dORF in *SLC9A3*; the two genes overlap on the same strand. The ATG and ORF are specific to the human genome, being a *de novo* emergence. The ORF is identical to unreviewed TrEMBL entry Q71RB1. One supporting peptide has been observed in normal tissues.

The ncORF using primate-only genomes in the Cactus alignment:
https://data.broadinstitute.org/compbio1/cav.php?controlsState=show&intervals=chr5%3A471%2C793-471%2C823%2Bchr5%3A472%2C679-472%2C935&strand=-&prologue=6&epilogue=6&alnset=hg38_470mammals_primate&wrap=50&spliceSites=human&hideInserts=on&hideJumps=on

#### c6norep103

c6norep103 is an uoORF in *DAXX*, sharing its first of two coding exons with c6riboseqorf53 and c6riboseqorf54, which are uORFs. This first coding exon can potentially be used to produce an alternative DAXX protein isoform via a splice site shift, as seen in model ENST00000706094. While the ATG and ORF are observed in most mammal genomes, several lineages are missing the former and others have alternative termination codons. The supporting peptide is observed in cancer samples.

The ncORF in the Cactus alignment:

https://data.broadinstitute.org/compbio1/cav.php?controlsState=show&intervals=chr6%3A33%2C321%2C345-33%2C321%2C567%2Bchr6%3A33%2C322%2C862-33%2C322%2C914&strand=-&prologue=6&epilogue=6&alnset=hg38_470mammals&wrap=50&spliceSites=human&hideInserts=on&hideJumps=on

#### c6norep174

c6norep174 is a dORF in *HDAC2*. The ATG is observed in most primate species, although the termination codon is unique to human. The supporting peptide is observed only in cancer cell lines.

The ncORF using primate-only genomes in the Cactus alignment:

https://data.broadinstitute.org/compbio1/cav.php?controlsState=show&intervals=chr6%3A113%2C939%2C147-113%2C939%2C239&strand=-&prologue=6&epilogue=6&alnset=hg38_470mammals_primate&wrap=50&spliceSites=human&hideInserts=on&hideJumps=on

#### c6riboseqorf17

c6riboseqorf17 is an uoORF in *MRS2*. The ATG and ORF are conserved in mammal genomes, although with some localised termination codon movement. The supporting peptide is observed only in cancer cell lines.

The ncORF in the Cactus alignment:

https://data.broadinstitute.org/compbio1/cav.php?controlsState=show&intervals=chr6%3A24%2C403%2C001-24%2C403%2C236%2Bchr6%3A24%2C405%2C168-24%2C405%2C171&prologue=6&epilogue=6&alnset=hg38_470mammals&wrap=50&spliceSites=human&hideInserts=on&hideJumps=on

#### c7norep19

c7norep19 is a dORF in *ACTB*. The evolutionary history is complex. The region as a whole is conserved in vertebrates, strikingly so across several sequences. In terms of phylogenetics, the ORF seems most likely to be a mammalian *de novo* emergence that has been lost in several lineages, although length variation across a simple repeat region complicates the picture. The single peptide is observed in cancer cell lines.

The ncORF in the Cactus alignment:

https://data.broadinstitute.org/compbio1/cav.php?controlsState=show&intervals=chr7%3A5%2C527%2C342-5%2C527%2C683&strand=-&prologue=6&epilogue=6&alnset=hg38_470mammals&wrap=50&spliceSites=human&hideInserts=on&hideJumps=on

#### c8norep15

c8norep15 is a uoORF in *LZTS1*. The ATG is conserved in mammals with the exception of rodents, and also in reptiles and avian species. The ORF is conserved only in higher primates due to indels and substantial variation in the position of the termination codon. The peptide is observed only in cancer samples.

The ncORF in the Cactus alignment:

https://data.broadinstitute.org/compbio1/cav.php?controlsState=show&intervals=chr8%3A20%2C255%2C055-20%2C255%2C279&strand=-&prologue=6&epilogue=6&alnset=hg38_470mammals&wrap=50&spliceSites=human&hideInserts=on&hideJumps=on

#### c8riboseqorf118

c8riboseqorf118 is a uORF in *MAF1*. The ATG and ORF are conserved in mammalian genomes. The single peptide is observed in urine, while *MAF1* has general expression.

The ncORF in the Cactus alignment:

https://data.broadinstitute.org/compbio1/cav.php?controlsState=show&intervals=chr8%3A144%2C104%2C625-144%2C104%2C747&prologue=6&epilogue=6&alnset=hg38_470mammals&wrap=50&spliceSites=human&hideInserts=on&hideJumps=on

#### c9riboseqorf74

c9riboseqorf74 is an uoORF in *PHF19*. The ATG is conserved in mammals, with the ORF conserved only in primates. The peptide is observed only in cancer cell lines.

The ncORF in the Cactus alignment:

https://data.broadinstitute.org/compbio1/cav.php?controlsState=show&intervals=chr9%3A120%2C874%2C047-120%2C874%2C060%2Bchr9%3A120%2C874%2C556-120%2C874%2C756%2Bchr9%3A120%2C877%2C091-120%2C877%2C142&strand=-&prologue=6&epilogue=6&alnset=hg38_470mammals&wrap=50&spliceSites=human&hideInserts=on&hideJumps=on

#### cXnorep77

cXnorep77 is a doORF in *NUP62CL*. However, it appears that the ncORF represents a truncation of an alternative protein isoform of *NUP62CL* that was not annotated when the set was constructed; splicing in a cassette exon at chrX:107,150,856-107,150,967 induces a frameshift in the *NUP62CL* CDS which puts the translation into the same frame as the ncORF. Given the absence of HHT data to support the internal initiation codon called for the ncORF, it seems most likely that the peptide supports instead this alternative protein isoform, which is represented by ENST00000432145. The peptide is observed only in cancer cell lines.
