## Supplemental Document S2 for "High-quality peptide evidence for annotating non-canonical open reading frames as human proteins"

### *Supplementary Document S2: Discussion of machine learning results predicting detectability of ncORFs.*

To better understand and estimate the differences between non-coding ORFs (ncORFs) that we detect and those that we do not detect and the differences between HLA-I peptides that we detect and those that we do not detect by Mass Spectrometry (MS), we have trained two models based on the properties of the detected and undetected ncORFs and HLA-I peptides. Inferences from these model results do not necessarily represent causality, yet they estimate how several amino acid sequence-based features may influence detectability by MS. Section 1 below describes the ncORF classification model, and Section 2 describes the HLA-I peptide classification model.

#### 1. ncORFs microproteins classification model

To discern potential distinctions between detected and undetected ncORF microproteins, we curated a dataset encompassing both categories and applied a Statistical Learning model for analysis.

Our analysis focused on a cohort of 7264 ncORFs, comprising 1785 that were detected and 5479 that remained undetected. Utilising the amino acid sequences corresponding to all ncORF microproteins, we computed a comprehensive set of 35 attributes. This process involved leveraging the Bio.SeqUtils.ProtParam module from BioPython and the Amino Acid Indices version 9.2 (https://www.genome.jp/). Subsequently, we employed the Boruta algorithm (https://gitlab.com/mbq/Boruta/) to select the most relevant features from the pool of 35 attributes.

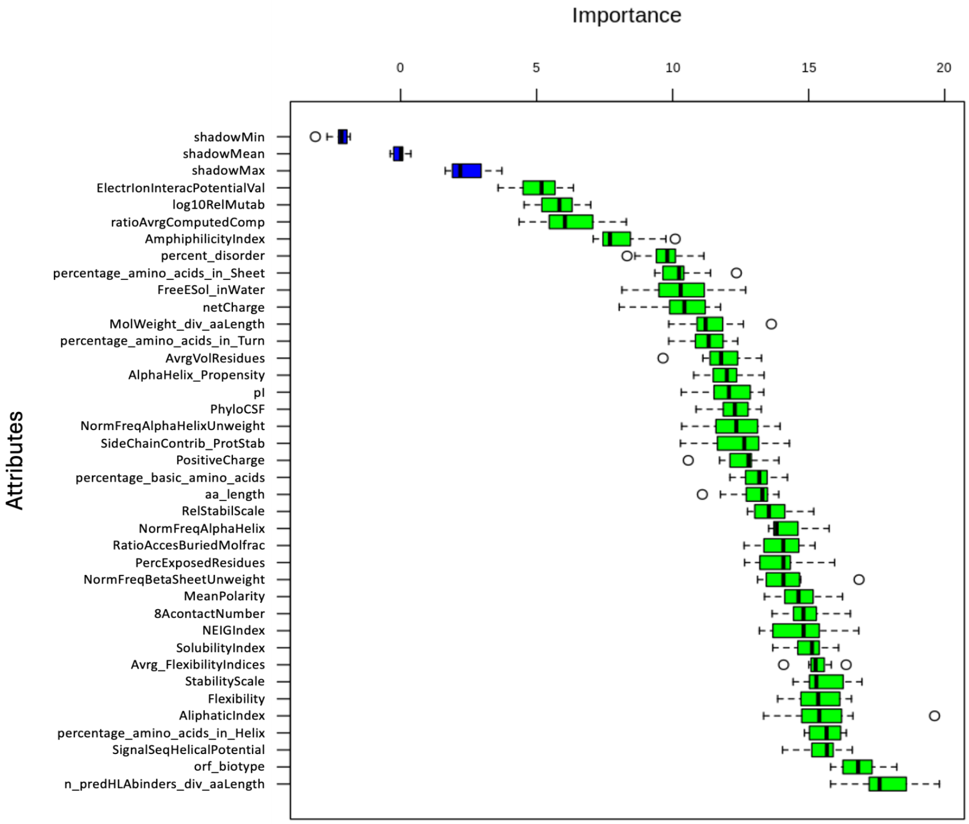


**Figure 1.** Boruta selected attributes. The Boruta algorithm tries to capture all the important features with respect to the outcome variable represented by ncORF peptide detectability. The Z-score, also known as the standard score, is a statistical measurement that describes a value's relationship to the mean of a group of values. The Boruta algorithm iteratively assesses the importance of each feature by comparing the Z-scores of actual features against those of randomly permuted shadow features. Features that consistently have higher Z-scores than the shadow features are considered important, while those with lower Z-scores are considered unimportant and are removed from the model. Blue box plots correspond to minimal, average and maximum Z score of a shadow attribute. Red and green box plots represent Z scores of respectively rejected and confirmed, and yellow boxplots are considered tentative attributes.

Boruta confirmed 36 attributes (**Figure 1**, green) and no attributes were deemed unimportant. Since orf_biotype is a 'categorical feature' with 7 different types, we replaced orf_biotype with 7 features representing each category. We added the Instability Index attribute, and altogether, we used 43 attributes to test a cost-sensitive binary classification TensorFlow-Keras model.

1. ElectrIonInteracPotentialVal - Electron-ion interaction potential (Veljkovic et al., 1985)
2. log10RelMutab - Relative mutability (Dayhoff et al., 1978b)
3. ratioAvrgComputedComp - Ratio of average and computed composition (Nakashima et al., 1990)
4. AmphiphilicityIndex - Amphiphilicity index (Mitaku et al., 2002)
5. percent_disorder - Bio.SeqUtils.ProtParam

6. netCharge - Net charge (Klein et al., 1984)
7. FreeESol_inWater - Free energy of solution in water, kcal/mole (Charton-Charton, 1982)
8. percentage_amino_acids_in_Sheet - Bio.SeqUtils.ProtParam
9. AvrgVolResidues - Average volumes of residues (Pontius et al., 1996)
10. AlphaHelix_Propensity - Alpha-helix propensity derived from designed sequences (Koehl-Levitt, 1999)
11. pI - Isoelectric point (Zimmerman et al., 1968)
12. SideChainContrib_ProtStab - Side-chain contribution to protein stability (kJ/mol) (Takano-Yutani, 2001)
13. percentage_amino_acids_in_Turn - Bio.SeqUtils.ProtParam
14. NormFreqAlphaHelixUnweight - Normalised frequency of alpha-helix (Maxfield-Scheraga, 1976)
15. PhyloCSF - Phylogenetic Codon Substitution Frequencies
16. Percentage_basic_amino_acids - Bio.SeqUtils.ProtParam
17. PositiveCharge - Positive charge (Fauchere et al., 1988)
18. aa_length - length of ncORFs expressed as the corresponding number of amino acids
19. RatioAccesBuriedMolfrac - Ratio of buried and accessible molar fractions (Janin, 1979)
20. PercExposedResidues - Percentage of exposed residues (Janin et al., 1978)
21. RelStabilScale - The relative stability scale extracted from mutation experiments (Zhou-Zhou, 2004)
22. NEIGIndex - NNEIG index (Cornette et al., 1987)
23. NormFreqAlphaHelix - Normalised frequency of alpha-helix (Maxfield-Scheraga, 1976)
24. MeanPolarity- Mean polarity (Radzicka-Wolfenden, 1988)
25. Flexibility - Bio.SeqUtils.ProtParam
26. 8AcontactNumber - 8 A contact number (Nishikawa-Ooi, 1980)
27. AliphaticIndex - Bio.SeqUtils.ProtParam
28. NormFreqBetaSheetUnweight - Normalised frequency of beta-sheet, unweighted (Levitt, 1978)
29. SolubilityIndex - Bio.SeqUtils.ProtParam
30. MolWeight_div_aaLength - ratio Molecular Weight (Molecular weight Fasman, 1976) to the corresponding length in number of amino acids
31. SignalSeqHelicalPotential - Signal sequence helical potential (Argos et al., 1982)
32. Avrg_FlexibilityIndices - Average flexibility indices (Bhaskaran-Ponnuswamy, 1988)
33. StabilityScale - The stability scale from the knowledge-based atom-atom potential (Zhou-Zhou, 2004)
34. percentage_amino_acids_in_Helix - Bio.SeqUtils.ProtParam
35. n_predHLAbinders_div_aaLength - ratio of number of predicted HLA-binder peptides (this study) to the corresponding length expressed in number of amino acids

36. InstabilityIndex - Bio.SeqUtils.ProtParam
37. Orf_biotype - uoORF, upstream overlapped open reading frame
38. Orf_biotype - uORF,  upstream open reading frame
39. Orf_biotype - intORF,  internal open reading frame
40. Orf_biotype - lncRNA, long non-coding RNA
41. Orf_biotype - dORF, downstream open reading frame
42. Orf_biotype - processed transcript
43. Orf_biotype - doORF, downstream overlapped open reading frame

As the orf_biotype values represented categorical values and numeric values are required to generate and test the statistical model, 1 (as a value) was assigned to each ncORF with the corresponding orf_biotype category, and 0 when this did not apply. The ncORF Molecular Weight and the number of predicted HLA-binding peptide values were divided by the corresponding amino acid length.

Given that 1785 ncORFs were detected while 5479 remain undetected, presenting an approximate ratio of 1:3, the dataset exhibits an inherent imbalance. Therefore, we employed a balanced weight for imbalanced classification in Keras to address the imbalance, and a neural network analysis to build, train, and evaluate a TensorFlow-Keras model (1). The dataset with the selected attributes was used to implement this model using Python 3, and pandas, numpy, matplotlib, and various components of TensorFlow and Keras for building and evaluating the model. Training and testing sets were separated from the target variable, allocating 80% for training and 20% for testing, using the train_test_split function from scikit-learn.

Before fitting the model, the features were standardised using the StandardScaler, a preprocessing step to verify the features are on the same scale. The model was built as a sequential model consisting of multiple layers: the input layer with 16 neurons, ReLU activation, and L2 regularisation. Batch normalisation and dropout layers were added after each hidden layer to prevent overfitting. The output layer consisted of a single neuron with sigmoid activation for binary classification.

The model was compiled using the Adam optimiser with a learning rate of 1e-3, binary cross-entropy loss function, and accuracy as the metric. It was trained on the training data. The training was run for 60 epochs with a batch size of 12. The model was used to predict the target variable for the test set, and the predictions were used to calculate the Receiving Operating Characteristic Area Under the Curve, ROC AUC score and make binary predictions.

Evaluation of the model:

**ROC AUC Score: 0.678**
**Confusion Matrix: [[620 454]**
**[119 260]]**
**Classification Report:**

| | precision | recall | f1-score | support |
|---|---|---|---|---|
| **0** | **0.84** | **0.58** | **0.68** | **1074** |
| **1** | **0.36** | **0.69** | **0.48** | **379** |
| **accuracy** | | | **0.61** | **1453** |
| **macro avg** | **0.60** | **0.63** | **0.58** | **1453** |
| **weighted avg** | **0.72** | **0.61** | **0.63** | **1453** |

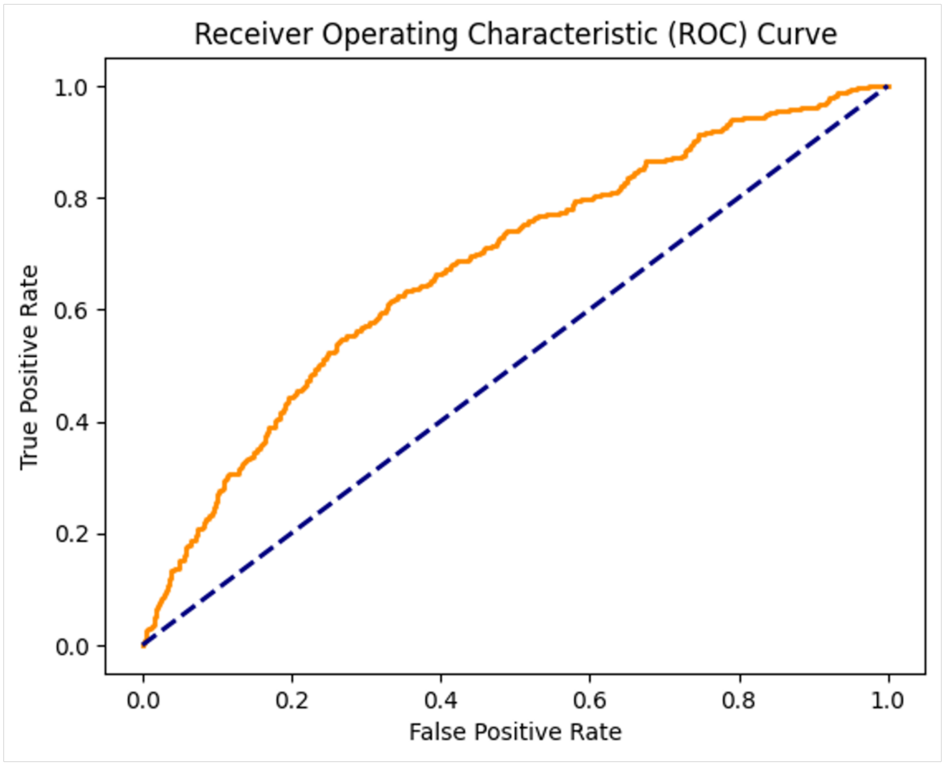


**Figure 2**. Receiving Operating Characteristic (ROC) Curve.

**Figure 2** shows the results of the model on the test set as a ROC plot. The ROC AUC score is an indicator of how well the Tensorflow-Keras binary classification model discriminated between detected and undetected ncORF peptides. A value of **0.68** indicates that the model exhibits a moderate capacity to distinguish between the detected and undetected open ORFs. The ROC AUC curve (orange) specifically shows the relationship between the true positive rate (sensitivity) and the false positive rate (1-specificity) for this model.

Considering the confusion matrix results, the model made 260 correct predictions for the negative class (undetected) and 620 correct predictions for the positive class. However, it also misclassified 119 negative instances as positive and 454 positive instances as negative.

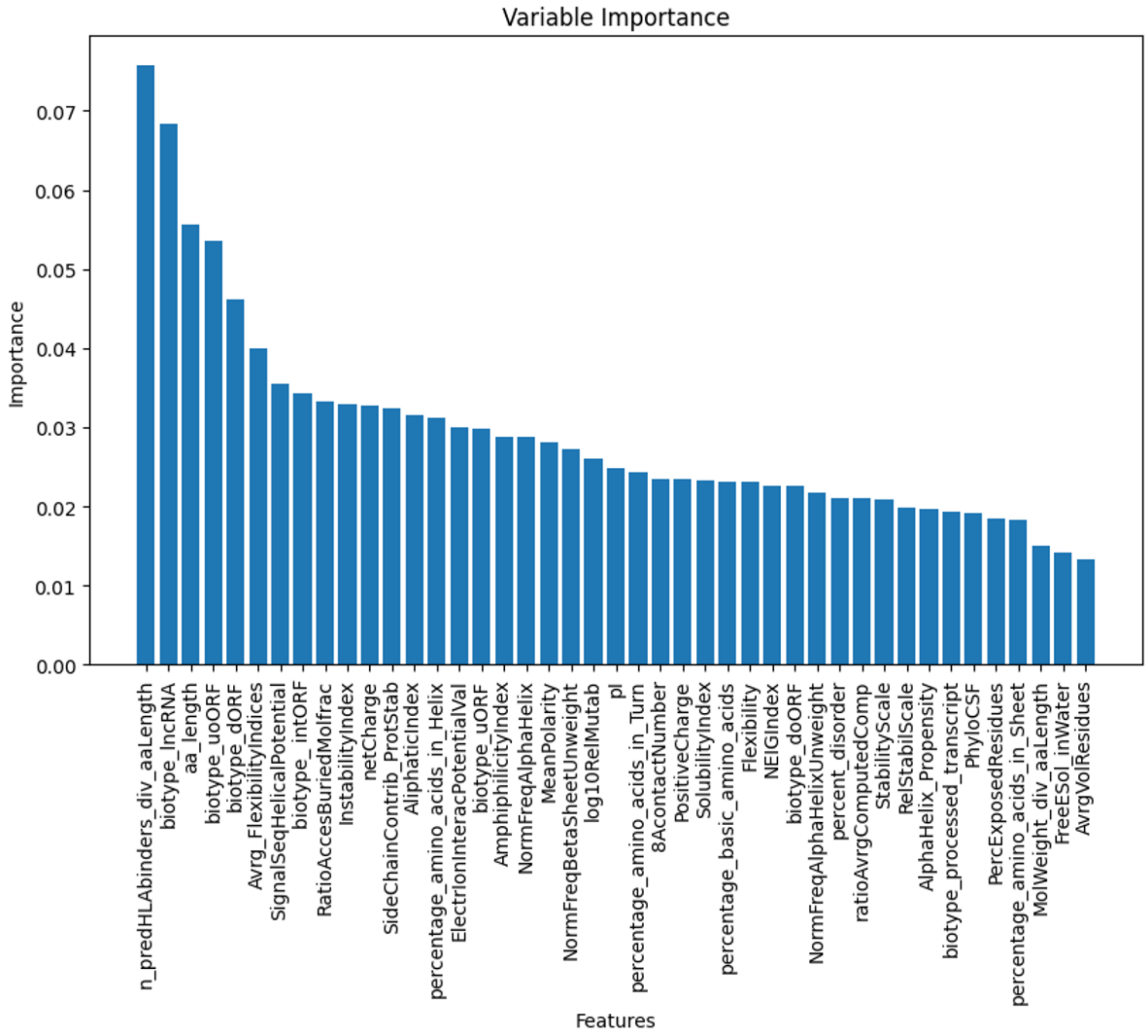


**Figure 3.** Variable Importance scores bar plot.

**Supplementary Table S8** provides a complete list of all 7264 ncORFs, including their identifier, sequence, the 43 features that the model used for training, and the output probabilities from the TensorFlow-Keras model.

The ratio of number of predicted HLA-binding peptides to ORF length (in amino acids) emerged as the highest importance attribute. The frequency of ORF detectability along this ratio and the proportions of detected and undetected are shown in **Figure 4**. Histogram distributions for detected and undetected ORFs appeared fairly similar, yet the model profits from these small differences. Besides, more than 94% of peptides (2,937 out of 3,116; 94.3%) matching ncORFs were found to be presented by HLA.

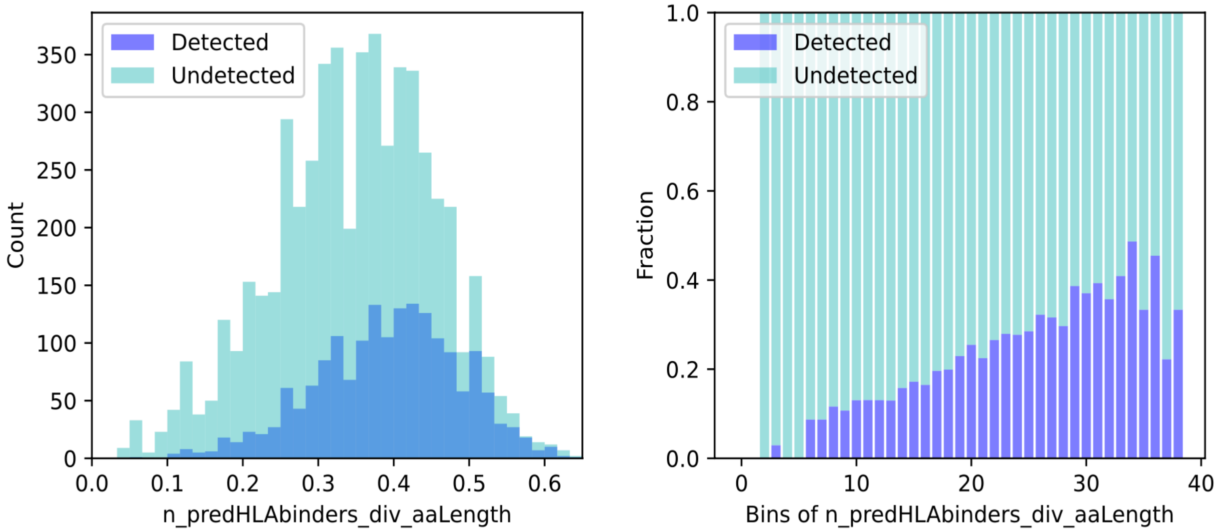


**Figure 4.** Ratio of the Number of predicted HLA-binding peptides to the ORF amino acid length's frequency represented as a Histogram plot (left side) and the corresponding Proportions plot (right side).

The ORF Biotype, specifically 'lncRNA', becomes the second more important variable. **Figure 5** represents the number of ORF lncRNA microproteins with all the designated ORF biotypes along the ORF detectability (left side) and the corresponding proportions on the right side.

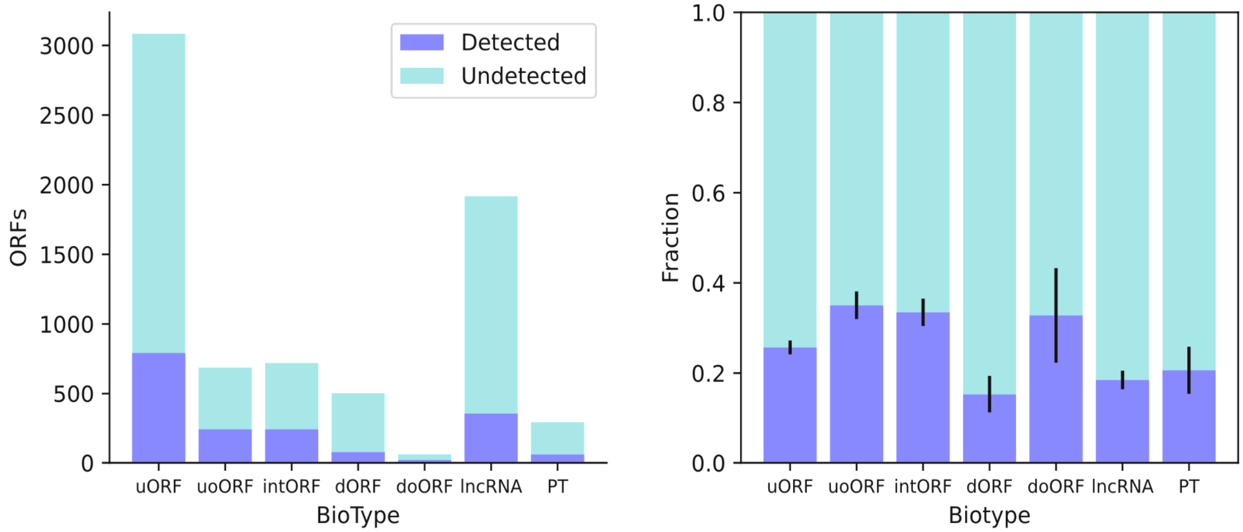


**Figure 5.** ORF Biotype counts versus Biotype. A bar plot (left side) representing the designated ORF biotypes along with ORF detectability (left side), and the corresponding Proportions plot (right side).

The ORF length expressed as the number of amino acids emerged as the third variable of importance. **Figure 6** represents the frequency of the detected or undetected ORFs along their amino acid length (left side) and the corresponding proportions (right side).

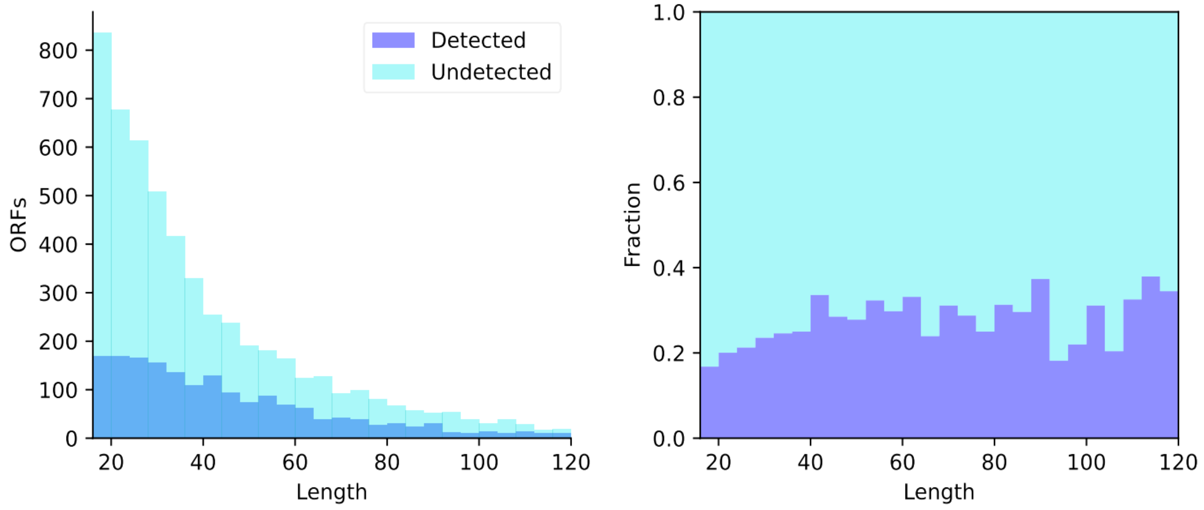


**Figure 6.** ORF amino acid length. The frequency of ORF detectability and the corresponding ORF length in number of amino acids are illustrated as a histogram plot (left side) together with the Proportions plot (right side).

While the model showed moderate discriminatory ability (as indicated by the ROC AUC score of 0.68), a substantial number of instances were misclassified, as revealed by the confusion matrix, thus there are likely additional factors that influence the detectability of ncORF.

The Number of Predicted HLA-binding peptides to length (amino acids) ratio appeared to be the most important attribute. Unsurprisingly, ncORFs rich in predicted HLA-binding peptides should be more easily detected.

The designated biotype of the ORF, specifically whether it originates from a lncRNA, emerged as the second most important attribute for their potential detectability. ORF microproteins derived from lncRNAs and dORFs are a little less likely to be detected, while the uORFs, uoORFs, and intORFs are moderately more likely to be detected.

Overall, the ncORF length affects detectability, as the shortest ncORFs are more difficult to detect.

#### 2. ncORF HLA-I predicted binder peptides classification model

Considering the model results in section 1, where the number of predicted HLA-binding peptides may play an influential role in the ncORF detectability, and most MS-based proteomics approaches identify peptides, the next step was to focus on the analysis of ncORF peptide sequences. Then, to discern potential differences between detected and undetected HLA-I predicted binder peptides, we curated a dataset comprising both categories and applied a Statistical Learning model for analysis.

We focused on 9-amino acid (9aa) long detected HLA-I predicted peptides, originating from ncORFs. We created a list of 341 9aa detected peptides, each with a binding score (min_rank) falling within the range of 0.1 to 1.7. We then paired these peptides, along with their corresponding best alleles, with undetected HLA-I predicted peptides of the same length (9aa), ensuring alignment with the closest possible binding score and allele.

From the combined set of all peptides (totaling 677), we derived a comprehensive array of 63 attributes. These attributes were computed utilising both the Bio.SeqUtils.ProtParam module from BioPython and the Amino Acid Indices version 9.2 (https://www.genome.jp/). Employing the Boruta algorithm (https://gitlab.com/mbq/Boruta/), we selected the most discriminative features from the pool of 63 attributes.

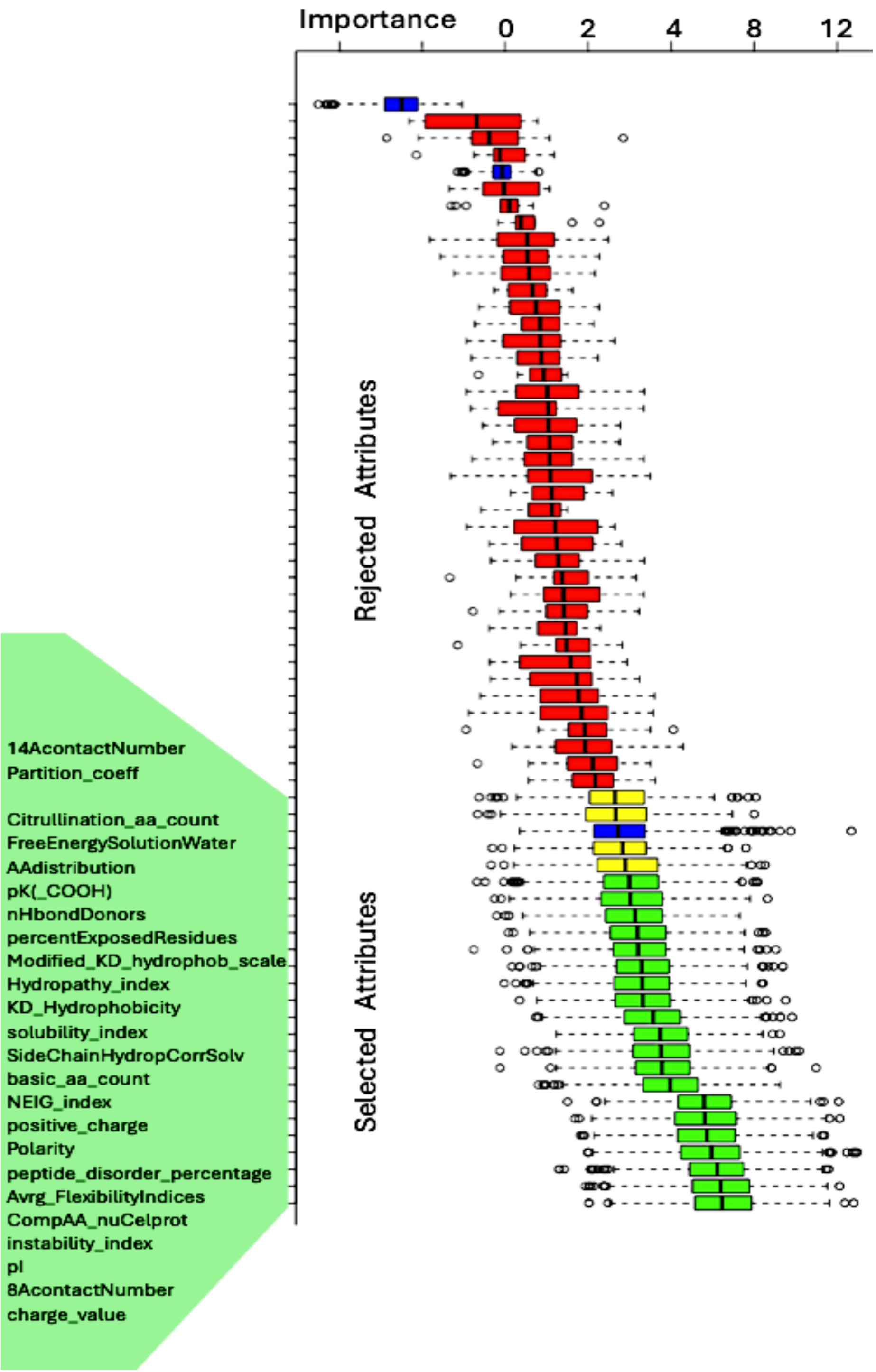


**Figure 7.** Boruta selected attributes. As described in the previous section, features that consistently have higher Z-scores than the shadow features are considered important, while those with lower Z-scores are considered unimportant and are removed from the model. Blue box plots correspond to minimal, average and maximum Z score of a shadow attribute. Red and green box plots represent Z scores of respectively rejected and confirmed, and yellow boxplots are considered tentative attributes.

Boruta selected and confirmed 20 attributes (**Figure 7**, green boxplots), 4 were assessed as tentative (yellow boxplots), all 24 are shown on the left side green panel. 39 attributes were rejected (red boxplots) as being not useful in discriminating between detected and undetected. We further selected all confirmed 20 (green boxplots) and added 2 tentative attributes. The following 22 attributes associated with these ncORF peptides were finally utilised to test a statistical learning Multilayer Perceptron (2) model, (MLPClassifier):

1. Partition_coeff - Partition coefficient (Garel et al., 1973)
2. Citrullination_aa_count - Number of residues that can be modified by Citrullation
3. FreeEnergySolutionWater - Free energy of solution in water, kcal/mole (Charton-Charton, 1982)
4. pK(_COOH) - pK-a(RCOOH) (Fauchere et al., 1988)
5. nHbondDonors - Number of hydrogen bond donors (Fauchere et al., 1988)
6. percentExposedResidues - Percentage of exposed residues (Janin et al., 1978)
7. Modified_KD_hydrophob_scale - Modified Kyte-Doolittle hydrophobicity scale (Juretic et al., 1998)
8. Hydrophathy_index - Hydropathy index (Kyte-Doolittle, 1982)
9. KD_Hydrophobicity - Hydrophobicity (Kyte & Doolittle, 1982)
10. solubility_index - Bio.SeqUtils.ProtParam
11. SideChainHydropathyCorrectSolvation - Side chain hydropathy, corrected for solvation (Roseman, 1988)
12. basic_aa_count - number of basic amino acid residues.
13. NEIG_index - NNEIG index (Cornette et al., 1987)
14. positive_charge - Positive charge (Fauchere et al., 1988)
15. Polarity - Polarity (Grantham, 1974)
16. peptide_disorder_percentage - Bio.SeqUtils.ProtParam
17. Avrg_FlexibilityIndices - Average flexibility indices (Bhaskaran-Ponnuswamy, 1988)
18. CompAA_nuCelprot - Composition of amino acids in nuclear proteins (percent) (Cedano et al., 1997)
19. Instability_index - Bio.SeqUtils.ProtParam
20. pI - Isoelectric point (Zimmerman et al., 1968)
21. 8AcontactNumber - 8 A contact number (Nishikawa-Ooi, 1980)
22. Charge_value - Bio.SeqUtils.ProtParam

A dataset with these attributes served as a basis to generate an MLP Classifier model using Python 3, and software/libraries: pandas, numpy, matplotlib, sklearn. This involved data preparation, model initialisation and tuning, model fitting, prediction, and evaluation. First, the features of the dataset were separated from the target variable, and the data was split into training and testing sets, allocating 80% for training and 20% for testing, while ensuring reproducibility by setting the random state to 42.

Before fitting the model, the features were standardised using the StandardScaler, a preprocessing step that removes the mean and scales the features to have unit variance. Next, an MLP Classifier model was initialised with a maximum of 8000 iterations and a random state of 42. The model was then tuned using grid search with cross-validation, exploring various hyperparameters including the hidden layer sizes, activation function, and regularisation parameter. Grid search with cross-validation tuned the model using the specified hyperparameters: Hidden layer sizes: (280,) Activation function: 'tanh', and Regularization parameter (alpha): 0.01.

Upon completion of the tuning process, the best performing model was identified based on the grid search results. This model was used to make predictions on the previously untouched test set, allowing for an assessment of its predictive capabilities. Finally, a range of evaluation metrics including ROC AUC, F1 score, accuracy, precision, recall, and the confusion matrix were calculated to gauge the model's performance on the test set.

Evaluation of the model:

ROC AUC Score: 0.694
F1 Score: 0.732
Accuracy Score: 0.699
Precision Score: 0.737
Recall Score: 0.727
Confusion Matrix: [[39 20]
[21 56]]

Considering all the metrics results, the model demonstrates relatively balanced performance. The accuracy score of 0.699 indicates that approximately 70% of the predictions were correct. The precision and recall scores, both around 0.73, suggest that the model is reasonably good at correctly identifying positive cases and minimising false positives.

Looking at the confusion matrix, the model made 39 correct predictions for the positive class (detected) and 56 correct predictions for the negative class. However, it also misclassified 21 negative instances as positive and 20 positive instances as negative.

**Figure 8** shows a ROC plot for the test set of 677 nc ORF peptides.

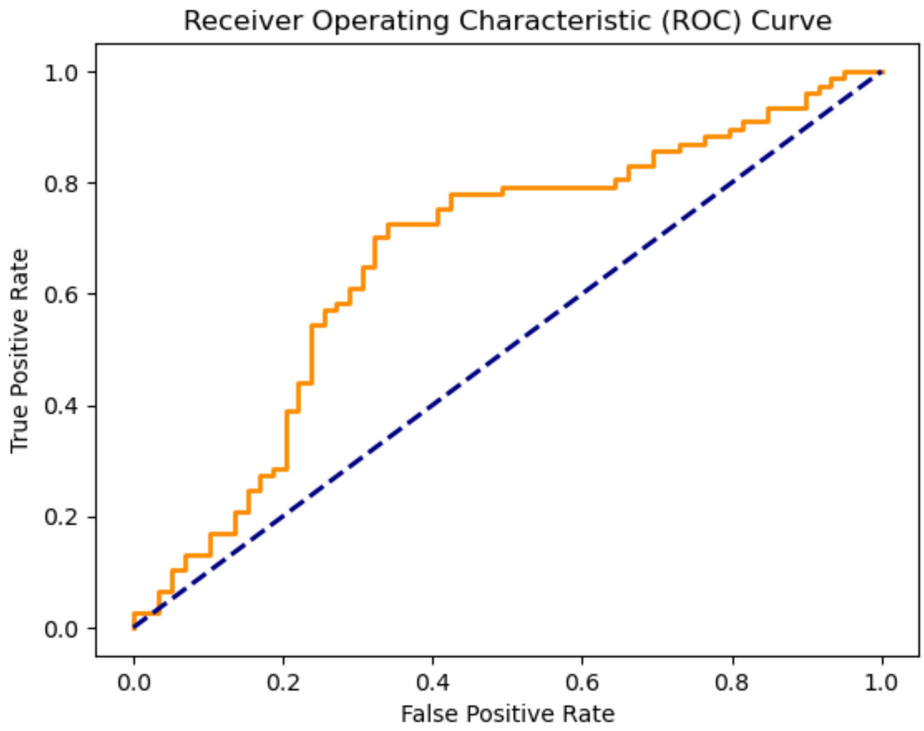


**Figure 9**. MLPClassifier's output ROC AUC curve. The ROC AUC score is an indicator of how well the MLP Classifier model distinguishes between detected and undetected ncORF peptides. A value of **0.69** suggests that the model has some ability to differentiate between the classes. The ROC AUC curve (orange) specifically shows the relationship between the true positive rate (sensitivity) and the false positive rate (1-specificity) for the MLP binary classification model. Here, the model's performance changes as the discrimination threshold is adjusted, providing insights into its ability to correctly classify positive and negative instances. Then, the AUC value quantifies the overall predictive accuracy of the MLP model, with a higher AUC indicating better performance in distinguishing between detected and undetected peptides.

**Supplementary Table S7** provides a complete list of all 677 peptides, including their sequence, best allele, the 22 features that the model used for training, and the output probabilities from the model.

**Figure 10** shows the highest importance computed property, Instability Index versus the model-predicted probability of detection. Instability index is usually calculated using various algorithms that consider factors such as amino acid composition, secondary structure, and other physicochemical properties. **Figure 10** suggests an increase in ncORF HLA-I peptide detectability correlates with an increase in stability.

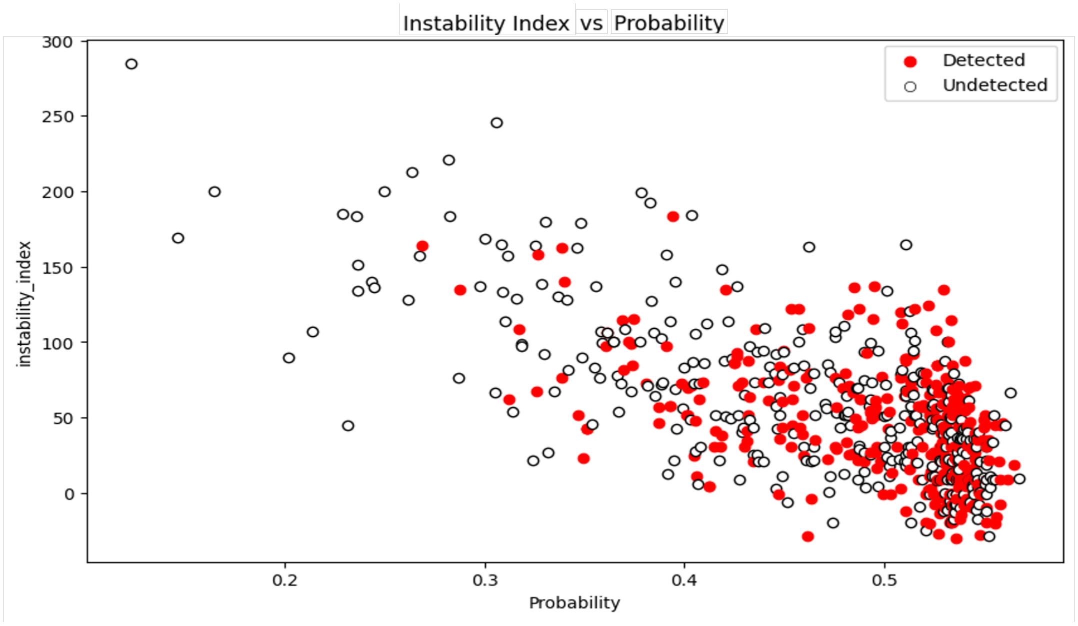


**Figure 10.** Scatter plot of the probability of detection for each ncORF HLA-I predicted peptide based on the Instability Index. The detected ORF peptides are depicted in red.

**Figure 4** shows the Variable Importance scores for the attributes associated with the Instability Index according to the MLP model.

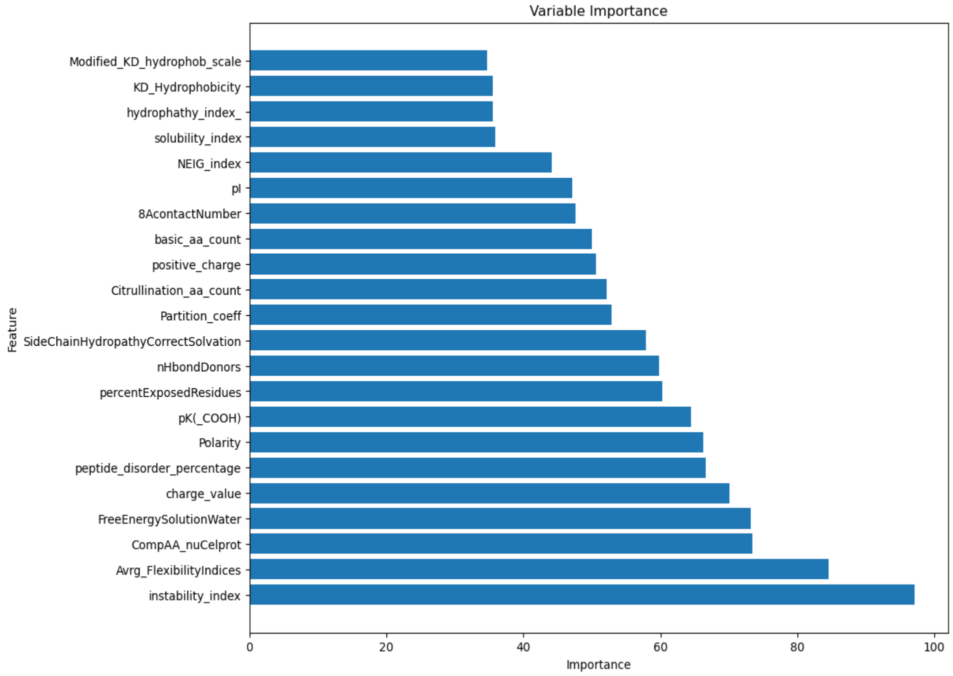


**Figure 11.** Variable Importance scores bar plot.

While the model showed moderate discriminatory ability (as indicated by the ROC AUC score) and achieved decent accuracy, precision, and recall scores, it also misclassified a notable number of instances. The model performance may improve e.g. when data on ncORFs HLA-I peptide abundance or other relevant attributes become available.

Although the performance of the model is not optimal to classify peptides as detectable or undetectable, the most important features appear to be correlated, with the peptide Instability Index as the most important attribute for their potential detectability.

A lower instability index suggests that a peptide is more stable, whereas a higher index indicates greater instability. The usefulness of this variable in the model can be explained by several key factors:

Peptide Degradation: peptides with high instability indices are more prone to degradation. In MS-based experiments, unstable peptides may degrade during sample preparation, storage, or analysis, leading to reduced detectability. Stable peptides are more likely to remain intact, ensuring they reach the mass spectrometer and produce reliable signals.

Ionisation Efficiency: the stability of a peptide can affect its ionisation efficiency. Unstable peptides might undergo partial degradation or modifications that alter their ionizable groups, impacting their ionisation efficiency and, consequently, their detectability by MS.

Fragmentation Patterns: in MS/MS (tandem MS), peptides are fragmented to provide sequence information. Peptides with a high instability index might produce unpredictable or incomplete fragmentation patterns, complicating their identification and reducing confidence in their detection.

Protein Expression and Processing: the stability of peptides can also reflect their processing and expression within the cell. Stable peptides are likely to be present in higher quantities and more consistently processed, leading to higher chances of detection.

Biological Relevance: The Instability Index might correlate with other biologically relevant properties such as protein half-life, subcellular localisation, and interaction with other cellular components. These factors collectively influence the overall abundance and accessibility of peptides for MS analysis.

1. Martín Abadi et al. TensorFlow: Large-scale machine learning on heterogeneous systems, 2015.
2. Cybenko, G. Approximation by superpositions of a sigmoidal function Mathematics of Control, Signals, and Systems, 1989. 2(4), 303–314.
